# White matter microstructure fingerprint of cerebral small vessel disease

**DOI:** 10.1101/2025.10.01.679737

**Authors:** Aurélie Bussy, Camille Cathala, Fábio Carneiro, Ferath Kherif, Antoine Lutti, Bogdan Draganski

**Affiliations:** Imaging Neuroscience of Ageing laboratory inAGE, Universitätsklinik für Neurologie, Inselspital Bern, University of Bern, Switzerland; Graduate School for Health Sciences, University of Bern, Bern, Switzerland; Institute for diagnostic and interventional neuroradiology, Inselspital Bern, University of Bern, Switzerland; Laboratory for Research in Neuroimaging LREN, Centre for Research in Neurosciences, Department of Clinical Neurosciences, Lausanne University Hospital and University of Lausanne, Lausanne, Switzerland; Signal Processing Laboratory 5, Ecole Polytechnique Fédérale de Lausanne, Switzerland

**Keywords:** White matter hyperintensities, normal-appearing white matter, microstructural imaging, cardiovascular risk factors

## Abstract

**Background:** White matter hyperintensities (WMHs) are the imaging hallmark of cerebral small vessel disease (SVD), yet their microstructural composition, spatial heterogeneity, and relationship to diffuse normal-appearing white matter (NAWM) damage and cardiovascular risk remain incompletely defined.

**Methods:** In 363 community-dwelling adults from the BrainLaus cohort (mean age 55.5 years; range 19.8–89.4; 48.8% male), we combined quantitative relaxometry (MTsat, R1, R2*) and diffusion-derived metrics (FA, MD, NODDI, g-ratio). WMHs were automatically segmented on FLAIR and microstructure was quantified across lobes, white matter compartments, and geodesic layers extending from the WMH core into surrounding NAWM. Multivariate organisation was assessed using principal component analysis, and associations with cardiovascular risk factors were tested using partial least squares.

**Results:** Our analysis revealed demyelination, axonal loss, and extracellular fluid accumulation, particularly in periventricular regions. Layer-specific profiles showed a centrifugal gradient, with myelin loss and oedema at the core and axonal alteration in surrounding tissue. These signatures were associated with age-related cardiovascular risk factors, including higher blood pressure, bioimpedance, and lower hemoglobin levels.

**Conclusion:** WMHs index the endpoint of a broader, spatially structured white matter injury process that extends into NAWM, is regionally concentrated in periventricular tissue, and covaries with systemic vascular/metabolic factors. These findings support sustained vascular risk management to mitigate CSVD-related white matter degeneration.

## Introduction

White matter hyperintensities (WMH) are radiological features commonly observed in conventional magnetic resonance images such as Fluid Attenuated Inversion Recovery (FLAIR), in older adults. They are considered a hallmark of cerebral small vessel diseases (SVD), most often reflecting vascular injury such as arteriolosclerosis or cerebral amyloid angiopathy (CAA). WMHs are frequent incidental findings and are associated with increased risk for gait impairment, stroke, and cognitive decline.^1,2^ Neuropathology studies indicate that WMHs reflect tissue damage due to demyelination, axonal loss, and gliosis,^3,4^ yet their biological mechanisms and temporal dynamics remain poorly defined. Although consistently associated with cardiovascular risk factors (CVR) and systemic inflammation,^5–8^ the spatial progression and microstructural consequences of WMHs in relation to these systemic factors remain largely underinvestigated.

Emerging evidence challenges the traditional view of WHM as discrete lesions. Rather, they suggest that WMHs represent the macroscale imaging signature of a widespread white matter (WM) damage that also affects the surrounding normal-appearing white matter (NAWM).^9^ This broader pathological process appears to follow a centrifugal gradient of tissue damage extending outward from the core lesion, potentially reflecting secondary neuroinflammatory response, axonal degeneration, or impaired remyelination. To support this gradient model, studies have demonstrated that the area surrounding WMHs is characterised by changes in diffusion tensor metrics indicative of microstructural damage, defined as WMH penumbra.^10,11^ The existence of lesion-centred gradients of microstructural pathology in the WMH penumbra is well-established ^10,11^, yet prior characterisation has predominantly relied on single or only a few magnetic resonance imaging contrasts and limited spatial frameworks, leaving the microstructural composition of this continuum incompletely resolved.

Regional variation in WMH and NAWM tissue microstructural properties in relation to ventricular or cortical proximity^12^ and the antero-posterior axis^13–15^ has likewise been described, but the comprehensive microstructural characterisation in the spatial context of these patterns and WMH load is absent from the literature.

In the radiology diagnostic setting, FLAIR imaging remains the standard for WMH detection and progression tracking.^16^ However, these findings can be difficult to interpret, as a range of pathological processes, including edema, inflammation, and demyelination lead to the same signal abnormalities.^17^ Quantitative MRI partly alleviates this limitation.^18,19^ Relaxometry metrics provide information on tissue composition: magnetization transfer saturation (MTsat) and longitudinal relaxation rate (R1 = 1/T1) are sensitive to myelin content,^20–22^ while the transverse relaxation rate (R2* = 1/T2*) primarily reflects iron accumulation but is also modulated by myelin.^23–25^ The fractional anisotropy (FA) index computed from diffusion MRI data reflects fiber coherence, mean diffusivity (MD) reflects overall water mobility, intracellular volume fraction (ICVF) captures axonal/neurite density, isotropic volume fraction (ISOVF) reflects extracellular free water, and orientation dispersion (OD) indexes neurite geometry and angular variability.^26^

Together, these complementary qMRI metrics provide a multiparametric characterization of brain tissue that goes beyond any single modality. While each parameter is sensitive to a specific biological substrate -such as myelin, iron, axonal density, or extracellular water-, their joint assessment enables a more holistic view of microstructural integrity. This integrative approach is often described as an *in vivo* histology framework ^27^, as it allows researchers to infer cellular- and molecular-level features non-invasively, bridging the gap between conventional MRI contrasts and histopathological validation. Most prior work has shown that WMH are associated with demyelination or axonal loss, however they have focused on isolated brain regions or single imaging metric, limiting our understanding of how lesions and surrounding NAWM interact across the brain.

In this study, we leverage a comprehensive multicontrast MRI dataset from a cohort of older adults to investigate: first, the microstructural properties of WMH and NAWM across distinct WM compartments. Second, using regional, multivariate, and layer-specific analyses, we characterize the pattern, extent, and spatial distribution of WM pathology. Third, we further examine how CVR relate to both lesion load and microstructural variations, aiming to disentangle shared and region-specific patterns of vulnerability. By integrating these dimensions, we provide detailed insights into how specific CVR are associated with microstructural brain tissue properties, informing strategies for targeted prevention of WM degeneration in aging.

## Methods

### Participants

We analysed brain imaging data acquired in the BrainLaus project, a nested study of the epidemiological CoLaus|PsyCoLaus longitudinal cohort, focusing on investigations of CVR and mental health.^28,29^ The imaging dataset included 423 BrainLaus participants with FLAIR images (**Supplementary Figure 1)**. After quality control (QC) of all multi-contrast MRI data, the final sample for further analyses was confined to 363 participants (55.5 ± 22.3, 48.8% males). Of these, 202 participants (69.4 ± 6.9 years; 46.5% male) had complete CVR data and were included in CVR–microstructure analyses (**Table 1**). Age, sex, and Fazekas score distributions at each exclusion step are shown in **Supplementary Figure 2** and **Supplementary Table 1**. QC-based exclusions did not bias age, sex, or Fazekas score distributions, but CVR availability excluded younger participants. Ethical approval was granted by the Ethics Commission of Canton de Vaud, and participants gave their informed consent prior to participation.

#### Cardiovascular risk factors

We included the following CVR: <u>physiological measures:</u> systolic (SBP) and diastolic blood pressure (DBP) in mmHg as the average of the second and third out of three measurements; body mass index (BMI); bio-impedance, providing the percentage of fat mass; waist-to-hip ratio (WHR); <u>blood work-up</u>: HDL cholesterol, LDL cholesterol, total cholesterol, triglycerides, blood glucose, insulin, glycated hemoglobin (HbA1c), high-sensitivity C-reactive protein (CRP), hemoglobin, thyroid-stimulating hormone (TSH), free thyroxine (FT4); <u>lifestyle factors</u>: alcohol consumption (units per week), caffeine intake (nr caffeinated drinks per day) and tobacco consumption (nr cigarette equivalents per day) (for details see **Supplementary figure 3**). Some CVR variables had missing values (**Supplementary figure 4**), but only bioimpedance and coffee consumption exceeded 1% missingness. As no differences in age, sex, SBP, total cholesterol, or WHR were observed between participants with and without missing data (**Supplementary figure 5**), missing values were imputed using variable means. This enabled inclusion of all variables in complete-case analyses (e.g., PLS) while minimizing potential bias. Cross-correlations in the imputed dataset are shown in **Supplementary figure 6**.

### MRI acquisition

All MRI data were acquired from 02/2022 to 07/2024 on a 3T whole-body Siemens Magnetom Prisma system with a 64-channel radiofrequency (RF) receive head coil and a body coil for transmission. No major hardware or software update was conducted over the duration of the study.

### FLAIR protocol

The FLAIR protocol included a repetition time (TR) of 5000 ms, an echo time (TE) of 389 ms and an inversion time (TI) of 1800 ms. Image resolution was 1 mm isotropic. To reduce the acquisition time, a GRAPPA acceleration factor of 2 was used along the phase-encoding direction and Partial Fourier factor of 7/8 was used along the slice direction.

### Relaxometry protocol

The relaxometry data was acquired using a custom-made 3D GRE pulse sequence.^30^ Multi-echo images were acquired with T1- (TR/α = 18.7 ms/20°), PD- and MT-weighted contrasts (23.7 ms/6°) and TE between 2.2 ms and 19.7 ms.^30,31^ Additional acquisition parameters included: 1 mm isotropic resolution, a matrix size of 256 × 240 × 176, parallel imaging with a GRAPPA factor of 2 in the phase-encoding direction, 6/8 partial Fourier in the partition direction, non-selective RF excitation, a readout bandwidth (BW) of 425 Hz/pixel, and an RF spoiling phase increment of 50°, resulting in a total acquisition time of approximately 19 minutes. B1-mapping data was also acquired using the 3D EPI SE/STE method^32,33^ to correct for the effect of radio-frequency field inhomogeneities on the relaxometry data ^22^.

### Diffusion-weighted protocol

For diffusion-weighted imaging (DWI), we used a 2D Echo-Planar Imaging (EPI) sequence with the following parameters: TR = 7400 ms, TE = 69 ms, parallel GRAPPA acceleration factor = 2, FOV = 192 × 212 mm², voxel size = 2 × 2 × 2 mm³, matrix size = 96 × 106, 70 axial slices, and 118 gradient directions (15 at b = 650 s/mm², 30 at b = 1000 s/mm², 60 at b = 2000 s/mm², and 13 at b = 0 interleaved throughout the acquisition).^34^ Additionally, B0-field maps were acquired using a 2D double-echo FLASH sequence with a slice thickness of 2 mm, TR = 1020 ms, TE1/TE2 = 10/12.46 ms, α = 90°, and readout BW = 260 Hz/pixel. These maps were used to correct for geometric distortions in the echo-planar imaging data.

### MRI preprocessing

#### Quantitative maps estimation

Quantitative maps of magnetization transfer saturation (MTsat), the transverse relaxation rate (R2* = 1/T2*), the effective longitudinal relaxation rate (R1 = 1/T1), and the effective proton density (PD*) were computed from the raw MRI data using the VBQ toolbox.^31,35^ The PD* maps were normalized assuming a mean value of 69% in WM.^36^ However, because this normalization leads to a bias of the PD* values in WM, PD* maps were excluded from further analyses (see Supplementary Methods). An example of the included multi-parameter maps are presented along with the other modalities in **Figure 1E**.

**Figure 1:**
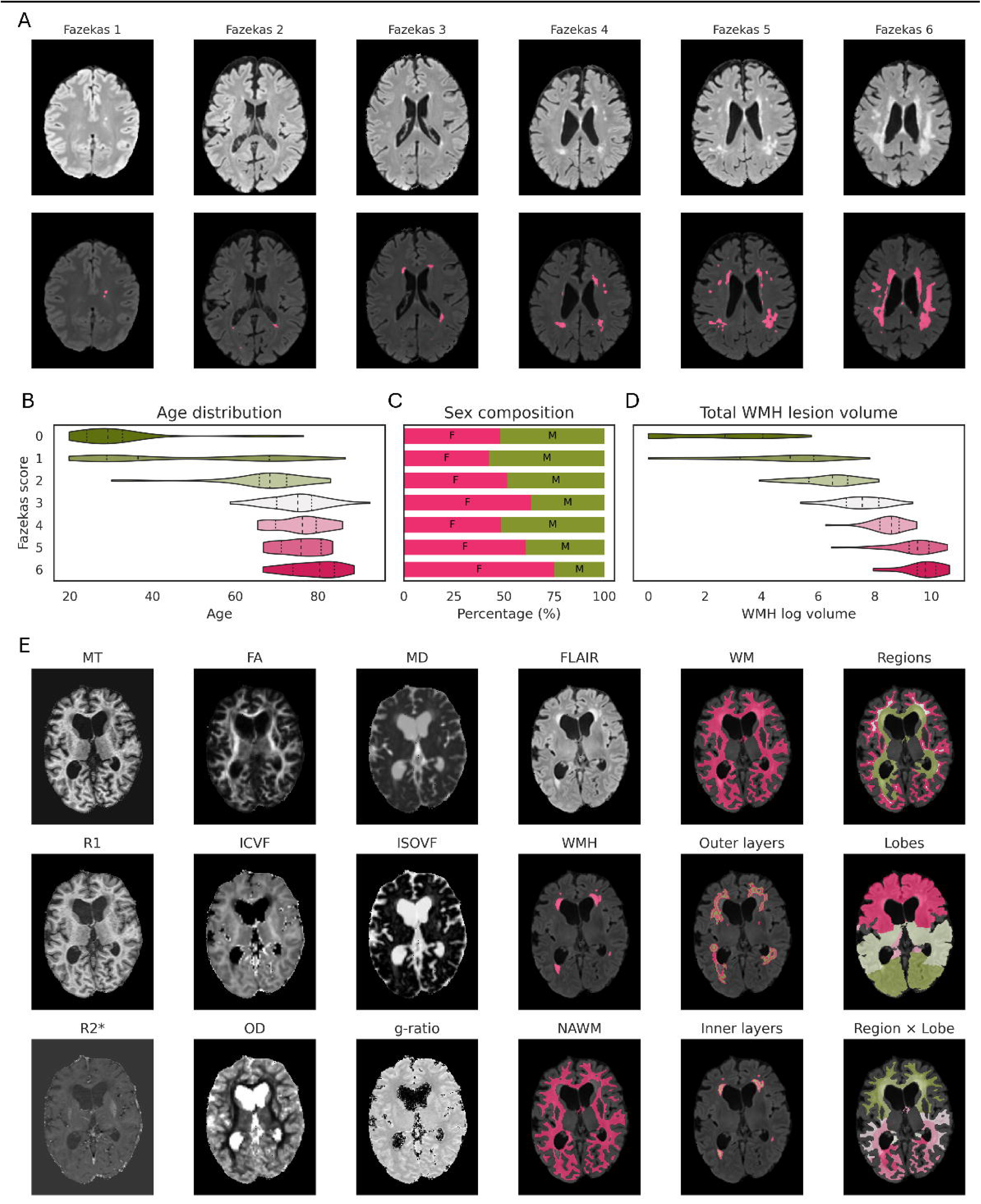
Representative subject data and white matter segmentation framework. **(A)** Examples of brains classified from Fazekas grade 0 (no WMH) to grade 6 (large WMH in deep and periventricular white matter). **(B)** Age distribution stratified by Fazekas grade. **(C)** Sex distribution (percentage male/female) across Fazekas grades. **(D)** Distribution of log-transformed WMH volume within each Fazekas grade. **(E)** Illustrative maps from one participant (81-year-old female) showing quantitative MRI metrics, FLAIR, and tissue masks. The maps include myelin- and iron-sensitive metrics (MTsat, R1, R2*), axon-sensitive metrics (ICVF, OD, FA), and fluid/composite metrics (ISOVF, MD, g-ratio), alongside masks for WM, WMH, NAWM, outer and inner layers, WM regions (SWM, DWM, PVWM), lobes (frontal, parietal, temporal, occipital), and individual WMH lesions. **Abbreviations:** magnetization transfer saturation (MTsat), longitudinal relaxation (R1), transverse relaxation (R2), intracellular volume fraction (ICVF), orientation dispersion (OD), fractional anisotropy (FA); isotropic volume fraction (ISOVF), mean diffusivity (MD), white matter (WM), white matter hyperintensities (WMH), normal-appearing white matter (NAWM), superficial white matter (SWM), deep white matter (DWM), periventricular white matter (PVWM).

#### Diffusion MRI Processing

The DWI data underwent preprocessing using MRtrix3,^37^ which included denoising and removal of Gibbs ringing artifacts.^34^ Eddy current distortions and head movements were corrected with the FSL 5.0 EDDY tool.^38^ Distortions of the EPI images were corrected using the FieldMap toolbox^39^ of SPM12 (Wellcome Trust Centre for Human Neuroimaging, https://www.fil.ion.ucl.ac.uk/spm) and the dedicated B0-field mapping data. The bias field was estimated from the mean b=0 images and corrected in all DWI datasets. Applying the tensor model, we obtained fractional anisotropy (FA) and mean diffusivity (MD) whole-brain maps. Using the neurite orientation dispersion and density imaging (NODDI) model and the multi-shell DWI data,^40^ we calculated maps of intracellular volume fraction (ICVF), isotropic volume fraction (ISOVF) and orientation dispersion index (OD) using the AMICO toolbox.^41^ MRI g-ratio maps, which represent the ratio between the inner and the outer diameter of the myelin sheath were created.^34,42^ All diffusion-derived maps were aligned to the MTsat images using SPM12s rigid-body registration and resampled accordingly to 1×1×1mm resolution. Layer-based sampling was subsequently performed in this common space, ensuring spatial correspondence across contrasts for the 1-mm layer analysis.

#### FLAIR preprocessing

We used SPM12s “unified segmentation” for bias field correction of the FLAIR data^43^ and CAT12 for denoising using a spatial adaptive non-local means denoising filter.^44^ For rigid registration of FLAIR scans to MPM space was performed using FSL FLIRT.^45,46^

#### Quality control

We performed visual QC on all FLAIR, MTsat, R1, R2*, and NODDI maps (see **Supplementary Figure 7** for details). Images rated ≤2.5 were included, while those ≥3 were excluded; participants were removed if any map failed QC. In total, 60 images were excluded, primarily due to motion-related artifacts in R2* and MPM maps (**Supplementary Figure 8**).

#### White matter hyperintensity assessment

FLAIR images were visually rated using the Fazekas scale, which grades periventricular (PVWM) and deep WMH (DWM) from 0 to 3 based on lesion extent.^47^ Scores were summed to yield a total severity score (0-6; **Figure 1A**). Age, sex, and WMH volume distributions across Fazekas scores are provided in **Figure 1B-D**. WMH were automatically segmented on co-registered FLAIR-MTsat images,^48^ and normal appearing WM (NAWM) was defined as WM excluding WMH voxels.

To assess the spatial extent of microstructural changes around WMH, geodesic distance maps were generated using the Fast Marching Method, with values sampled in 0-5 mm shells restricted to NAWM (**Supplementary Figure 9**). First, participant-level layers combined all WMHs to characterise individual spatial gradients from lesion core to surrounding NAWM and relate them to clinical measures. Second, lesion-level layers were extracted to assess spatial microstructural profiles for each lesion.

For detailed characterisation of WMHs spatial distribution, we defined four WM compartments: i. ventricular mask - combined left and right ventricles from FreeSurfer SynthSeg; ii. PVWM mask - WM voxels within 8 mm geodesic distance of the ventricles; iii. superficial WM (SWM) mask - WM voxels within 2 mm geodesic distance of the cortical boundary, excluding PVWM; iv. DWM mask - remaining WM after excluding PVWM and SWM.

Cortical labels from the MICCAI 2012 multi-atlas dataset^49^ were grouped into frontal, parietal, temporal, and occipital lobes and propagated into WM via distance-based nearest-lobe assignment (**Figure 1E**). Participant numbers and demographics by WM region are provided in **Supplementary Table 2** and **Supplementary Figure 10**.

#### Parameter estimates extraction

Regional averages of R1, MTsat, R2*, FA, MD, g-ratio, ISOVF, ICVF, and OD were extracted in native MTsat space across WMH, NAWM, PVWM, SWM, and DWM regions, as well as within layer masks (−1 to −2, +1 to +5), lobes, and their intersections. Analyses were performed in native space to avoid interpolation and smoothing effects from MNI registration, preserving lesion boundaries and quantitative MRI fidelity.

### Statistical analyses

#### Methodology

Our statistical analyses follow six complementary analyses, to address our main hypotheses. First, we characterized *“Lobar and regional tissue microstructural fingerprints”* to establish fundamental WMH-NAWM differences. We then investigated *“associations between WMH load and microstructural signatures”* to understand volume-damage relationships. To identify dominant patterns, we examined *“regional WM microstructure”* using principal component analysis (PCA). Next, we mapped *“layer-based microstructural profiles”* to characterize spatial gradients around WMH lesions in relation to age. Then we investigated *“microstructural gradients vs WMH volume”*. Finally, we tested *“associations with cardiovascular risk”* to connect imaging findings to systemic health factors using multivariate methods.

#### Technical implementation

All statistical analyses were performed in Python 3.8.19 using Jupyter Notebook, including the layer masks creation, metric sampling, and multivariate analyses.

### Lobar and regional tissue microstructural fingerprints

Within-subject WMH-NAWM differences were quantified using Cohen’s dz, calculated from paired raw values. Statistical significance was assessed with paired Wilcoxon signed-rank tests, with Benjamini–Hochberg FDR correction across metric×region comparisons. Regional and lobar effects were evaluated using Kruskal-Wallis tests.

For visualization, metrics were concatenated across regions and tissue classes and z-scored across all participants and regions, without outlier removal or within-region/tissue standardization. Additional analyses using coarser WM and lobar parcellations replicated the findings (**Supplementary Figure 11**-**12**).

### Associations between WMH load and microstructural signatures

Linear associations between MRI metrics and regional WMH volumes were tested using ordinary least squares regression, which provides a parsimonious first-order approximation of monotonic relationships and allows direct interpretation of effect sizes. Within each region-specific analysis subset, microstructural metrics were z-scored after exclusion of missing values and WMH volumes were log-transformed. The obtained standardized beta coefficients enabled the direct comparison of WMH volume effects across MRI metrics. Analyses were conducted on per-participant mean values within WMH and NAWM masks per participant, limiting the impact of extreme value voxels.

### Regional WM microstructure

We used PCA (Scikit-learn v1.3.2) to identify microstructural patterns. The input matrix included one column per metric, containing regional means across participants, treating NAWM and WMH as separate tissue types. Each participant contributed one value per region per tissue type for each metric. Metrics were z-scored across the full region-by-participant matrix, without within-region or within-tissue normalization to ensure equal weighting. After excluding missing data (e.g., regions without WMHs in a given participant), the matrix comprised 1,849 WMH and 4,158 NAWM observations, reflecting the more variable spatial distribution of WMH. Robustness was evaluated via separate tissue-type analyses (**Supplementary Figures 13-14**), subject-level PCA (**Supplementary Figure 15**), and bootstrap resampling, permutation testing, and split-half replication (**Supplementary Figure 16**).

### Layer-based microstructural profiles

We selected third-order polynomial models after comparing fits across model orders; higher-order models did not provide meaningful improvement (**Supplementary Figures 17, 18 and Supplementary Table 3**). Age effects across WMH layers were assessed using mixed-effects models including age, layer, their interaction, and log-transformed WMH volume, with random intercepts and age slopes per participant. All continuous variables were standardized across subjects to ensure that effect sizes were directly comparable across layers and metrics. The outermost layer served as the reference, and age effects for other layers were derived from the age × layer interactions. Standardized slopes quantified relative age sensitivity across metrics and layers (**Figure 5**).

### Microstructural gradients vs WMH volume

To characterise how the tissue microstructure gradients vary within individual WMHs, we analysed each lesion’s microstructural profile using linear mixed-effects models. Raw metric values were predicted as a function of geodesic distance (modelled as a cubic polynomial), lesion volume (log-transformed), and their interaction, with random intercepts for both participant and lesion. This per-lesion modelling framework allowed us to quantify how the WMH-to-NAWM gradient evolved with distance, to determine whether the gradient’s shape depends on lesion size, and to generate lesion-centred depth profiles that are directly comparable across lesions of different volumes (**Figure 6**).

### Associations with cardiovascular risk

To test for multivariate associations between WM microstructure and CVR or medication, we used partial least squares (PLS) analysis (https://github.com/rmarkello/pyls; ^50^). The brain input matrix contained one column per metric for WMH (9 columns) and one for NAWM (9 columns), each representing the participant-level mean value. This design allowed us to differentiate associations between CVR and microstructure in WMH versus NAWM. Here, we used PLS as an exploratory method to identify latent covariance patterns between MRI-derived tissue microstructural metrics and CVR,^51^ not to construct a predictive model. The CVR matrix included one column per risk factor. The few missing MRI metrics values (1 to 4 entries across 15 columns) were imputed using column-based mean replacement prior to standardization. All variables were then z-scored across participants before entering the PLS analysis. This approach decomposes the brain and CVR matrices into latent variables (LVs) that capture shared variance across both domains. The significance of each LV was tested via permutation, and variable contributions were assessed using bootstrap resampling, following established protocols.^50,52^ Split-half resampling analyses (500 iterations) were conducted to evaluate the stability of the latent variable structure (**Supplementary Figure 19**).

### Multiple comparisons correction

All statistical tests were corrected for multiple comparisons using the Benjamini–Hochberg false discovery rate (FDR). Corrections were applied separately within each analytical family (e.g., region × metric tests, layer × metric tests, PLS resampling tests), ensuring appropriate control of false positives within each specific set of analyses.

## Results

### Lobar and regional microstructural fingerprints of WMH versus NAWM

We observed MRI parameter differences between WMH and NAWM across almost all metrics and regions (**Figure 2AB**). When we compared WMH vs NAWM overall, we found lower values of MTsat (*d* = −3.28), R1 (*d* = −3.64), R2* (*d* = −3.28), ICVF (*d* = −3.14), FA (*d* = −1.47), and OD (*d* = −1.10); in WMH compared to NAWM (**Supplementary Figure 20**). We found higher values of MD (*d* = 3.16) and ISOVF (*d* = 1.63) in WMH compared to NAWM, and small differences of the MRI g-ratio (*d* = −0.43). Across most metrics, WMH showed the expected directionality relative to NAWM: MT, R1, R2*, and ICVF were generally lower in WMH, whereas ISOVF and MD were higher (**Supplementary Table 4**). However, there were metrics that demonstrated regionally-specific WMH-NAWM heterogeneity, particularly OD, FA, and g-ratio, for which the direction of the effects varied across lobes and WMHs layers. For example, OD was higher in WMH compared to NAWM in PVWM frontal regions (d = 0.98) but lower in SWM occipital regions (d = –3.71), suggesting that lesion-related differences in neurite orientation and fiber architecture are not uniform, but instead vary according to regional WM organization.

**Figure 2.**
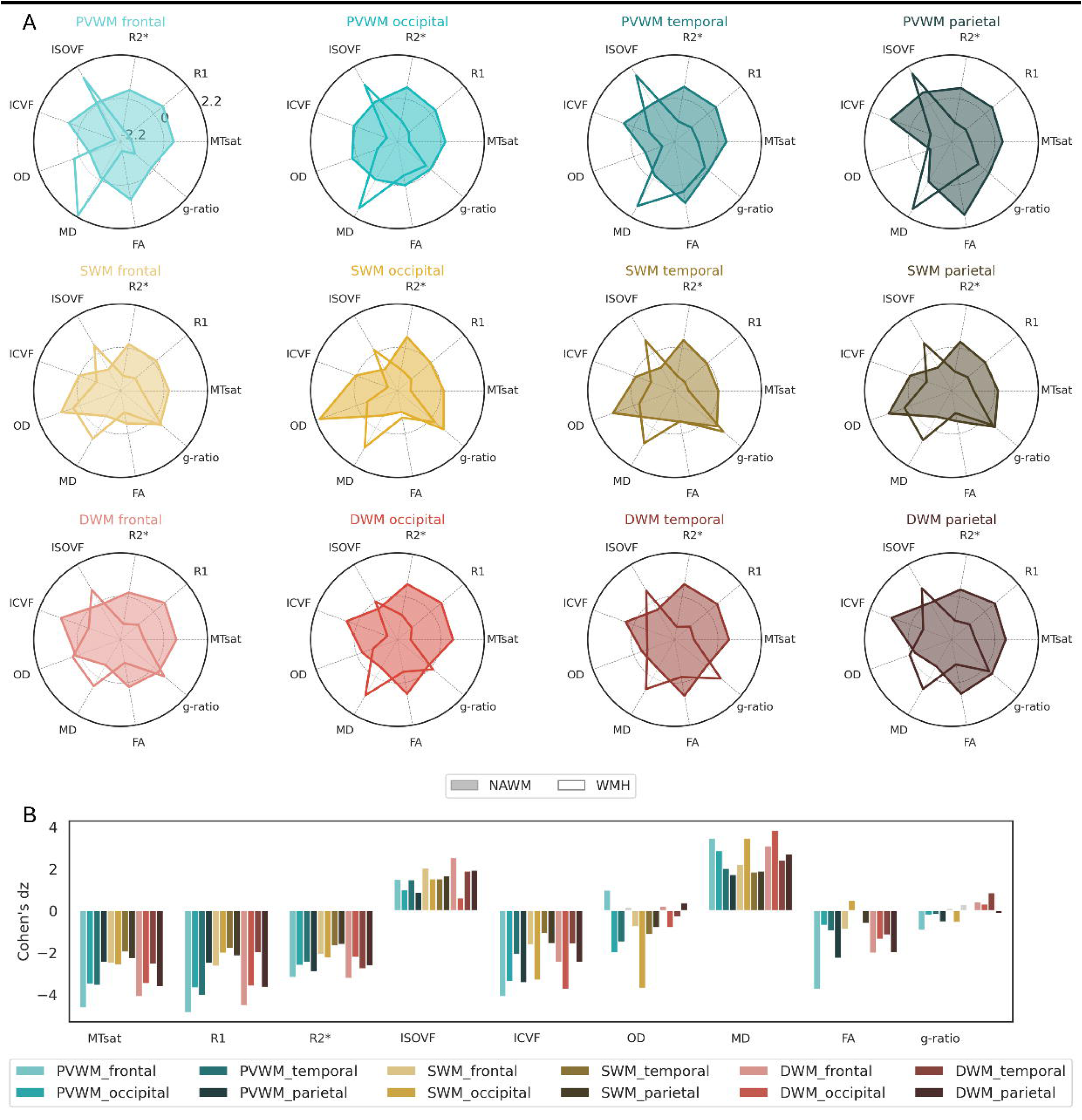
Regional comparison of white matter MRI microstructure metric between WMH and NAWM. **(A)** Mean values of MTsat, R1, R2*, ISOVF, ICVF, OD, MD, FA and g-ratio calculated for WMH and NAWM across lobar - frontal, occipital, temporal, parietal, and regional white matter subdivisions - SWM, DWM, PVWM. All metrics are z-scored once across the full dataset - all WMH and NAWM values from all regions, after which the analysis is restricted to participants with valid WMH-NAWM pairs for each region to calculate the regional microstructural profiles for each tissue type. **(B)** Group differences (WMH vs NAWM) quantified using Cohen’s dz. Coloured bars indicate regions with group differences after FDR correction (p<0.05); non-significant effects are shown in grey (OD PVWM parietal, FA SWM temporal, g-ratio SWM frontal and DWM occipital). Abbreviations: white matter hyperintensities (WMH), normal-appearing white matter (NAWM), magnetization transfer saturation (MTsat), longitudinal relaxation (R1), transverse relaxation (R2*), isotropic volume fraction (ISOVF), intracellular volume fraction (ICVF), orientation dispersion (OD), fractional anisotropy (FA), mean diffusivity (MD), superficial white matter (SWM), deep white matter (DWM), periventricular white matter (PVWM).

### Associations between WMH load and regional microstructural signature in NAWM and WMH

The WMH load, expressed by the logarithmic WMH volume per region and participant, showed marked regional differences (Kruskal–Wallis H = 290.66, p < 0.001; **Figure 3A, Supplementary Table 5**). WMH load was highest in the frontal PVWM (mean log volume ± SD: 5.76 ± 2.29) and lowest in SWM temporal regions (2.20 ± 1.35). To contextualize these differences, we examined the total volume of NAWM across WM partitions (log-transformed; Kruskal–Wallis H = 3866.35, p < 0.001). As expected, NAWM volumes varied substantially by partition due to anatomical definitions, with SWM regions showing the largest volumes (frontal SWM: 11.52 ± 0.15) and occipital DWM the smallest (8.41 ± 0.54; **Figure 3B**). Importantly, the volume fraction of WMHs in each WM region was highest in the PVWM parietal (3.51%, **Figure 3C**), highlighting the highest vulnerability of PVWM regions.

**Figure 3.**
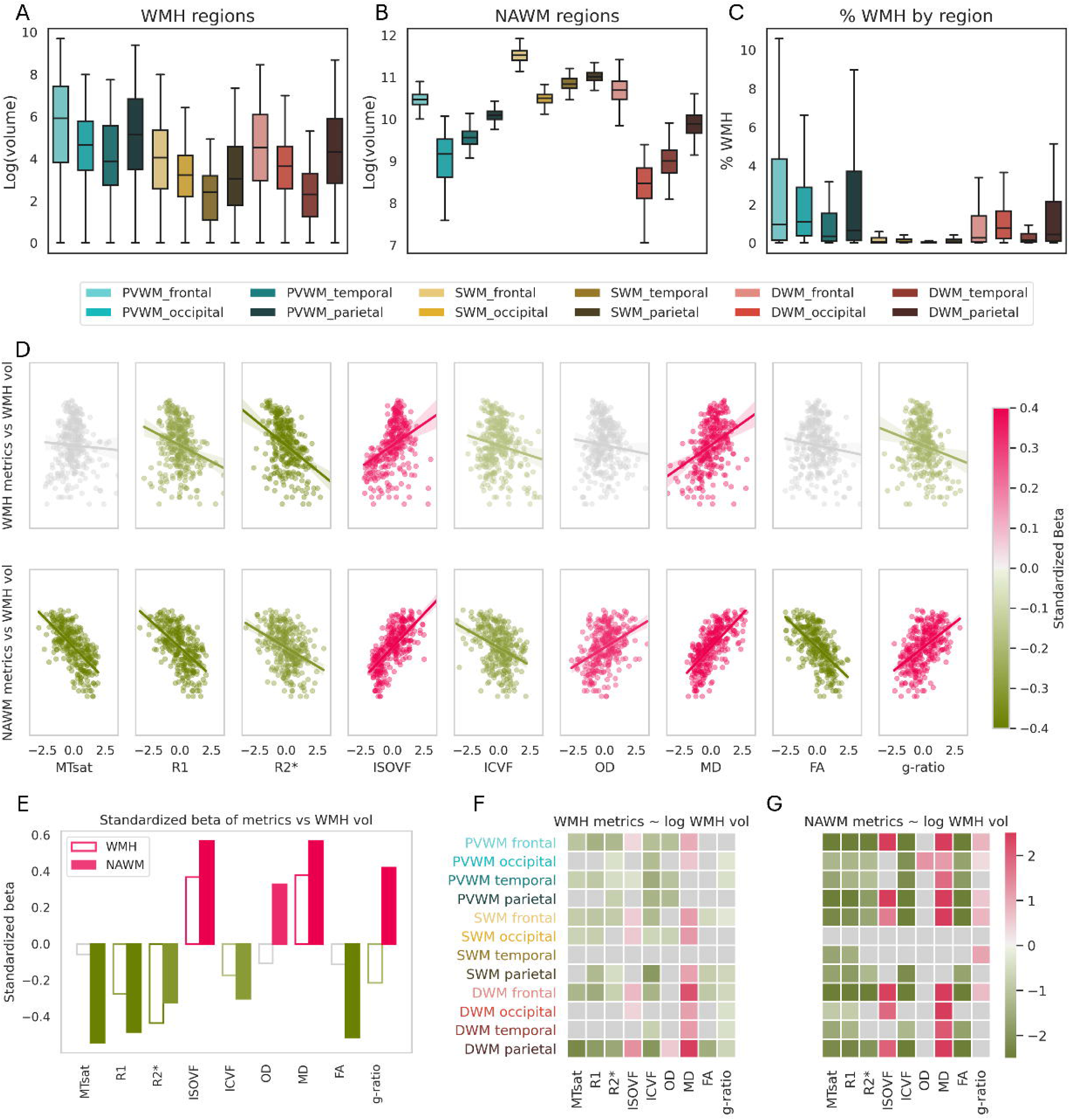
Regional distributions and associations of WMH and NAWM microstructure. **(A)** Log-transformed volumes of WMH masks across the same regions. **(B)** Log-transformed volumes of NAWM masks across lobar and regional white matter subdivisions. **(C)** Proportion of WMH volume relative to total white matter volume per region. Data are presented as boxplots, where the central line indicates the median, the box represents the interquartile range (Q1–Q3), and the whiskers extend to 1.5× the interquartile range. **(D)** Linear relationships between regional WMH volume and microstructural metrics. **(E)** Standardized beta coefficients summarizing the strength and direction of these associations for both WMH and NAWM. **(F)** Heatmap of standardized beta values for WMH microstructure in relation to log WMH volume, highlighting region-dependent effects. **(G)** Heatmap of standardized beta values for NAWM microstructure in relation to log WMH volume, also showing region-specific variation. Coloured bars and plots indicate statistically significant associations after FDR correction (p < 0.05); non-significant results are shown in grey. ***Abbreviations:*** *white matter hyperintensities (WMH), normal-appearing white matter (NAWM), magnetization transfer saturation (MTsat), longitudinal relaxation (R1), transverse relaxation (R2*), isotropic volume fraction (ISOVF), intracellular volume fraction (ICVF), orientation dispersion (OD), fractional anisotropy (FA), mean diffusivity (MD), superficial white matter (SWM), deep white matter (DWM), periventricular white matter (PVWM).*

We next examined whether regional WMH load is associated with the magnitude of microstructural alterations both within lesions and in the surrounding NAWM. Here, we tested associations between regional WMH volume and quantitative MRI metrics (**Figure 3D**). Greater WMH volume was associated with higher MD (+0.43) and ISOVF (+0.48) and lower R1 (−0.19), R2* (−0.43), ICVF (−0.08), and g-ratio (−0.24). Importantly, greater WMH load was also accompanied by more pronounced alterations in the surrounding NAWM, with lower MTsat (−0.73), R1 (−0.65), R2* (−0.39), ICVF (−0.36), FA (−0.68), and higher ISOVF (+0.77), MD (+0.73) and g-ratio (+0.58). We further tested these associations at the regional and lobar levels (**Figures 3F-G**). The strongest effects were observed in NAWM PVWM regions, where MD showed the largest positive association (β = +3.85) and R1 the largest negative association in the PVWM frontal (β = −4.87). In contrast, SWM occipital region showed no significant associations across metrics (*p* > 0.05).

### Regional WM microstructure in WMH and NAWM

Regional microstructural PCA of qMRI metrics in WMH and NAWM identified two components: PC1, explaining 58.1% of the variance (*p* = 0.0001 after 10,000 permutations), and PC2, explaining 21.5% (*p* = 0.0001). PC1 reflected a pattern of low MTsat, R1, R2*, ICVF, and FA, combined with high ISOVF and MD, with minimal contributions from g-ratio and OD (**Figure 4B**). This profile showed the same directionality as the Cohen’s dz contrasts between WMH and NAWM in **Figure 2B**. PC2, in contrast, represented low ISOVF, FA, and MD alongside high OD and g-ratio (**Figure 4C**). Notably, relaxometry metrics (MTsat, R1, R2*) contributed minimally to PC2, indicating that this axis of variation is largely independent of myelin- and iron-related processes, and instead captures differences in white matter geometry and fiber organization.

**Figure 4.**
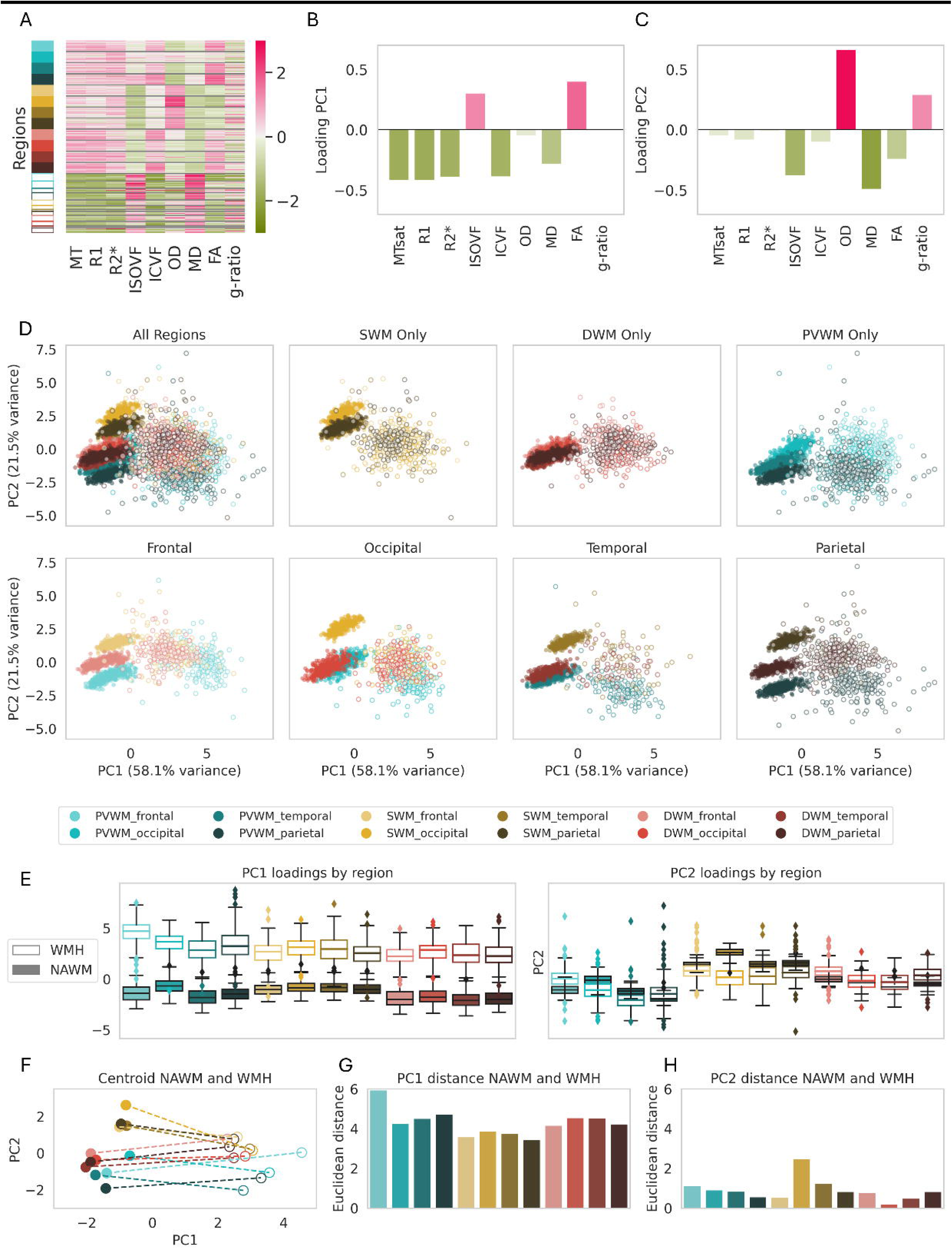
Principal component analysis of regional white matter microstructure in WMH and NAWM. We performed PCA on region-wise microstructural metrics across participants, using one NAWM measurement per region per individual, and multiple WMH regions per individual depending on the distribution of lesions. Each point represents a region-based microstructural profile (one observation per participant × region × tissue); NAWM profiles are available for all regions and participants, whereas WMH profiles are present only in regions containing lesions. **(A)** Heatmap of the standardized input matrix (regions × metrics) used for PCA. **(B-C)** Barplots showing the contribution (loadings) of each metric to PC1 and PC2. **(D)** Projection of all regional WMH and NAWM samples onto the PC1-PC2 space. **(E)** Boxplots showing the distribution of PC1 and PC2 scores across white matter regions for WMH and NAWM. **(F)** Regional centroids in PC space summarizing the average WMH (empty dots) and NAWM (solid dots) profile per region. **(G-H)** Barplots showing the Euclidean distance between paired WMH and NAWM centroids along PC1 and PC2, quantifying microstructural divergence across regions. ***Abbreviations:*** *white matter hyperintensities (WMH), normal-appearing white matter (NAWM), magnetization transfer saturation (MTsat), longitudinal relaxation (R1), transverse relaxation (R2*), isotropic volume fraction (ISOVF), intracellular volume fraction (ICVF), orientation dispersion (OD), fractional anisotropy (FA), mean diffusivity (MD), superficial white matter (SWM), deep white matter (DWM), periventricular white matter (PVWM), principal component (PC).*

**Figure 5.**
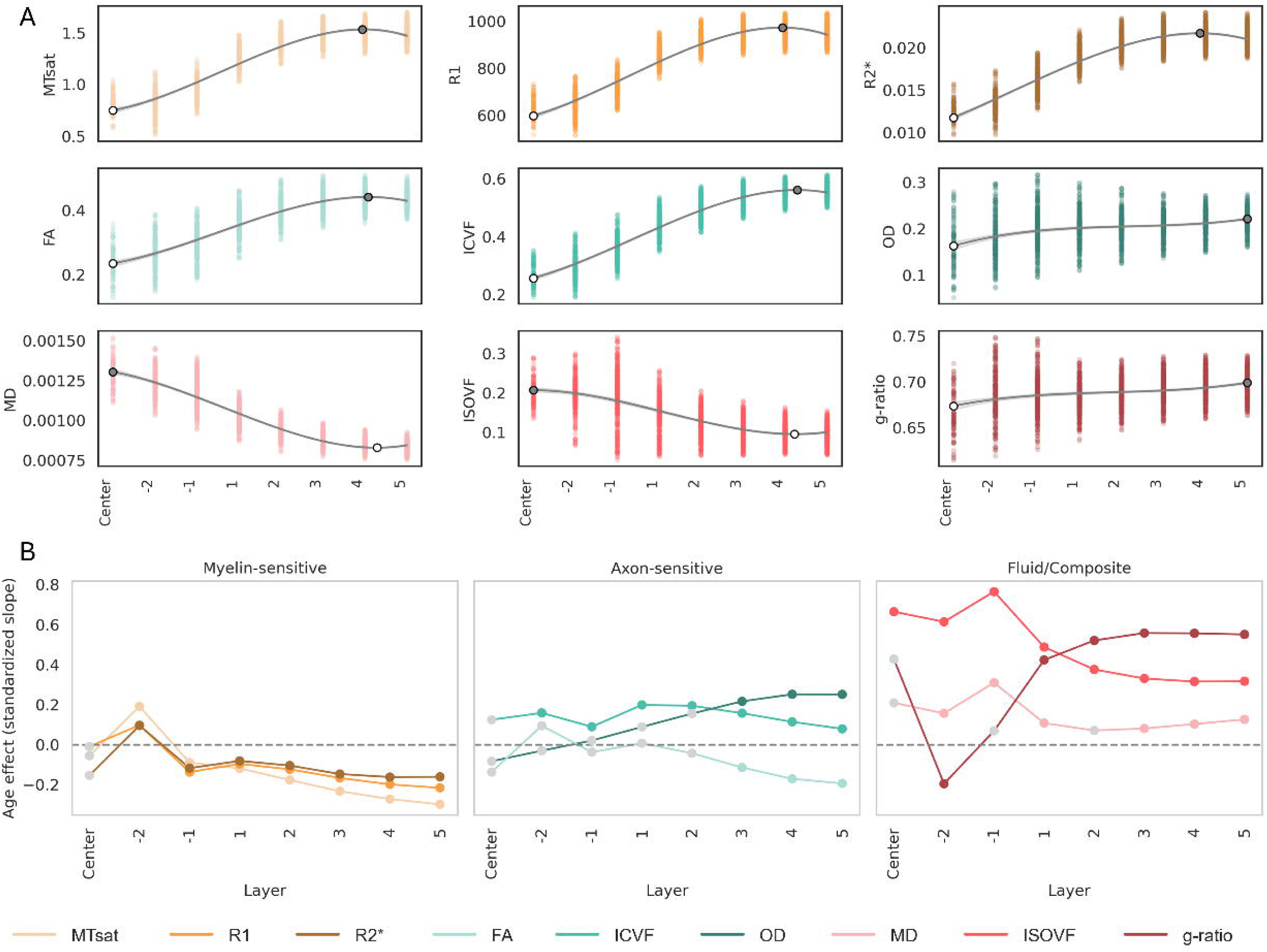
Layer-based tissue microstructural changes in the WMH penumbra. **(A)** Scatterplots showing layer-based WMH microstructural profiles across metrics, modeled using third-order polynomial fits. Layers span from the WMH center, through inner layers (−2, −1), to perilesional layers (1-5). White and grey dots indicate the layers at which the minimum and maximum metric values were observed, respectively. **(B)** Layer-specific age effects were tested using linear mixed-effects models that included standardized age, lesion layer, their interaction, and standardized lesion volume as fixed effects, and both random intercepts and random age slopes per participant. For each metric and layer, the age slope was computed by combining the fixed-effect age coefficient in the reference layer (Outer5) with the corresponding age × layer interaction term. Statistical significance of each age slope was assessed using linear contrasts and controlled for multiple comparisons using FDR across all metrics and layers. Lines depict the standardized age effect across layers; coloured points mark layers showing significant associations after correction. ***Abbreviations****: magnetization transfer saturation (MTsat), longitudinal relaxation (R1), transverse relaxation (R2*), isotropic volume fraction (ISOVF), intracellular volume fraction (ICVF), orientation dispersion (OD), fractional anisotropy (FA), mean diffusivity (MD).*

**Figure 6:**
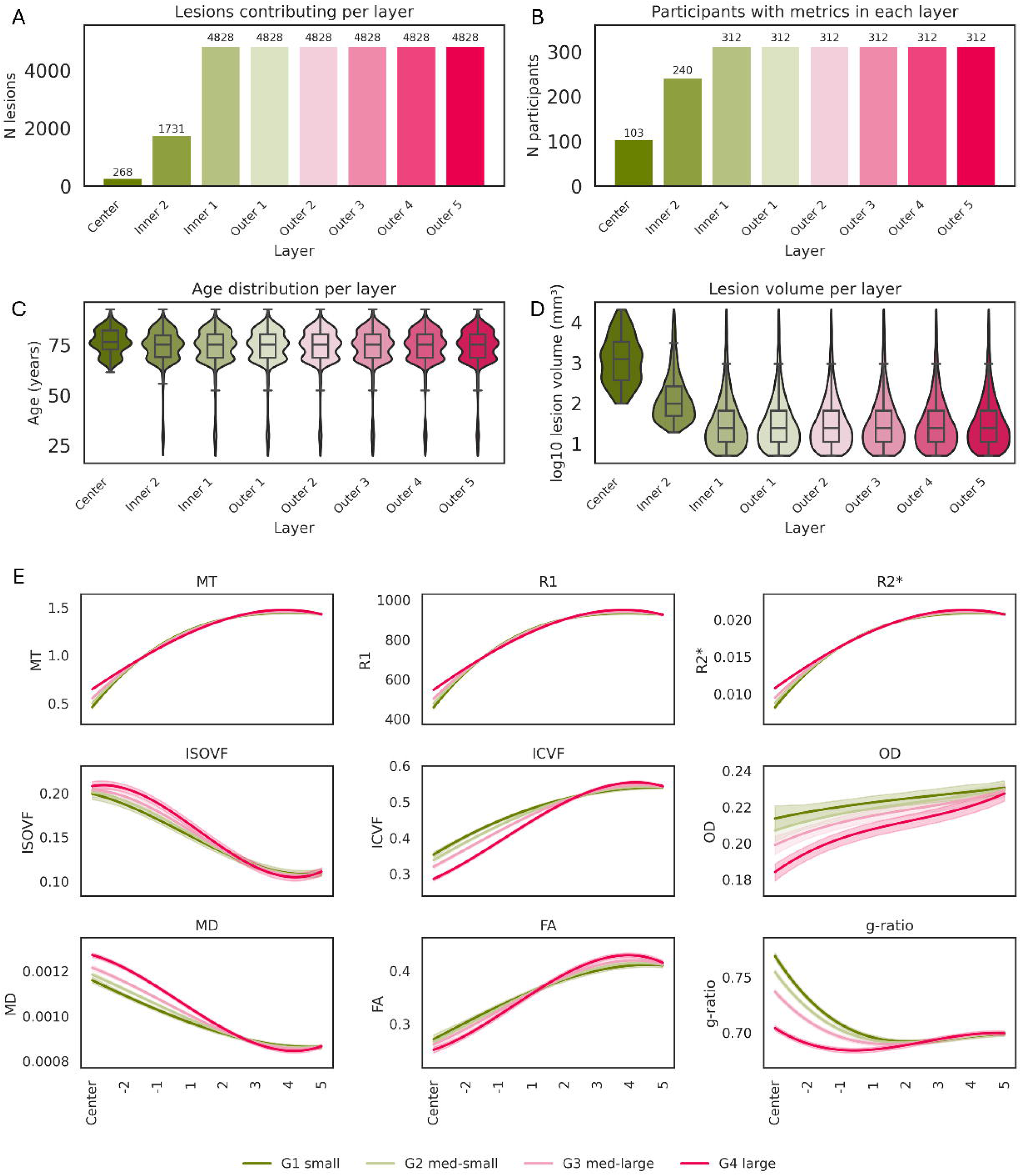
Layer-wise microstructural profiles across lesion-size using the geodesic per-lesion distance method. **(A)** Total number of valid lesion contributions to each layer measurements. **(B)** Number of participants contributing at least one lesion to each layer. **(C)** Participant age distribution per layer and **(D)** Lesion volume distribution to each layer’s contribution. **(E)** For each microstructural metric, smooth depth-profiles are shown from the WMH center to surrounding NAWM, stratified by lesion size. Curves are derived from a mixed-effects model in which metric values are predicted by lesion volume and a cubic polynomial of geodesic layer position (including their interaction), with random intercepts for participant and lesion. Shaded bands show 95% confidence intervals.

PC space projections revealed distinct regional qMRI signatures for WMH and NAWM (Figure 4D) with PC1 primarily reflecting the WMH – NAWM contrast (WMH mean = 3.06 vs. NAWM = −1.36; Mann–Whitney U = 7,684,158, p < 0.0001). Regional differences were less pronounced for WMH in DWM, suggesting reduced tissue microstructural variability in this compartment, especially along PC2. The greatest NAWM-WMH divergence along PC1 was observed in frontal PVWM (Euclidean distance = 5.95; **Figures 4G**). This is also the region that shows the strongest associations between WMH volume and microstructural alterations in NAWM (**Figure 3G**). In contrast, the smallest divergence NAWM-WMH was in the parietal SWM (distance = 4.23). PC2 also contrasted WMH and NAWM (U = 3,497,437, p < 0.0001), with the greatest NAWM-WMH difference along PC2 observed in the frontal PVWM region (Euclidean distance = 1.12 ; **Figures 4H**).

To further contextualize these components, we examined how PC1 and PC2 relate to age, sex, tissue class, and total WMH load using mixed-effects modelling (**Supplementary Table 6**). Age showed strong positive associations with both components, whereas sex had no detectable effect. WMH voxels exhibited substantially higher PC1 loadings and moderately higher PC2 loadings than NAWM, with a marked age-by-tissue interaction indicating attenuated age effects within WMH compared to NAWM. Total WMH volume was associated with PC1 but not PC2, suggesting that these components capture distinct sources of microstructural variation. PC1 reflects differences related to lesion load while PC2 captures variation independent of it. PC2 likely reflects differences in tissue organization or architecture, consistent with its associations with age and spatial distribution.

### Layer-based microstructural WMH profiles

We examined the spatial profile of microstructural differences from the WMH center outwards using a layer-centred analysis. In **Figure 5A**, we tested the difference between the layers vs the center per metric, and found differences across layers for all metrics (FDR-corrected p < 0.01 for all layers in all metrics except for ISOVF layer −2 vs center). Myelin-and axon-sensitive metrics (MTsat, R1, R2*, ICVF, FA) increased progressively from the WMH center toward NAWM, while markers of extracellular water content (MD, ISOVF) decreased along the same gradient. Most metrics reached their highest or lowest values either at the lesion center or in distant NAWM.

Across MRI contrasts, age showed a coherent and spatially structured pattern of associations with white-matter microstructure (**Figure 5B**). Myelin- and axon-related metrics (MT, R1, R2*, FA) exhibited their strongest negative age effects in the outer layers, whereas g-ratio and OD showed their strongest positive age effects in these same regions, indicating that age-related differences in microstructural organization were most pronounced at greater distances from the lesion centre (**Supplementary Figure 21**). Toward the center, these associations generally weakened, and in some inner layers they attenuated or shifted direction. In contrast, fluid-sensitive metrics (MD, ISOVF) demonstrated positive age effects across nearly all layers, with the largest increases within the lesion itself, supporting the notion that age-related changes in diffusivity and free-water content are most prominent in the lesion core and its immediate surroundings. ICVF also showed predominantly positive age slopes across layers -a pattern that differs from typical cross-sectional contrasts between WMH and normal-appearing tissue. Together, these findings highlighted distinct spatial signatures of age across WM microstructural domains, with outer-layer vulnerability in myelin- and axon-sensitive measures and core-centered increases in fluid-related metrics.

### Association between microstructural gradients and lesion size

To investigate whether the depth-dependent analyses depend on the number of lesions at each layer, we first quantified the sampling characteristics across geodesic depth. Across all WMH lesions, we observed a dependency of layer sampling on WMH size. The progressive reduction in lesion count toward the inner layers (**Figure 6A**) reflects the limited geodesic depth of smaller WMHs that precludes their contribution to the inner layer analysis. This resulted in only one-third of participants contributing to the center layer (**Figure 6B**). Conversely, all participants contributed data to the outermost layers, as NAWM masking only partially truncates these layers without eliminating them entirely. The participants contributing to inner-layer analysis were, on average, older (**Figure 6C**) and had larger lesions (**Figure 6D**). Thus, the sampling density across layers is not uniform but is systematically shaped by lesion size, with deeper layers disproportionately reflecting measurements from larger WMH.

Lesion size also influenced the magnitude and shape of the depth-dependent tissue microstructural gradients (**Figure 6E**). Across all nine microstructural metrics, mixed-effects models showed significant volume-by-layer interactions (FDR-corrected p<0.001, **Supplementary Table 7**), indicating that the WMH-to-NAWM gradients vary with lesion size. The corresponding unsmoothed layer-wise profiles, together with the underlying raw data distributions, are shown in **Supplementary Figure 22**. Larger lesions exhibited steeper and more abnormal center-to-periphery profiles, whereas smaller lesions showed shallower but still clearly detectable gradients. Notably, gradients persisted in the outer layers, where lesions of all sizes are represented, confirming that depth-dependent microstructural variation is a general property of WMH rather than being driven only by large lesions.

Similarly, while we observed baseline shifts in NAWM for myelin-related metrics, and center-layer differences for oedema-related metrics, demonstrating disease severity, the layer-specific gradients from NAWM to WMH center follow a very similar trajectory across all Fazekas scores (**Supplementary Figures 23** and **24**). Together, these findings indicated that the spatial microstructural gradient is a consistent feature of WMH pathology. While lesion size and load influenced the magnitude of effects, particularly in deeper layers, the overall pattern of microstructural change across layers remained stable.

### Associations between WM microstructure and cardiovascular risk

Figure 7 summarizes the latent variable (LV1) linking WM microstructure with cardiovascular and physiological measures. On the brain side (Figure 7A), LV1 was dominated by NAWM alterations, characterized by lower MTsat, R1, R2*, ICVF, and FA, together with higher g-ratio, MD, and ISOVF. WMH contributed modestly, with lower R2*, OD, and R1, and higher ISOVF and MD. From cardiovascular/physiological point of view (Figure 7B), LV1 was associated with higher age, systolic blood pressure, and bioimpedance, while hemoglobin showed a negative association. These associations appeared more pronounced in women, although the effect was weak.

**Figure 7:**
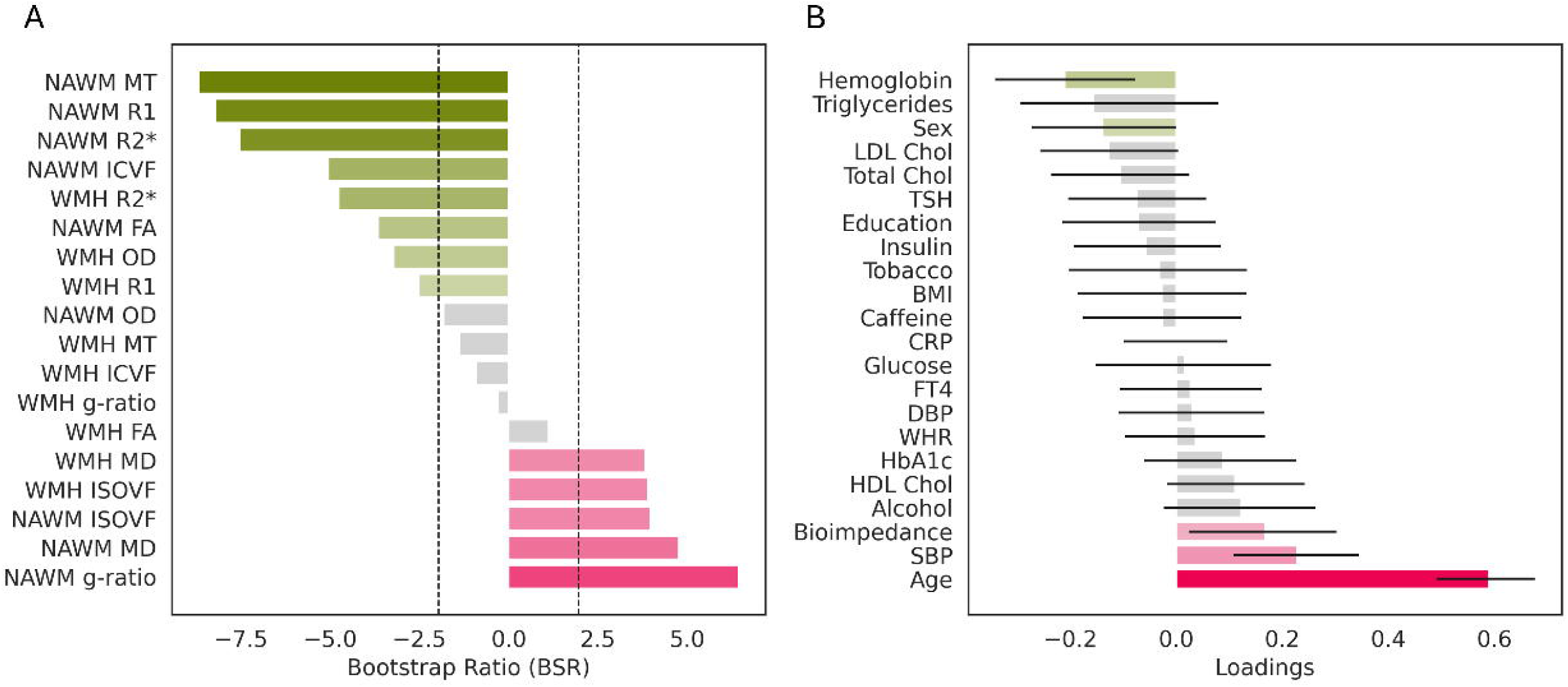
Partial least squares (PLS) analysis linking regional white matter microstructure to cardiovascular and medication-related variables. **(A)** Bootstrap ratio (BSR) plot showing the contribution of MRI metrics to the first PLS component explaining 61.7% of the variance. Lower NAWM R1, R2*, ICVF, FA, and MTsat, and higher NAWM MD, ISOVF, and g-ratio, contribute most strongly. WMH R1 and OD show negative loadings, while WMH MD and ISOVF load positively. Significant contributors (|BSR| > 1.96) are colored; non-significant in light gray. **(B)** Loadings for cardiovascular and medication variables on the first PLS component. Significant variables (95% CI not crossing zero) are colored; others in light gray. This pattern associates lower myelin and axon integrity and higher fluid-related metrics with increased age, systolic blood pressure (SBP), bioimpedance, and with lower hemoglobin. ***Abbreviations:*** *white matter hyperintensities (WMH), normal-appearing white matter (NAWM), magnetization transfer saturation (MTsat), longitudinal relaxation (R1), transverse relaxation (R2*), isotropic volume fraction (ISOVF), intracellular volume fraction (ICVF), orientation dispersion (OD), fractional anisotropy (FA), mean diffusivity (MD), superficial white matter (SWM), deep white matter (DWM), periventricular white matter (PVWM), bootstrap ratio (BSR), systolic (SBP) and diastolic blood pressure (DBP), body mass index (BMI), waist-to-hip ratio (WHR), glycated hemoglobin (HbA1c), high-sensitivity C-reactive protein (CRP), hemoglobin, thyroid-stimulating hormone (TSH), free thyroxine (FT4).*

## Discussion

In this community-based cohort, we provide a detailed regional characterization of WMH and NAWM microstructure in aging-associated SVD. Our analyses show that WMHs are associated with reduced myelin- and axon-sensitive measures and increased extracellular water, varying with distance from the lesion core. We also demonstrated that NAWM in individuals with WMH have regionally-specific pathology that correlates with the lesions volume. Finally, we provide significant evidence for the strong association between CVRs, and WM microstructure. Together, these findings advance the characterisation of SVD-related WM damage by providing a multiparametric and spatially resolved account of lesion-centred microstructural gradients across regions and tissue compartments whilst complementing and extending the existing literature on the WMH penumbra.

### Damage extends beyond visible lesions

In line with the gradient model, WMHs show gradual rather than abrupt tissue changes, with distinct microstructural profiles across core and peripheral layers. While previous work brought evidence for centrifugal gradients of tissue damage,^10,11,53^ we extend these findings by showing that lesion layers follow coherent age-sensitive patterns. WMH cores were characterised by age-independent loss of myelin. One interpretation is that myelin is already markedly reduced in these regions, limiting detectable age-related decline. Nonetheless, establishing whether this represents an irreversible stage of tissue pathology cannot be determined from cross-sectional data. In contrast, outer layers showed age-related differences in demyelination and axonal rarefaction, suggesting increased tissue vulnerability in regions surrounding the lesion. Extracellular water behaved differently, accumulating in both lesion cores and at intermediate distances, consistent with gliosis and extracellular matrix reorganization. Together, these findings refine the “WMH penumbra” model,^11,12^ showing that lesion centers, the areas captured by FLAIR imaging, represent an end-stage phenomenon, whereas the periphery remains dynamic, with age-sensitive zones of tissue vulnerability. This has direct implications for lesion tracking, as detection based on conventional MRI would likely underestimate the true extent of WM pathology.

### Cardiovascular drivers of microstructural damage

Our analysis demonstrated that WM microstructure covaries not only with local lesion properties but also with systemic vascular and metabolic health markers. Lower NAWM myelin content, axonal disruption, and higher extracellular water were associated with older age, elevated systolic blood pressure (SBP), being female, and reduced hemoglobin. The findings indicate that age-related WM differences occur alongside cardiovascular and metabolic factors showing further associations with local lesion characteristics.

In line with the existing empirical evidence, higher systolic blood pressure^54^ was strongly associated with tissue microstructural damage, consistent with the established link between hypertension and endothelial dysfunction, hypoperfusion, and blood-brain barrier disruption^55–57^. Low hemoglobin concentration, which may signal reduced oxygen-carrying capacity, was similarly associated with poorer WM microstructure, in line with evidence linking anemia-related hypoxic stress to WM vulnerability^58,59^. Together, these results support the notion that vascular, metabolic, and hematologic stressors are closely associated with the microstructural patterns observed in cerebral SVD.

### Microstructural fingerprints of WMH and NAWM

Our results challenge the notion of “normal” appearing WM by demonstrating that NAWM, although labelled as ‘normal’ on conventional imaging, should be understood as a descriptive radiological term rather than a biological assumption of intact tissue. In our data, NAWM exhibits subtle but systematic microstructural pathology that scales with WMH volume. Specifically, lower myelin- and axon-sensitive metrics and higher extracellular water indices in NAWM mirrored, albeit less severely, the characteristic profile seen within WMHs. This diffuse pattern is consistent with prior reports of NAWM abnormalities and suggests that WMH load^9,60–62^ reflects widespread tissue compromise rather than focal lesions. Histopathological evidence supports these observations, showing that tissue beyond lesion borders may already harbor myelin rarefaction, axonal degeneration, and gliosis.^63^ Together, these findings underscore the importance of moving beyond lesion-centric analyses: interventions and risk assessments should target the global WM network, as NAWM pathology may reflect a more diffuse manifestation of SVD with clinical relevance for aging and dementia risk. They also highlight the added value of quantitative, non-FLAIR imaging, since FLAIR captures only the lesion cores and misses diffuse pathology in surrounding tissue.

### Distinct regional patterns revealed by multivariate analysis

Multivariate and regional analyses revealed anatomical vulnerabilities shaping WMH-related pathology. The first principal component reflected a consistent pattern of myelin and axonal loss with elevated fluid-related metrics, clearly separating WMH from NAWM and peaking in periventricular regions. In contrast, the second component captured anatomical differences between SWM, DWM, and PVWM, largely driven by axonal orientation metrics, which may reflect either intrinsic tract organization or secondary axonal injury following myelin loss.^64^

Greater lesion load was associated with larger WMH–NAWM divergence, particularly in frontal PVWM, where NAWM microstructure was also most sensitive to lesion extent. This pattern was not explained by regional volume differences (**Supplementary** Figure 25), indicating true PVWM vulnerability. Although not a classical watershed zone,^65–67^ PVWM is especially susceptible to hypoperfusion-related pathology^68^ and may also be affected by venous collagenosis^69^, ependymal disruption^70,71^, and distal axonopathy.^72^ In contrast, SWM, especially in occipital and temporal lobes, showed weaker associations with lesion load, suggesting relative resilience or distinct pathogenic mechanisms.^73^

### Community-based sampling reveals early, diffuse changes

A key strength of our study is the use of the BrainLaus dataset, a community-based cohort derived from the general population of Lausanne, Switzerland. This cohort provides critical insights into the asymptomatic stages of WMH development, which are particularly relevant for primary prevention strategies.^1^ While a large number of WMH studies rely on clinical populations, which are highly valuable for understanding symptomatic or advanced disease, they are often subject to referral bias and may not reflect the broader aging population.^74^ In contrast, community-based samples offer greater generalizability but may be affected by healthy volunteer bias.^75,76^ Integrating evidence from both study types is essential to fully capture the spectrum of WMH progression and the systemic mechanisms that drive it.

### Limitations

We acknowledge that the number of participants contributing WMH data varies across regions and layers due to the biological distribution of WMH, with older individuals contributing more data in WMH-rich regions and deeper layers. As a result, statistical power is uneven, and findings in sparsely sampled areas should be interpreted cautiously. Ventricular enlargement in older individuals may affect the relative extent of PVWM and DWM, as the fixed distance boundary definition is sensitive to ventricular size^77^

PLS and PCA capture patterns of shared variance rather than discrete biological mechanisms and their output are sensitive to cohort characteristics. These results should be considered exploratory and require independent or longitudinal validation.

Although our cross-sectional analyses suggest tissue loss in WMH cores and degeneration at lesion edges, WMHs can expand, stabilize, or regress over time.^4,63,78^ Longitudinal studies are therefore needed to confirm whether these spatial patterns generalise across lesion trajectories. Although age analyses were adjusted for total WMH load, inner-layer data derive from larger lesions, more common in older participants, and should be interpreted with this sampling dependency in mind. This imbalance may influence layer-wise comparisons, such that deeper layer differences do not exclusively reflect spatial variation within lesions but may partly capture differences in lesion characteristics across participants. This potential sampling bias should be considered when interpreting microstructural patterns in inner lesion layers.

Automated WMH segmentation is less reliable at low lesion load, where small lesion volumes increase Dice sensitivity to minor misclassifications– a limitation recognised across methods, including WHITE-Net. As WMH load is generally lower in younger participants, segmentation accuracy may be reduced in this subgroup, representing a potential bias in the characterization of WMH microstructure. Segmentation uncertainty is higher at lesion boundaries, which may affect voxel assignment in near-boundary layers. Together with reduced reliability in small lesions, this may introduce some variability in layer-based analyses, particularly in younger participants. This effect is expected to remain confined to the immediate lesion edge, and is unlikely to substantially affect spatial gradients across layers. Diffusion tensor metrics including FA, are limited in regions with complex fiber architecture and susceptible to partial volume effects, particularly in superficial WM and near lesion boundaries; however, complementary use of NODDI indices and g-ratio partially mitigates these limitations.

As lesion volume does not directly capture WMH-related network disruption, tractography-derived disconnection metrics may provide more sensitive measures of network disruption, particularly for phenotypical and cardiovascular associations. Combining such approaches with microstructural holds the promise of a more comprehensive understanding of WMH-related pathology.

## Conclusions

WMHs delineate regions of WM vulnerability, characterised by myelin loss, axonal damage, and increased extracellular water, linked to cardiovascular and metabolic risk factors. Integrating spatial gradients of tissue microstructure and regional susceptibility, our findings provide a whole-brain framework for aging-related SVD. The predominant microstructural patterns - myelin loss and increased extracellular water, especially in lesion cores, point to these as principal drivers of the observed pathology, with axonal involvement playing a more variable role in the surrounding NAWM tissue.

## Data availability statement

Data from BrainLaus dataset is not publicly available but can be requested upon reasonable and formal request (https://www.colaus-psycolaus.ch/professionals/how-to-collaborate). All our data extraction, analysis code, results and figures can be found on GitHub (https://github.com/AurelieBussy/WMH_analyses).

## Acknowledgments

We wish to thank all BrainLaus participants for their participation in the study.

## Funding

B.D. is supported by the Swiss National Science Foundation (project grant no. 213595, 32003B_135679, 32003B_159780, 324730_192755 and CRSK-3_190185), InnoSuisse Flagship Swiss brAInHealth project, ERA_NET NEURON JTC2020: iSEE and JTC2023-ELSA: BrainTree projects. F.K. is funded by the PHASE IV AI/Horizon Europe grant, grant agreement ID: 1010953844:01 and Horizon 2020 (871643-MORPHEMIC). A.L. is supported by the Swiss National Science Foundation (Grant Nos. 320030_184784, CR00I5-235940). The Laboratory for Research in Neuroimaging (LREN) is very grateful to the Roger De Spoelberch and Partridge Foundations for their generous financial support.

## Competing interests

The authors report no competing interests.

## Supplementary material

Supplementary material is available at Brain online.

## Notes

### Competing Interest Statement

The authors have declared no competing interest.

### Summary of Updates

Summary of main changes Conceptual clarification. We refined the positioning of the study and clarified its novelty. Interpretation. We moderated causal and mechanistic statements to better reflect the cross-sectional design. Methods. We improved clarity and justification of key analytical and processing steps. Limitations. We expanded the discussion of methodological constraints and potential biases. Supplementary material. We added new figures (including unsmoothed profiles with raw data and polynomial fit selection) and expanded supporting analyses. Readability. We streamlined wording and improved clarity throughout the manuscript.

## References

1. de Leeuw, F. E. et al. Prevalence of cerebral white matter lesions in elderly people: a population based magnetic resonance imaging study. The Rotterdam Scan Study. J. Neurol. Neurosurg. Psychiatry 70, 9–14 (2001).

2. Sharma, R., Sekhon, S., Lui, F. & Cascella, M. White Matter Lesions. in StatPearls (StatPearls Publishing, Treasure Island (FL), 2024).

3. Grinberg, L. T. & Thal, D. R. Vascular pathology in the aged human brain. Acta Neuropathol. 119, 277–290 (2010).

4. Wardlaw, J. M., Smith, C. & Dichgans, M. Small vessel disease: mechanisms and clinical implications. Lancet Neurol. 18, 684–696 (2019).

5. Wartolowska, K. A. & Webb, A. J. S. Midlife blood pressure is associated with the severity of white matter hyperintensities: analysis of the UK Biobank cohort study. Eur. Heart J. 42, 750–757 (2021).

6. Lampe, L. et al. Visceral obesity relates to deep white matter hyperintensities via inflammation. Ann. Neurol. 85, 194–203 (2019).

7. Solé-Guardia, G. et al. Association between hypertension and neurovascular inflammation in both normal-appearing white matter and white matter hyperintensities. Acta Neuropathol. Commun. 11, 2 (2023).

8. Schilling, S. et al. Plasma lipids and cerebral small vessel disease. Neurology 83, 1844–1852 (2014).

9. Maniega, S. M. et al. White matter hyperintensities and normal-appearing white matter integrity in the aging brain. Neurobiol. Aging 36, 909–918 (2015).

10. Maillard, P. et al. White matter hyperintensities and their penumbra lie along a continuum of injury in the aging brain. Stroke 45, 1721–1726 (2014).

11. Maillard, P. et al. White matter hyperintensity penumbra. Stroke 42, 1917–1922 (2011).

12. Voorter, P. H. M. et al. Heterogeneity and penumbra of white matter hyperintensities in small vessel diseases determined by quantitative MRI. Stroke (2024) doi:10.1161/strokeaha.124.047910.

13. Habes, M. et al. White matter hyperintensities and imaging patterns of brain ageing in the general population. Brain 139, 1164–1179 (2016).

14. Veldsman, M., Kindalova, P., Husain, M., Kosmidis, I. & Nichols, T. E. Spatial distribution and cognitive impact of cerebrovascular risk-related white matter hyperintensities. NeuroImage Clin. 28, 102405 (2020).

15. Piguet, O. et al. White matter loss in healthy ageing: a postmortem analysis. Neurobiol. Aging 30, 1288–1295 (2009).

16. Hajnal, J. V. et al. Use of fluid attenuated inversion recovery (FLAIR) pulse sequences in MRI of the brain. J. Comput. Assist. Tomogr. 16, 841–844 (1992).

17. Wardlaw, J. M., Smith, C. & Dichgans, M. Mechanisms of sporadic cerebral small vessel disease: insights from neuroimaging. Lancet Neurol. 12, 483–497 (2013).

18. Iordanishvili, E. et al. Quantitative MRI of cerebral white matter hyperintensities: A new approach towards understanding the underlying pathology. Neuroimage 202, 116077 (2019).

19. Courtney, M. et al. Connecting the dots: microstructural properties of white matter hyperintensities predict longitudinal cognitive changes in ageing. Front. Aging Neurosci. 17, 1520069 (2025).

20. Callaghan, M. F. et al. Widespread age-related differences in the human brain microstructure revealed by quantitative magnetic resonance imaging. Neurobiol. Aging 35, 1862–1872 (2014).

21. Helms, G., Dathe, H. & Dechent, P. Quantitative FLASH MRI at 3T using a rational approximation of the Ernst equation. Magn. Reson. Med. 59, 667–672 (2008).

22. Lutti, A., Dick, F., Sereno, M. I. & Weiskopf, N. Using high-resolution quantitative mapping of R1 as an index of cortical myelination. Neuroimage 93, 176–188 (2014).

23. Fukunaga, M. et al. Layer-specific variation of iron content in cerebral cortex as a source of MRI contrast. Proc. Natl. Acad. Sci. U. S. A. 107, 3834–3839 (2010).

24. Stüber, C. et al. Myelin and iron concentration in the human brain: A quantitative study of MRI contrast. Neuroimage 93, 95–106 (2014).

25. Oliveira, R. et al. In vivo characterization of magnetic inclusions in the subcortex from nonexponential transverse relaxation decay. NMR Biomed. 38, e70051 (2025).

26. Parent, O. et al. Characterizing spatiotemporal white matter hyperintensity pathophysiology in vivo to disentangle vascular and neurodegenerative contributions. medRxiv 2025.06.10.25329342 (2025) doi:10.1101/2025.06.10.25329342.

27. Weiskopf, N., Edwards, L. J., Helms, G., Mohammadi, S. & Kirilina, E. Quantitative magnetic resonance imaging of brain anatomy and in vivo histology. Nat. Rev. Phys. 3, 570–588 (2021).

28. Firmann, M. et al. The CoLaus study: a population-based study to investigate the epidemiology and genetic determinants of cardiovascular risk factors and metabolic syndrome. BMC Cardiovasc. Disord. 8, 6 (2008).

29. Preisig, M. et al. The PsyCoLaus study: methodology and characteristics of the sample of a population-based survey on psychiatric disorders and their association with genetic and cardiovascular risk factors. BMC Psychiatry 9, 1–12 (2009).

30. Weiskopf, N. et al. Quantitative multi-parameter mapping of R1, PD(*), MT, and R2(*) at 3T: a multi-center validation. Front. Neurosci. 7, 95 (2013).

31. Draganski, B. et al. Regional specificity of MRI contrast parameter changes in normal ageing revealed by voxel-based quantification (VBQ). Neuroimage 55, 1423–1434 (2011).

32. Lutti, A., Hutton, C., Finsterbusch, J., Helms, G. & Weiskopf, N. Optimization and validation of methods for mapping of the radiofrequency transmit field at 3T. Magn. Reson. Med. 64, 229–238 (2010).

33. Lutti, A. et al. Robust and Fast Whole Brain Mapping of the RF Transmit Field B1 at 7T. PLoS One 7, e32379 (2012).

34. Slater, D. A. et al. Evolution of white matter tract microstructure across the life span. Hum. Brain Mapp. 40, 2252–2268 (2019).

35. Tabelow, K. et al. hMRI - A toolbox for quantitative MRI in neuroscience and clinical research. Neuroimage 194, 191–210 (2019).

36. Tofts, P. Quantitative MRI of the Brain: Measuring Changes Caused by Disease. (John Wiley & Sons, 2003). doi:10.1002/0470869526.

37. Tournier, J.-D. et al. MRtrix3: A fast, flexible and open software framework for medical image processing and visualisation. Neuroimage 202, 116137 (2019).

38. Andersson, J. L. R. & Sotiropoulos, S. N. An integrated approach to correction for off-resonance effects and subject movement in diffusion MR imaging. Neuroimage 125, 1063–1078 (2016).

39. Hutton, C. et al. Image distortion correction in fMRI: A quantitative evaluation. Neuroimage 16, 217–240 (2002).

40. Zhang, H., Schneider, T., Wheeler-Kingshott, C. A. & Alexander, D. C. NODDI: practical in vivo neurite orientation dispersion and density imaging of the human brain. Neuroimage 61, 1000–1016 (2012).

41. Daducci, A. et al. Accelerated Microstructure Imaging via Convex Optimization (AMICO) from diffusion MRI data. Neuroimage 105, 32–44 (2015).

42. Stikov, N. et al. In vivo histology of the myelin g-ratio with magnetic resonance imaging. Neuroimage 118, 397–405 (2015).

43. Statistical Parametric Mapping: The Analysis of Functional Brain Images. (Academic Press, San Diego, CA, 2007). doi:10.1016/b978-0-12-372560-8.x5000-1.

44. Manjón, J. V., Coupé, P., Martí-Bonmatí, L., Collins, D. L. & Robles, M. Adaptive non-local means denoising of MR images with spatially varying noise levels. J. Magn. Reson. Imaging 31, 192–203 (2010).

45. Jenkinson, M. & Smith, S. A global optimisation method for robust affine registration of brain images. Med. Image Anal. 5, 143–156 (2001).

46. Jenkinson, M., Bannister, P., Brady, M. & Smith, S. Improved optimization for the robust and accurate linear registration and motion correction of brain images. Neuroimage 17, 825–841 (2002).

47. Fazekas, F., Chawluk, J. B., Alavi, A., Hurtig, H. I. & Zimmerman, R. A. MR signal abnormalities at 1.5 T in Alzheimer’s dementia and normal aging. American Journal of Roentgenology 149, 351–356 (1987).

48. Cathala, C., Kherif, F., Thiran, J.-P., Bussy, A. & Draganski, B. WHITE-Net : White matter HyperIntensities Tissue Extraction using deep learning Network. medRxiv (2025) doi:10.1101/2025.01.09.25320242.

49. Yan, Y., Balbastre, Y., Brudfors, M. & Ashburner, J. Factorisation-based Image Labelling. arXiv [cs.CV] (2021).

50. McIntosh, A. R. & Lobaugh, N. J. Partial least squares analysis of neuroimaging data: applications and advances. Neuroimage 23 **Suppl 1**, S250–63 (2004).

51. Bussy, A. et al. Exploring morphological and microstructural signatures across the Alzheimer’s spectrum and risk factors. Neurobiol. Aging 149, 1–18 (2025).

52. Zeighami, Y. et al. A clinical-anatomical signature of Parkinson’s disease identified with partial least squares and magnetic resonance imaging. Neuroimage 190, 69–78 (2019).

53. Duering, M. et al. Incident lacunes preferentially localize to the edge of white matter hyperintensities: insights into the pathophysiology of cerebral small vessel disease. Brain 136, 2717–2726 (2013).

54. Mok, V. & Kim, J. S. Prevention and management of cerebral small vessel disease. J. Stroke 17, 111–122 (2015).

55. Zhang, C. E. et al. Blood-brain barrier leakage in relation to white matter hyperintensity volume and cognition in small vessel disease and normal aging. Brain Imaging Behav. 13, 389–395 (2019).

56. Li, Y. et al. Higher blood–brain barrier permeability is associated with higher white matter hyperintensities burden. J. Neurol. 264, 1474–1481 (2017).

57. Katsi, V. et al. Blood-brain barrier dysfunction: the undervalued frontier of hypertension. J. Hum. Hypertens. 34, 682–691 (2020).

58. Mairbäurl, H. & Weber, R. E. Oxygen transport by hemoglobin. Compr. Physiol. 2, 1463–1489 (2012).

59. Inzitari, M. et al. Anemia is associated with the progression of white matter disease in older adults with high blood pressure: the cardiovascular health study. J. Am. Geriatr. Soc. 56, 1867–1872 (2008).

60. Parent, O. et al. Assessment of white matter hyperintensity severity using multimodal magnetic resonance imaging. Brain Commun. 5, fcad279 (2023).

61. Bastin, M. E. et al. Diffusion tensor and magnetization transfer MRI measurements of periventricular white matter hyperintensities in old age. Neurobiol. Aging 30, 125–136 (2009).

62. Parent, O. et al. Characterizing spatiotemporal white matter hyperintensity pathophysiology in vivo to disentangle vascular and neurodegenerative contributions. medRxiv 2025.06.10.25329342 (2025) doi:10.1101/2025.06.10.25329342.

63. Wardlaw, J. M., Valdés Hernández, M. C. & Muñoz-Maniega, S. What are white matter hyperintensities made of? Relevance to vascular cognitive impairment. J. Am. Heart Assoc. 4, 001140 (2015).

64. Bullock, D. N. et al. A taxonomy of the brain’s white matter: twenty-one major tracts for the 21st century. Cereb. Cortex 32, 4524–4548 (2022).

65. Torvik, A. The pathogenesis of watershed infarcts in the brain. Stroke 15, 221–223 (1984).

66. Kang, P. et al. Oxygen metabolic stress and white matter injury in patients with cerebral small vessel disease. Stroke 53, 1570–1579 (2022).

67. Kim, K. W., MacFall, J. R. & Payne, M. E. Classification of white matter lesions on magnetic resonance imaging in elderly persons. Biol. Psychiatry 64, 273–280 (2008).

68. ten Dam, V. H. et al. Decline in Total Cerebral Blood Flow Is Linked with Increase in Periventricular but Not Deep White Matter Hyperintensities1. Radiology (2007) doi:10.1148/radiol.2431052111.

69. Keith, J. et al. Collagenosis of the Deep Medullary Veins: An Underrecognized Pathologic Correlate of White Matter Hyperintensities and Periventricular Infarction? J Neuropathol Exp Neurol 76, 299–312 (2017).

70. Jochems, A. C. C. et al. Magnetic resonance imaging tissue signatures associated with white matter changes due to sporadic cerebral small vessel disease indicate that white matter hyperintensities can regress. J. Am. Heart Assoc. 13, e032259 (2024).

71. Fazekas, F., Schmidt, R. & Scheltens, P. Pathophysiologic mechanisms in the development of age-related white matter changes of the brain. Dement Geriatr Cogn Disord 9 **Suppl 1**, 2–5 (1998).

72. Beach, T. G. et al. Cerebral white matter rarefaction has both neurodegenerative and vascular causes and may primarily be a distal axonopathy. J Neuropathol Exp Neurol 82, 457–466 (2023).

73. Moody, D. M., Bell, M. A. & Challa, V. R. Features of the cerebral vascular pattern that predict vulnerability to perfusion or oxygenation deficiency: an anatomic study. AJNR Am. J. Neuroradiol. 11, 431–439 (1990).

74. Kokmen, E., Ozsarfati, Y., Beard, C. M., O’Brien, P. C. & Rocca, W. A. Impact of referral bias on clinical and epidemiological studies of Alzheimer’s disease. J. Clin. Epidemiol. 49, 79–83 (1996).

75. Weuve, J. et al. Guidelines for reporting methodological challenges and evaluating potential bias in dementia research. Alzheimers. Dement. 11, 1098–1109 (2015).

76. Brayne, C. & Moffitt, T. E. The limitations of large-scale volunteer databases to address inequalities and global challenges in health and aging. *Nat*. Aging 2, 775–783 (2022).

77. Griffanti, L. et al. Classification and characterization of periventricular and deep white matter hyperintensities on MRI: A study in older adults. Neuroimage 170, 174–181 (2018).

78. Promjunyakul, N.-O. et al. Baseline NAWM structural integrity and CBF predict periventricular WMH expansion over time. Neurology 90, e2119–e2126 (2018).

