## Supplementary material for "White matter microstructure fingerprint of cerebral small vessel disease"

**Supplementary methods**

**Participant Selection**

The imaging database consisted of deidentified MRIs collected across several studies. We first identified 631 participants with an available FLAIR image, as this sequence was required for WMH segmentation. From this group, we selected participants belonging to the BrainLaus study, yielding 423 individuals. After applying visually-guided quality-control across all MR contrasts (FLAIR, R1, R2*, MTsat, and diffusion), 363 participants met the inclusion criteria. This full sample was used for all brain imaging analyses. Among these 363 individuals, 202 had complete cardiovascular risk (CVR) data; the participants without CVR information were part of a nested study that included younger individuals for whom CVR measures were not collected. This subset of 202 participants was therefore used exclusively for the PLS analyses linking CVR to brain microstructure.

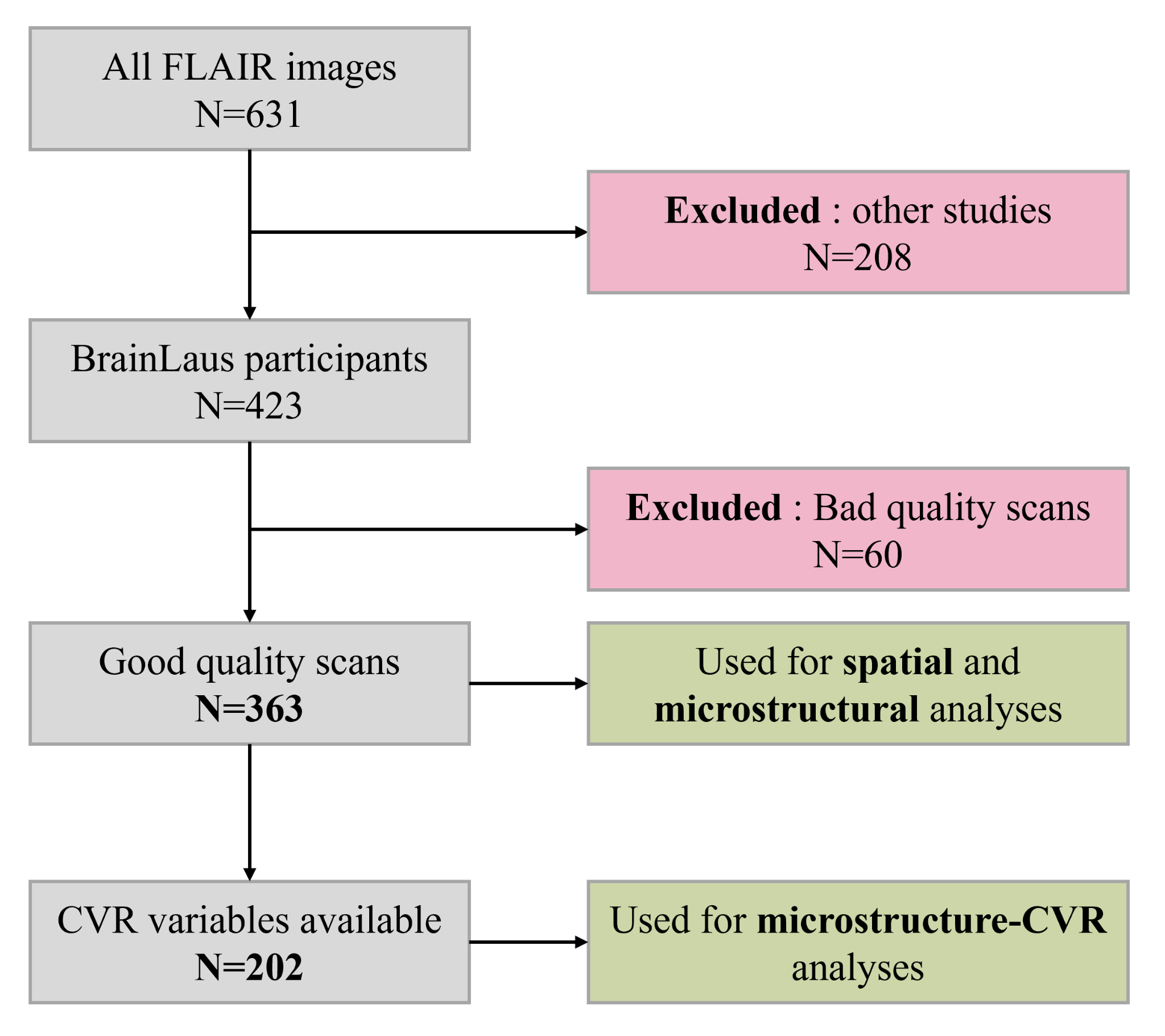

**Supplementary Figure 1.** Overview of participant inclusion and exclusion criteria and the resulting samples used across analyses. The final dataset used for the spatial and microstructural analyses includes all participants with high-quality FLAIR, MPM, and diffusion-derived maps and WMH segmentations (n = 363). A more restricted subset of participants with complete cardiovascular risk factor data (n = 202) was used for the PLS analyses examining associations between cardiovascular burden and microstructure. This schematic clarifies the rationale for the two analytic samples and how each was derived.

**MRI quality control**

To investigate potential selection bias caused by quality control assessment, we compared the demographic characteristics and Fazekas scores along the different steps of participants’ selection. As shown in **Supplementary Figure 2** and **Supplementary Table 1**, the reduced sample after QC did not differ in age, sex distribution, or Fazekas scores compared to the full set. Given that the younger participants of a nested study lack CVR assessments. Their exclusion resulted in a final sample that was older and had, on average, larger lesion loads. This pattern reflects the expected effect of age rather than any bias introduced by QC-related participants exclusion.

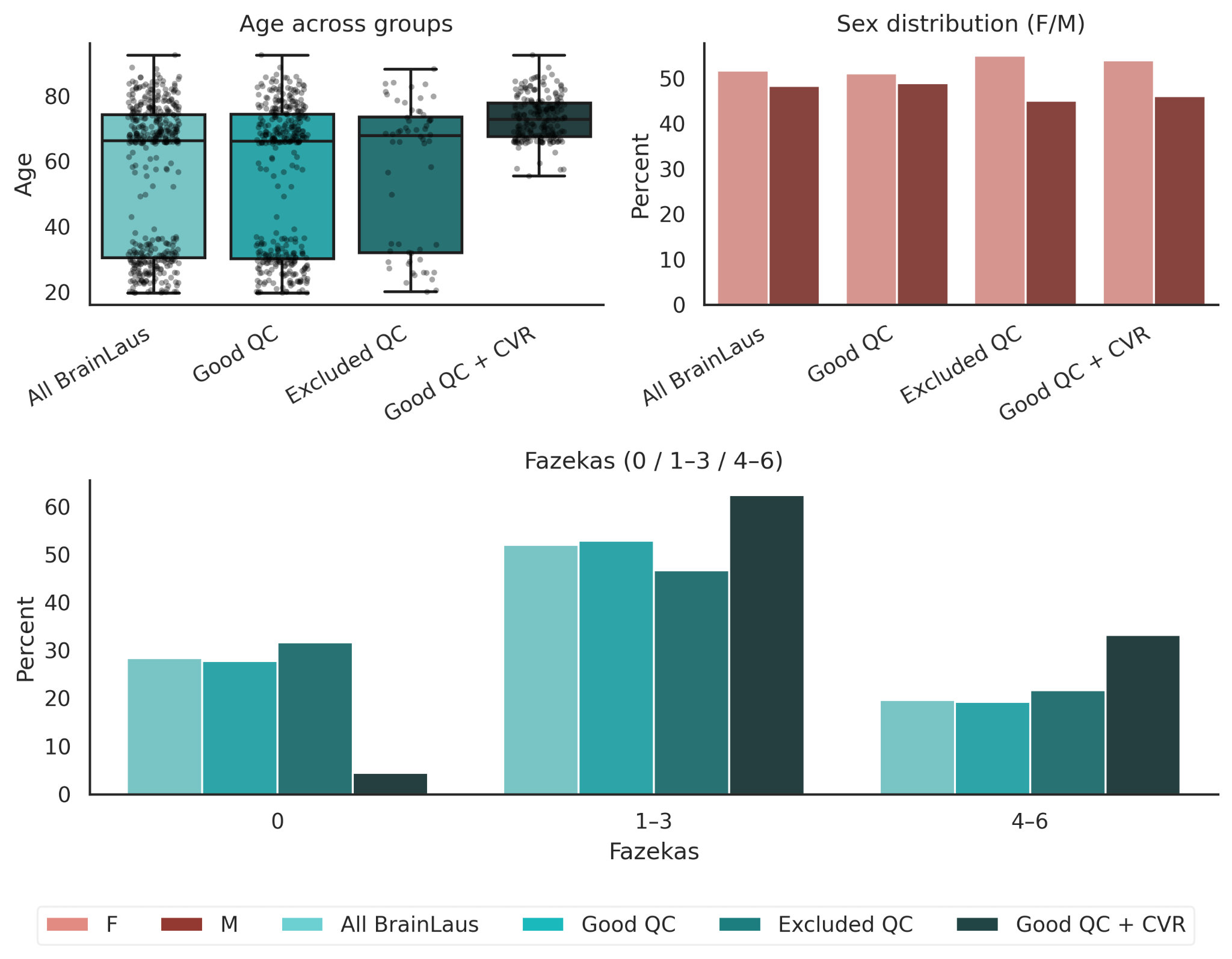

**Supplementary Figure 2**: Comparison between subsets of the BrainLaus cohort: the full sample (n = 423), participants who passed quality control (QC < 3; n = 363), participants excluded due to QC (n = 60), and participants who passed QC < 3 and had CVR data available (n = 202). Top left: age distribution; top right: sex distribution; bottom: Fazekas score comparison.

**Supplementary Table 1**: Statistical comparison between subsets of the BrainLaus cohort: the full sample (n = 423), participants who passed quality control (QC < 3; n = 363), participants excluded due to QC (n = 60), and participants who passed QC < 3 and had CVR data available (n = 202). Comparisons were performed for age (Welch’s t-test), and for sex and Fazekas score proportions (Chi-square tests).

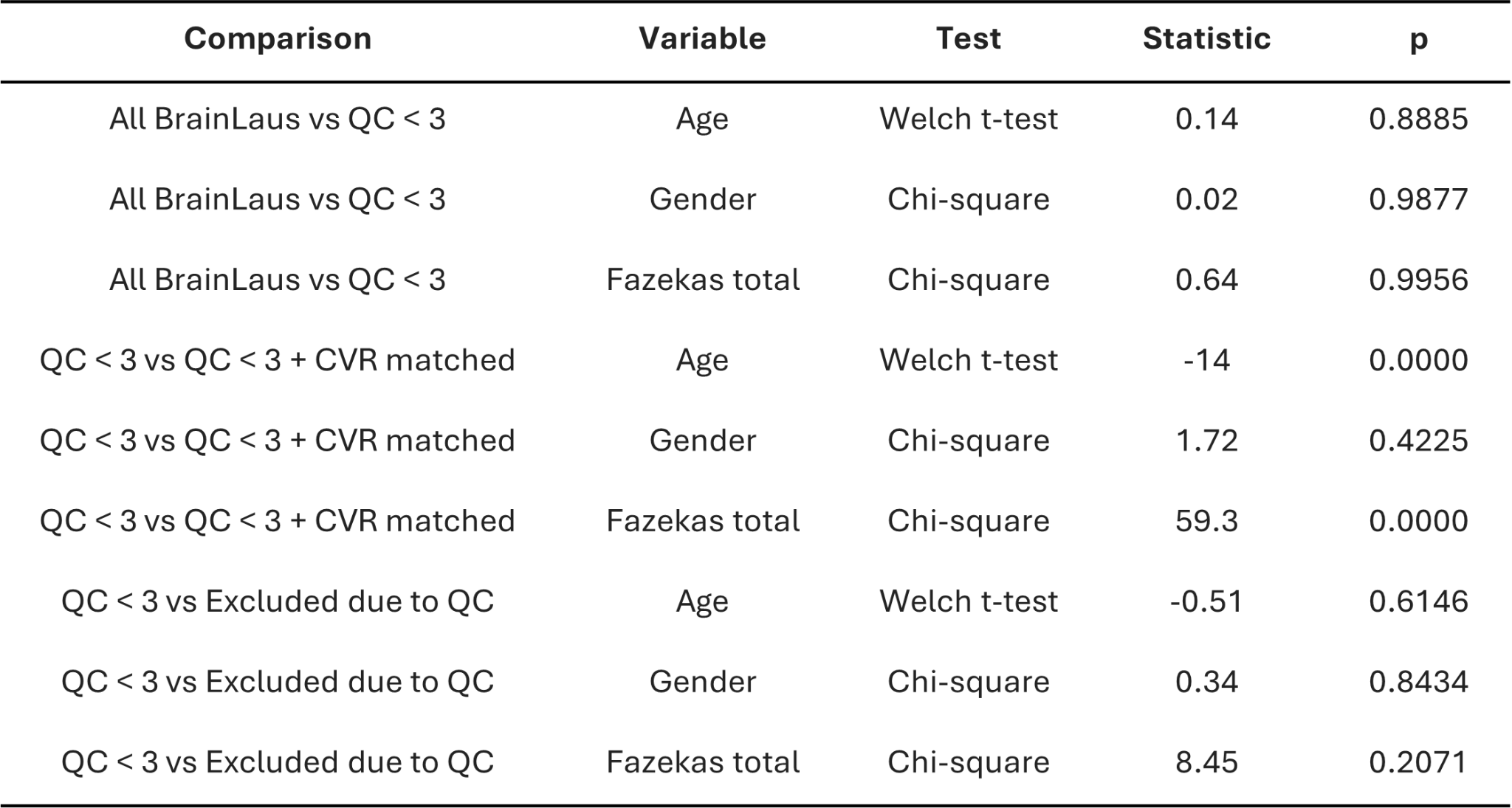

The visually-guided QC scoring procedure consisted of rating each contrast (FLAIR, R1, R2*, MT, and diffusion) on a scale from 1 to 4, with 0.5-point increments allowed.A score of 1 indicated excellent quality, whereas a score of 4 indicated poor quality. Scans with QC ratings up to 2.5 were included, while those with ratings equal to or above 3 were excluded. Quality control of the images was initially performed by a primary trained rater (AB) and subsequently validated by a neurologist (FC). As shown in **Supplementary Figure 4**, the 60 excluded scans were removed based on different combinations of failed QC criteria. Specifically, 18 scans were excluded due to poor R2* quality, 11 due to failure across all MPM-derived maps, 8 due to failure across all MPM maps and FLAIR, and the remaining scans due to various other combinations of QC failures. **Supplementary Figure 3** provides illustrative examples of excluded scans. Motion artifacts mainly affected the R2* maps and, to a lesser extent, the other MPM maps. Diffusion-weighted scans were discarded due to incorrect masking during preprocessing or improper FOV positioning during data acquisition. Additionally, two scans were excluded due to the presence of a meningioma, and a few others were removed because of acquisition artifacts in the MPM maps.

**Cardiovascular risk factor assessment**

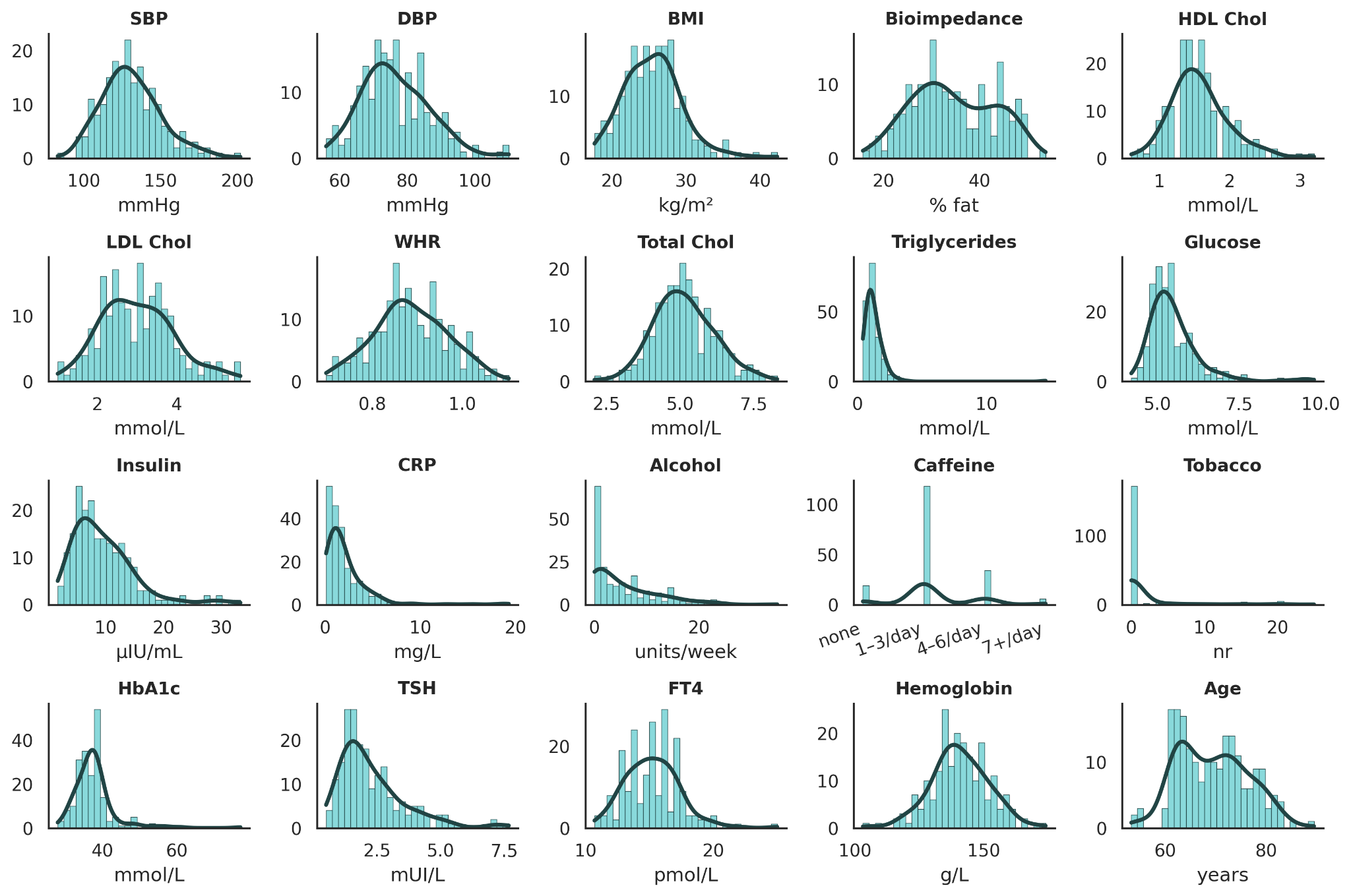

**Supplementary Figure 3 :** Histograms of frequency distributions of 20 original cardiovascular risk (CVR) variables, each annotated with its corresponding unit of measurement.

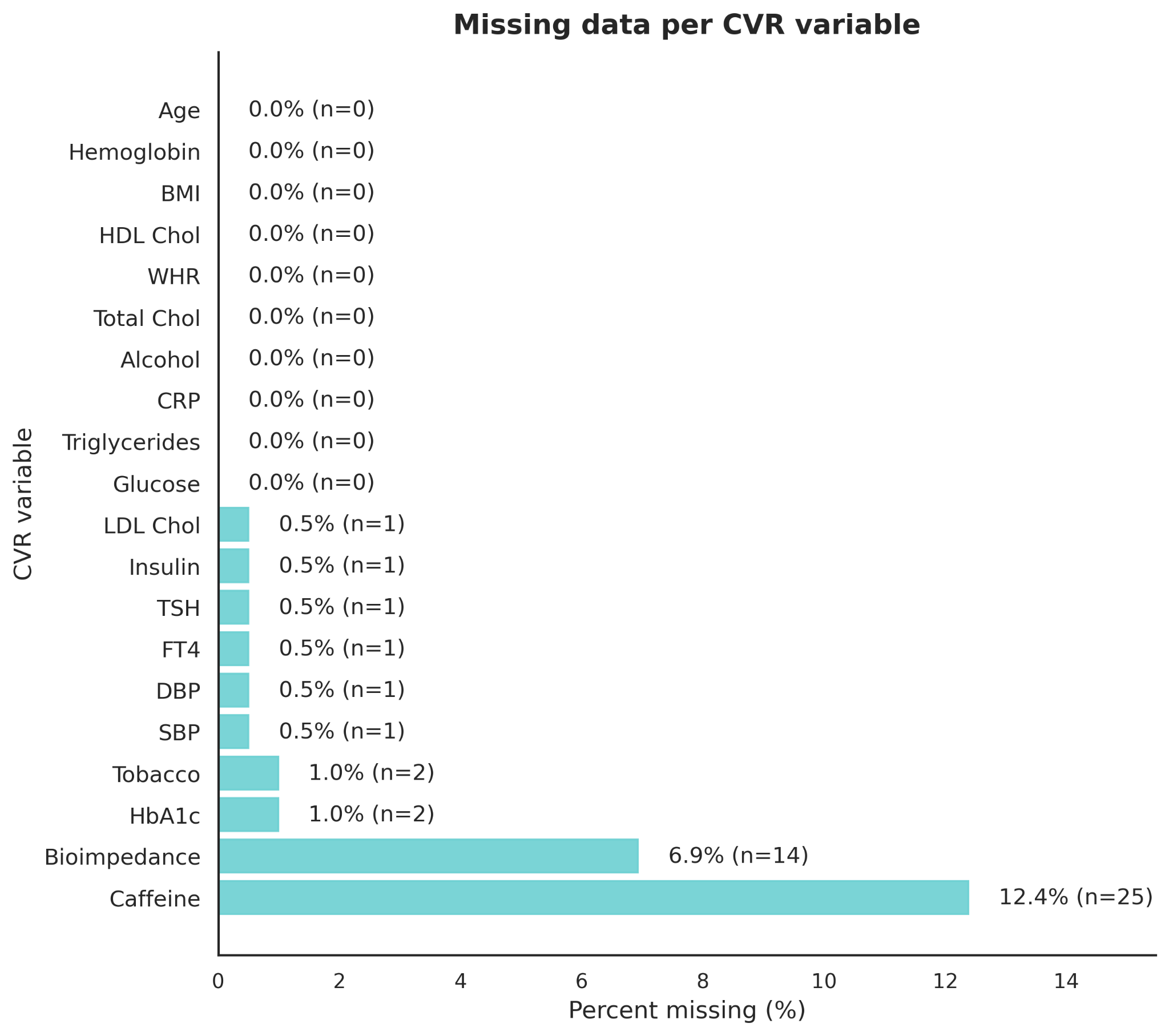

**Supplementary Figure 4:** Overview of missing CVR data among the 202 participants included in the PLS analysis. Both percentages and absolute counts (n) are shown.

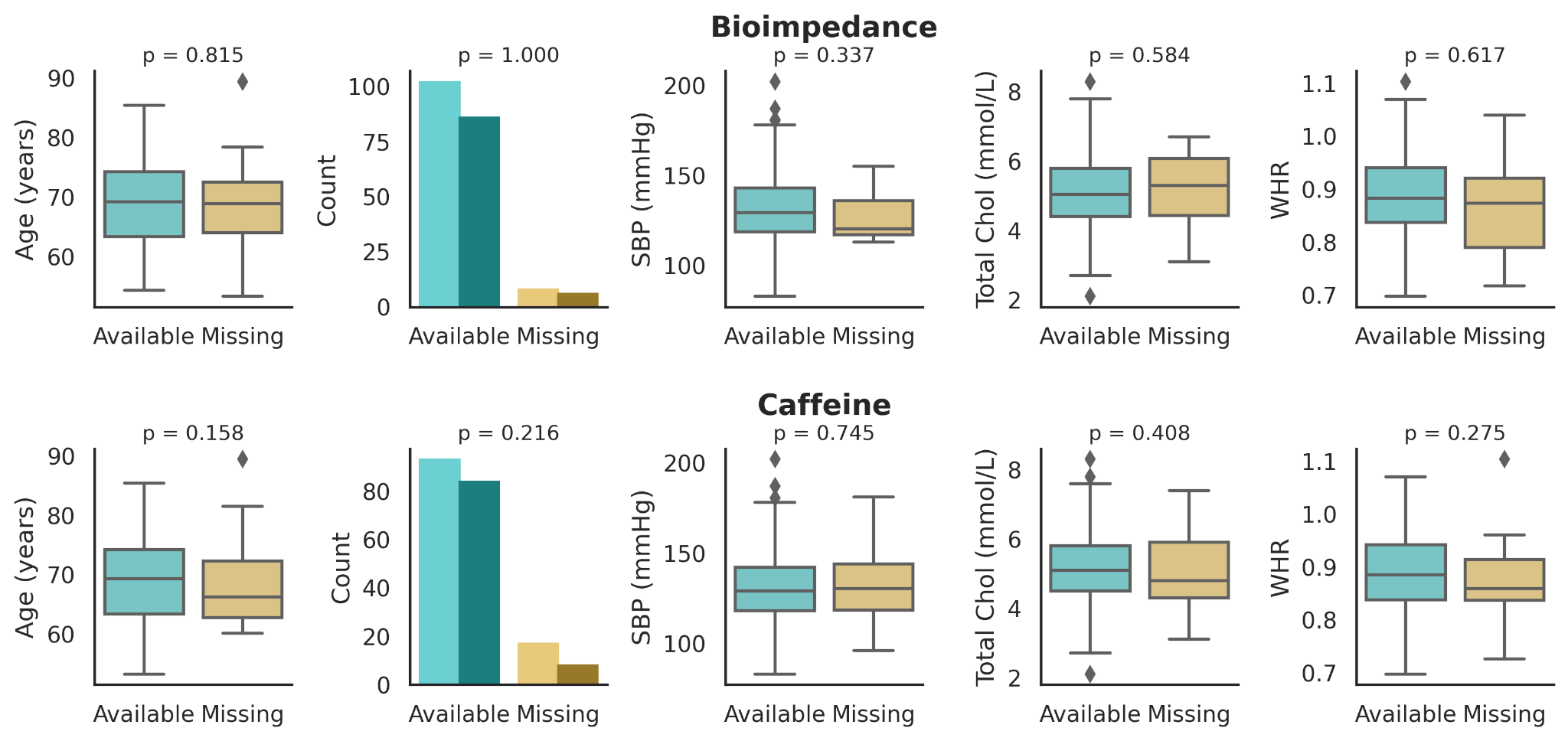

**Supplementary Figure 5:** Description of CVR variables with more than 1% missing data. Missingness is characterized across age, sex (light = females; dark = males), SBP, total cholesterol, and WHR. All p-values are non-significant (Mann–Whitney tests for continuous variables and Chi-square tests for binary variables comparing the distribution of sex between available and missing data).

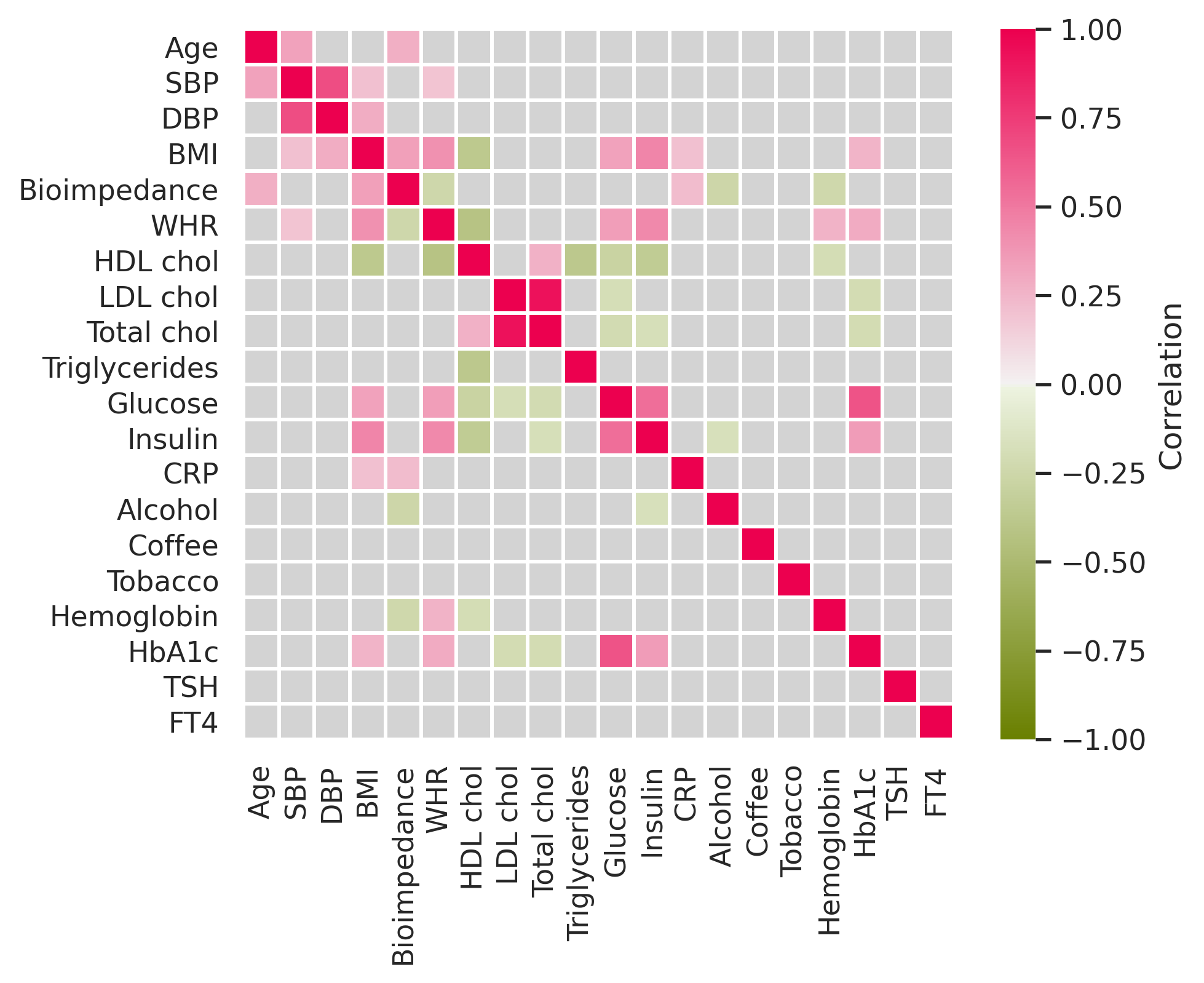

**Supplementary Figure 6**: Heatmap showing significant cross-correlations between the cardiovascular risk factors based on the imputed dataset. Pink color indicates significant positive correlations, green indicates significant negative correlations, and grey indicates non-significant correlations after false discovery rate (FDR) correction at p < 0.05. SBP: systolic blood pressure, DBP: diastolic blood pressure, BMI: body mass index, WHR: waist-to-hip ratio, HbA1c: glycated hemoglobin, CRP: high-sensitivity C-reactive protein, TSH: thyroid-stimulating hormone and FT4: free thyroxine.

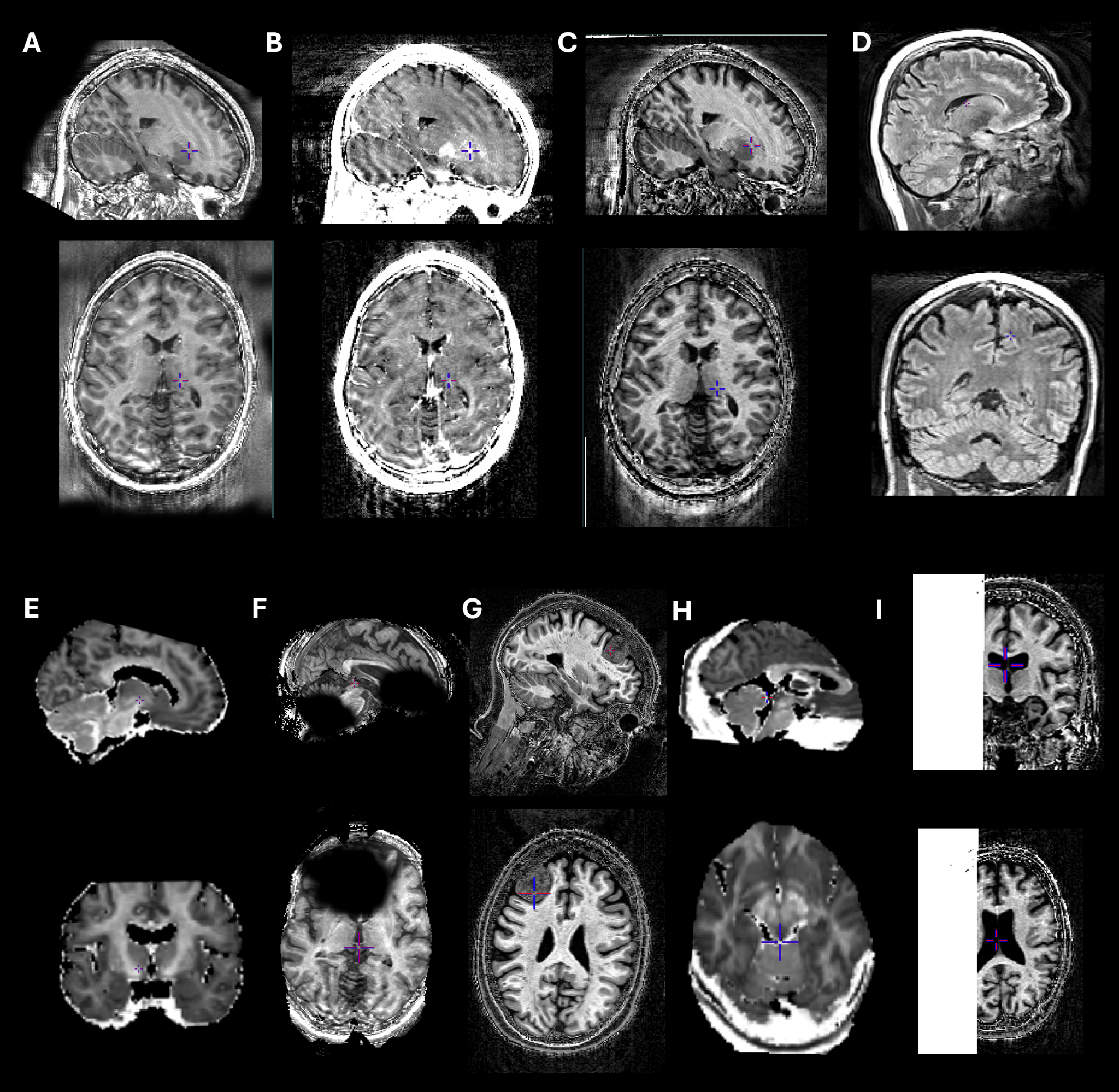

**Supplementary Figure 7:** Examples of excluded scans. (A–C) Motion “ringing” artifact in MPM maps; (D) FLAIR image excluded due to motion artifacts; (E) DWI-derived map excluded due to incorrect field-of-view; (F) MPM map excluded due to bias field artifacts; (G) dataset excluded due to presence of large meningioma; (H) DWI-derived map excluded due to incorrect masking; (I) MPM map excluded due to acquisition errors.

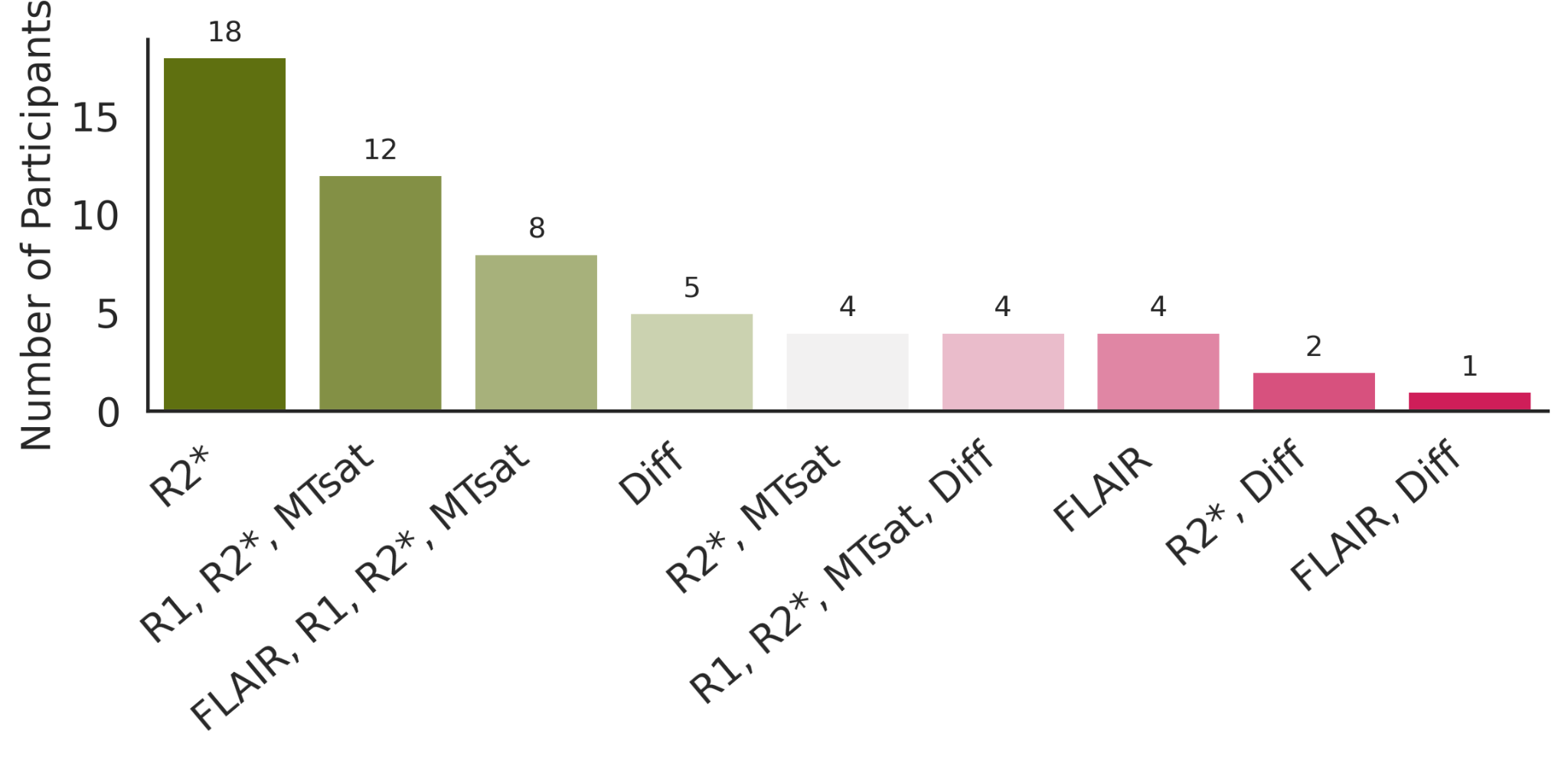

**Supplementary Figure 8:** Breakdown of the 60 excluded participants according to the combinations of QC failures.

**Imaging protocol**

We acquired three multi-echo FLASH T1- (TR/α = 18.7 ms/20°), PD- and MT-weighted contrasts (23.7 ms/6°) and TE between 2.2 ms and 19.7 ms (Draganski et al. 2011; Weiskopf et al. 2013). MT-weighting was accomplished by applying an off-resonance Gaussian-shaped RF pulse (4 ms duration, 220° nominal flip angle, 2 kHz frequency offset from water resonance) prior to excitation. For the T1- and MT-weighted acquisitions, multiple gradient echoes were obtained with alternating readout polarity at six equidistant TE ranging from 2.2 to 14.7 ms, and for the PD-weighted acquisition, echoes were captured at eight equidistant TE between 2.2 ms and 19.7 ms. Additional acquisition parameters included: 1 mm isotropic resolution, a matrix size of 256 × 240 × 176, parallel imaging with a GRAPPA factor of 2 in the phase-encoding direction, 6/8 partial Fourier in the partition direction, non-selective RF excitation, a readout bandwidth (BW) of 425 Hz/pixel, and an RF spoiling phase increment of 50°, resulting in a total acquisition time of approximately 19 minutes.

RF transmit field maps were generated and estimated using a 3D EPI scan that captured spin and stimulated echoes (SE and STE) with different refocusing flip angles (Lutti et al. 2010, 2012). The imaging parameters included a resolution of 4 mm isotropic, a matrix size of 64 × 48 × 48, and a FOV of 256 mm × 192 mm × 192 mm along the readout, phase encoding (PE), and partition directions. Parallel imaging was achieved with a GRAPPA factor of 2 × 2 in both the PE and partition directions. The sequence timings were set to TESE/TESTE/TM (mixing time)/TR = 37.06/37.06/31.2/500 ms, resulting in a total acquisition time of 3 minutes. The SE/STE refocusing pulse flip angles were reduced incrementally from 230°/115° to 130°/65° in steps of 10°/5°. The total acquisition time was 3 minutes.

**PD* exclusion**

PD* maps require rescaling of the mean white matter value to approximately 69 % to correct for receive bias field effects (Weiskopf et al. 2013). Given that this normalisation removes interindividual white matter variability, rescaled PD* maps no longer reflect true absolute proton density differences between participants and were therefore not included in our analyses.

**Layer-based extraction**

Layers were defined in absolute units (millimeters) using the geodesic distance implementation. We calculated a geodesic distance map within a volume restricted to NAWM. Voxels belonging to the WMH itself were excluded from the calculation, ensuring that geodesic layers could not extend inside the lesion or toward other lesions. Distance maps were then partitioned into 1-mm bands (0-1 mm, 1-2 mm, …, up to 5 mm), and each band was masked with NAWM, thereby constraining outer layers to NAWM tissue. For the lesion-wise approach, the geodesic distance transform was applied independently to each lesion. Each 1-mm distance band was again restricted to NAWM before binarisation, ensuring that outer shells around a given lesion could never include voxels from other lesions or non-WM tissue.

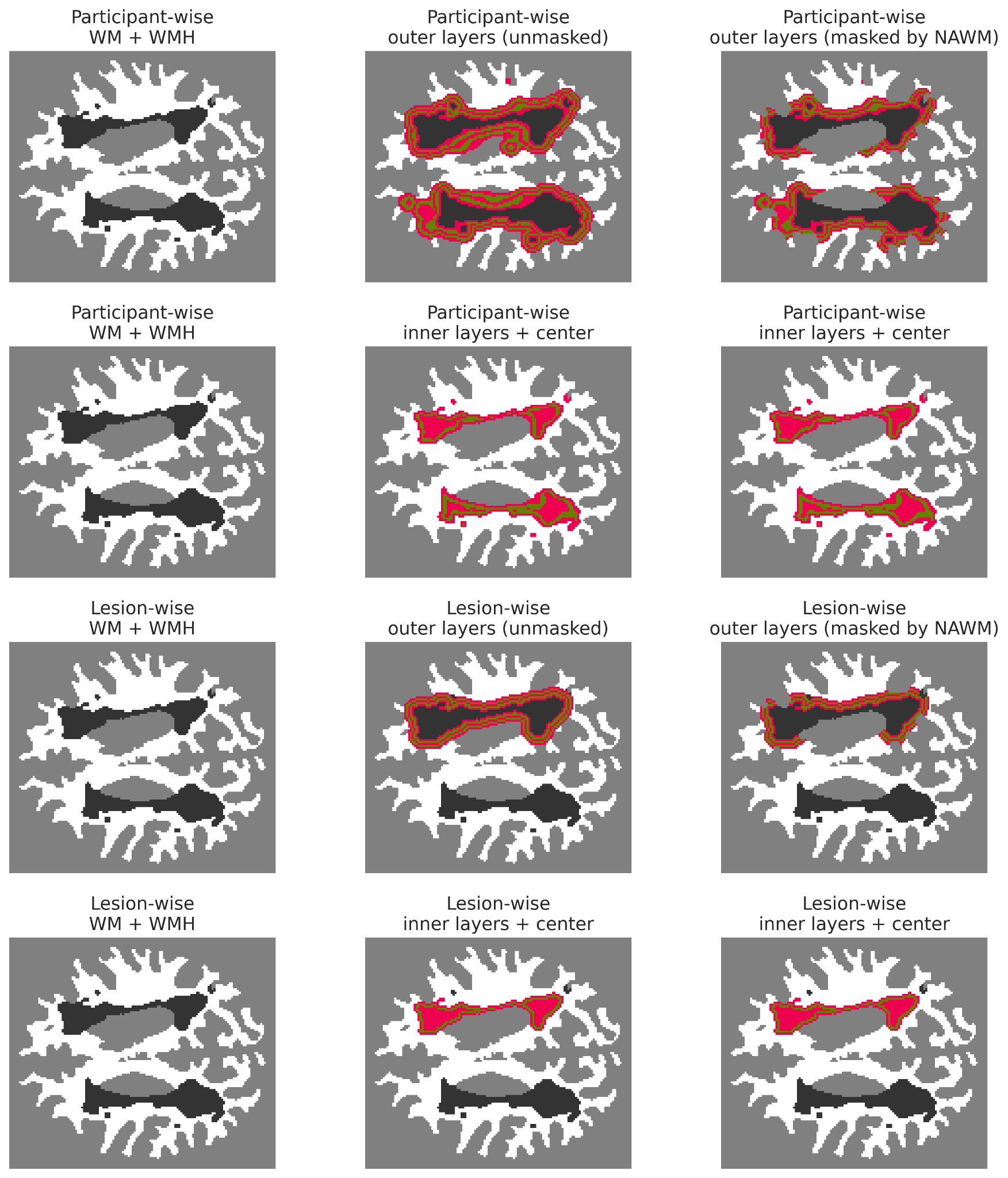

**Supplementary Figure 9:** Schematic of WMH layer construction and NAWM masking. **Row 1**: Participant-wise extraction of outer layers using the geodesic distance transform, before and after NAWM masking. **Row 2**: Participant-wise construction of inner layers using iterative erosion of the WMH mask. **Row 3**: Lesion-wise outer layer generation, where layers are defined independently for each lesion, illustrating potential discontinuities when neighbouring lesions are excluded. **Row 4:** Lesion-wise construction of inner layers through erosion. Colours denote successive 1-mm distance bands. NAWM is shown in white and WMH in dark grey.

This results in fewer subjects contributing to the innermost (“center”) layers. To address this issue, we revised our methodology so that layers are now extracted on a per lesion basis, rather than per subject. We quantified the number of lesions contributing to each layer (**Figure 6A**). As anticipated, many lesions disappear as morphological erosion progresses, yet approximately one-third of participants still contribute data points at the center layer (**Figure 6B**). We further examined which participants remain represented and found that younger individuals are disproportionately removed (**Figure 6C**), while those retained tend to have larger lesions (**Figure 6D**), consistent with the behaviour of the erosion process. We further examined how spatial change across layers varies as a function of lesion size (**Figure 6E**). While larger lesions expectedly show more pronounced layer-dependent changes, small lesions still follow the same gradual spatial trend, indicating that the phenomenon we report is not restricted to large lesions.

**Region-wise analyses**

Sampling of WMH and NAWM across regions (defined by lobe × WM compartment) resulted in 12 NAWM regions and a variable number of WMH samples per individual, depending on their lesion distribution. **Supplementary Table 2** makes explicit the composition of each analytical subset, allowing readers to see how inclusion patterns vary across regions as a direct consequence of the anatomical distribution of WMH. As shown in the demographic summaries, participants who contribute WMH data points in a given region tend to be older, whereas individuals without lesions in that region are younger. This pattern reflects the well-established age dependency of WMH burden rather than any methodological artifact.

**Supplementary Table 2:** Number of participants with non-missing NAWM and WMH measurements per region and per MRI-derived metric, together with the corresponding age and sex distributions for each subgroup.

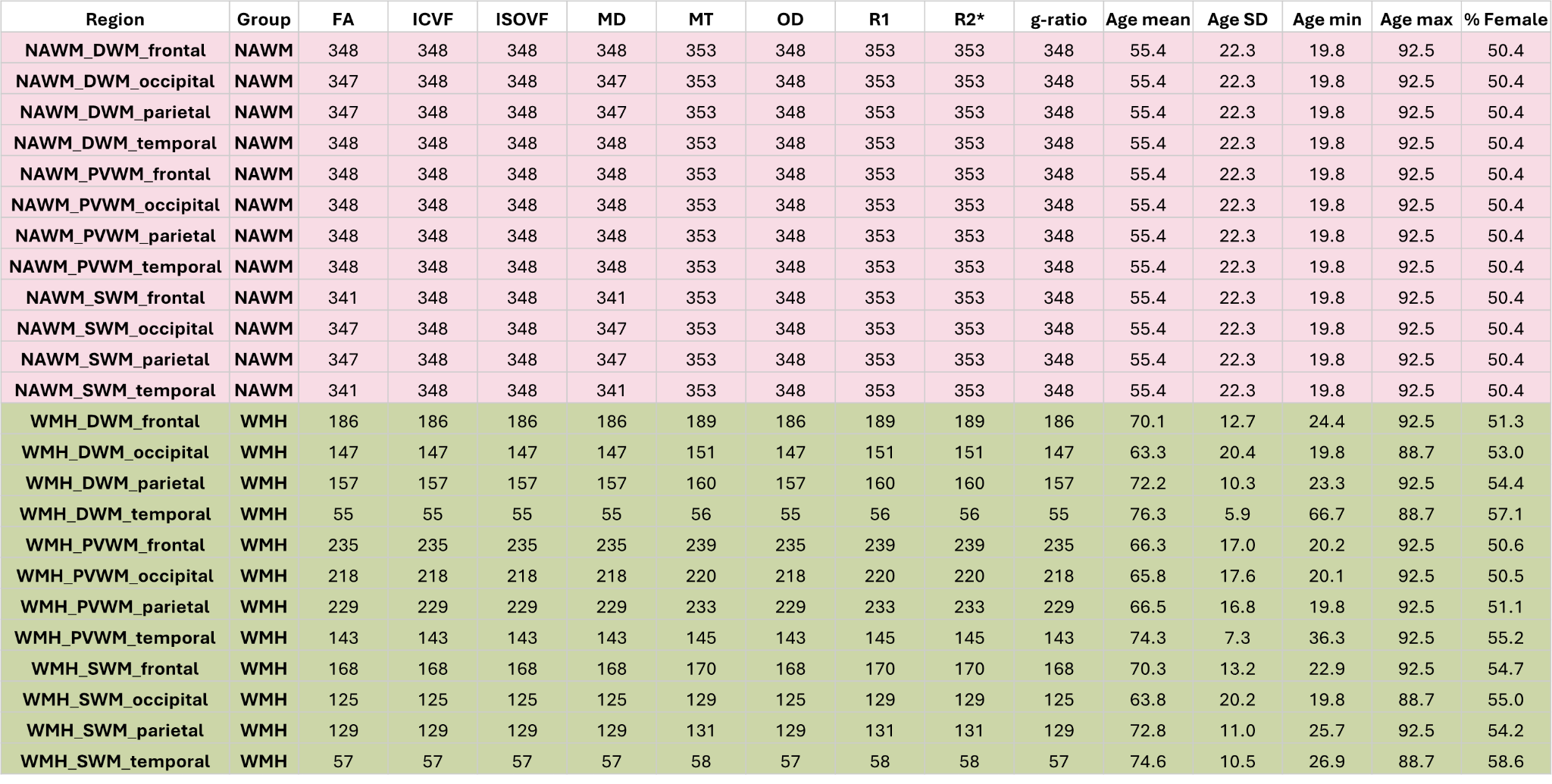

Two complementary analyses were provided in **Supplementary Figure 10**. In A, we visualised the age distribution of all participants who had WMH data in each region. It summarises the demographic composition of each subset transparently. The plots demonstrate that certain WMH patterns occur predominantly in older individuals. For example, temporal WMH were observed in the oldest participants, whereas frontal and occipital WMH were also present in early adulthood. This reflects known anatomical and age-related patterns of WMH burden rather than methodological choices.

In B, we quantified these differences statistically using a non-parametric Kruskal-Wallis test followed by FDR-corrected Dunn comparisons between all regions. The heatmap shows only significant results. Consistent with the KDE distributions, individuals with temporal WMHs are older than most other WMH carrying groups, and individuals with parietal DWM/SWM WMHs are older than individuals with frontal or occipital WMHs.

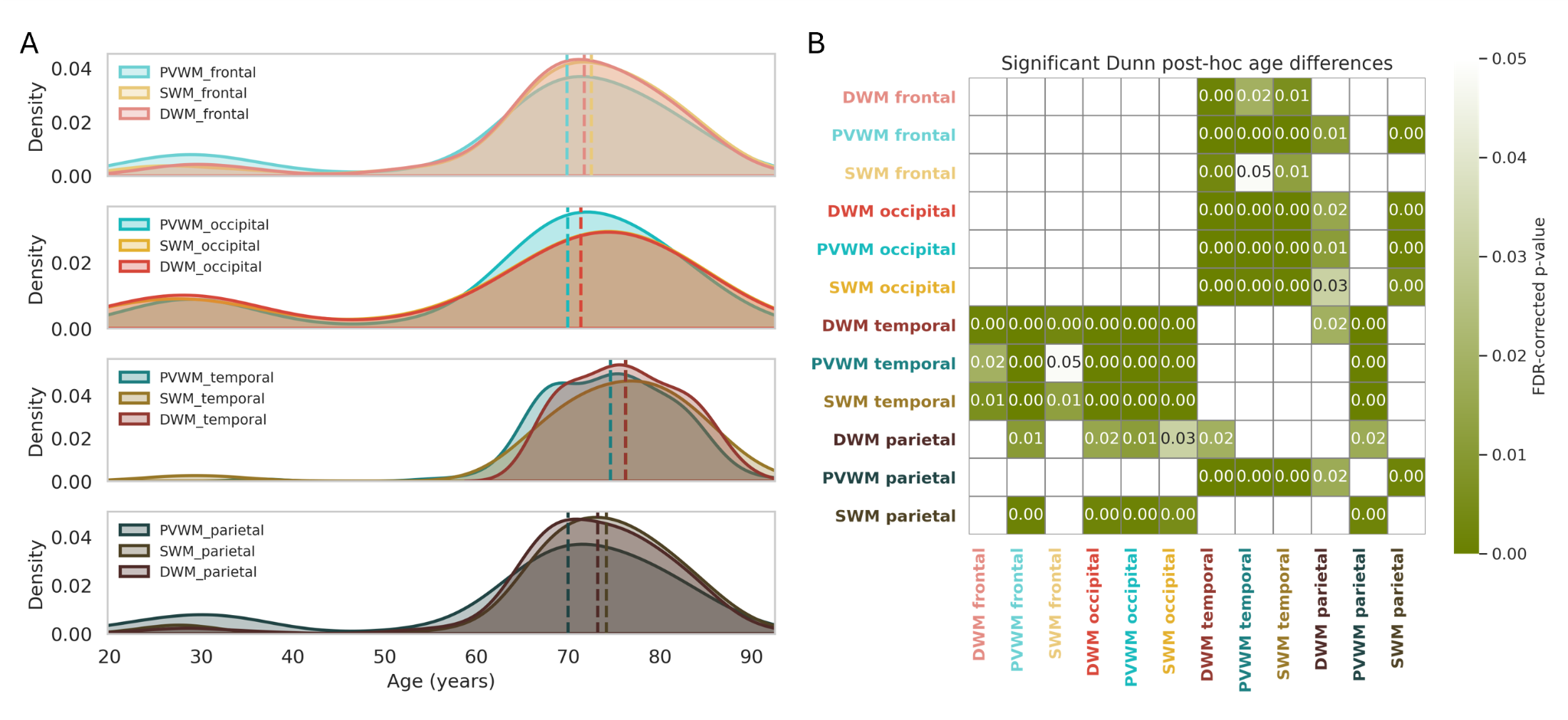

**Supplementary Figure 10:** A) Age distributions (kernel density estimates) of participants with non-missing WMH data in each region. Each curve represents the empirical age distribution of individuals contributing WMH data points for that region. B) Significant pairwise age differences between WMH-positive regional groups, assessed via Dunn post-hoc tests following a Kruskal–Wallis test after FDR correction for multiple comparisons. The heatmap displays only significant comparisons (p < 0.05, FDR-corrected). Darker colours indicate smaller p-values.

For transparency, we also replicated our analyses at coarser spatial scales, per lobe and per white matter compartment, to ensure that each subregion retained a larger and more comparable number of participants. **Supplementary Figure 11** presents the microstructural fingerprints for each WM compartment (n = 254 DWM, n = 301 PVWM, n = 236 SWM). The pattern is consistent with the main results: SWM exhibits substantially smaller WMH-NAWM differences than both DWM and PVWM, indicating that SWM is overall less affected, whereas periventricular and deep lesions show the strongest microstructural disruption.

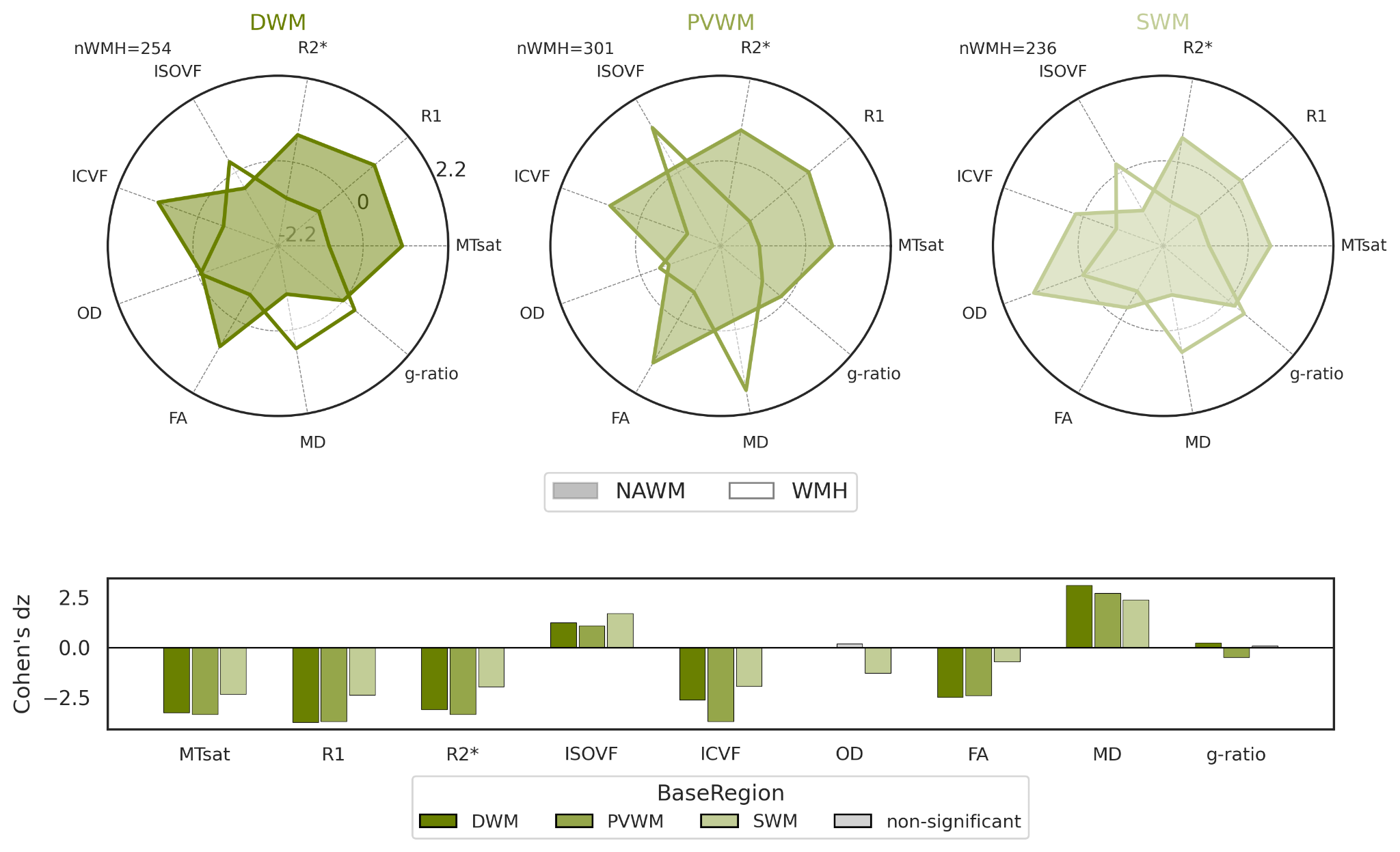

**Supplementary Figure 11 :** Fingerprint plots for white matter compartments - DWM, PVWM, SWM, accompanied by paired Cohen’s dz effect sizes for WMH versus NAWM. Effect sizes were computed using within-subject differences to quantify region-specific microstructural alterations.

We also performed complementary lobe-wise analyses (frontal: n = 255, occipital: n = 262, parietal: n = 239, temporal: n = 154). Across all four lobes, we observed significant paired WMH-NAWM differences for every microstructural metric. Among these, the frontal and occipital lobes consistently showed the largest effect sizes, indicating more pronounced WMH-related microstructural disruption in these regions. Taken together, the compartment- and lobe-focused results converge with our main regional findings: periventricular and deep white matter, particularly within the frontal and occipital lobes, exhibit the strongest microstructural differences associated with WMH. This cross-validated pattern reinforces the robustness of our interpretation regarding spatial heterogeneity in WMH pathology.

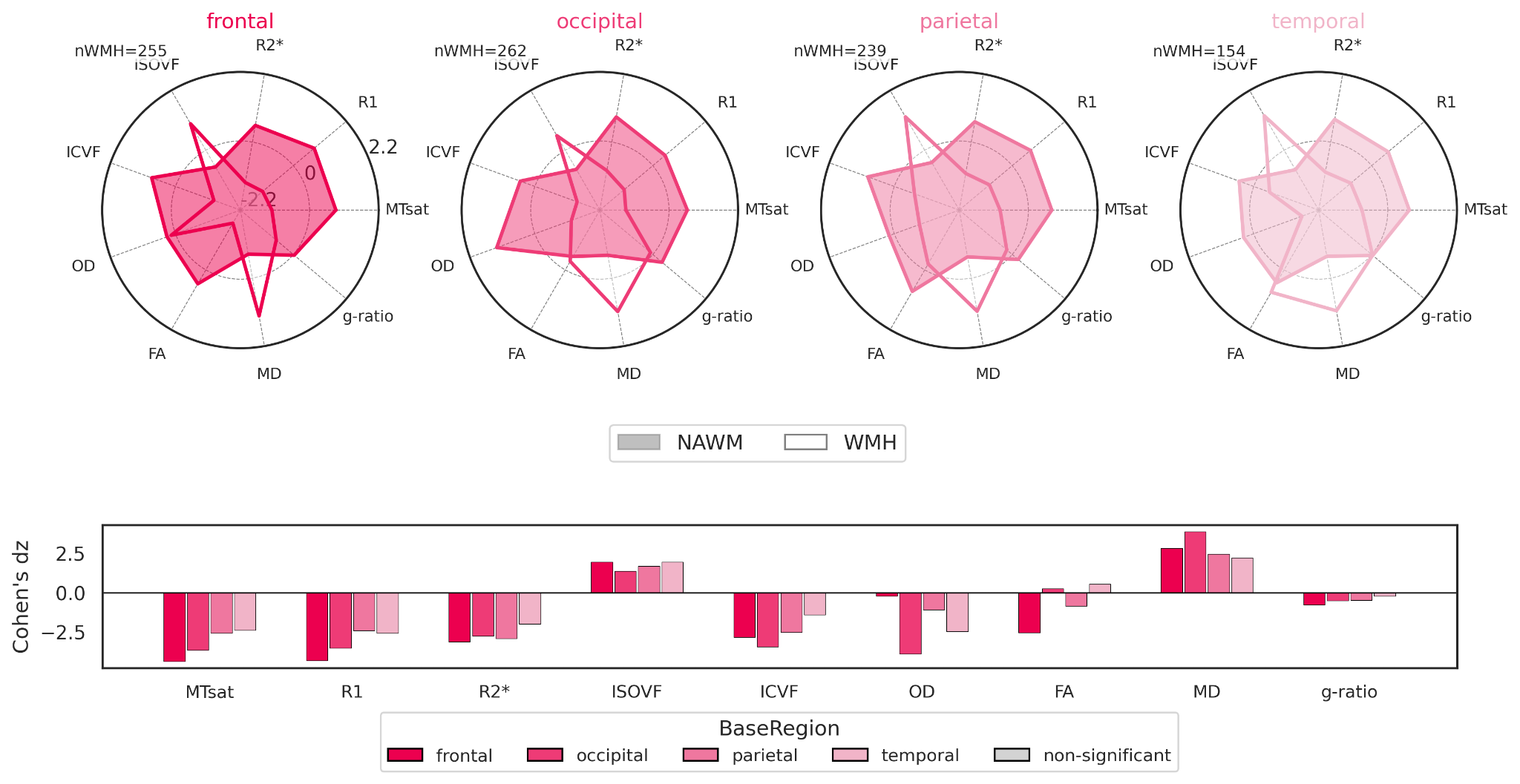

**Supplementary Figure 12** : Fingerprint plots for white matter adjacent to the frontal, occipital, parietal and temporal lobes accompanied by paired Cohen’s dz effect sizes for WMH versus NAWM. Effect sizes were computed using within-subject differences to quantify region-specific microstructural alterations.

**Principal component analyses**

Three complementary PCA analyses were performed in addition to the main model to verify that the principal findings were robust across different analytical specifications. In the NAWM-only PCA (**Supplementary Figure 13**), the first two principal components captured consistent microstructural gradients across anatomical regions, indicating that the structure of NAWM variation is stable and does not depend on WMH presence. In the WMH-only PCA (**Supplementary Figure 14**), PC1 similarly captured a dominant axis of WMH microstructural variation that was highly consistent with the WMH-NAWM separation observed in the combined PCA. Finally, the subject-level aggregated PCA (**Supplementary Figure 15**), where each participant contributes with only one NAWM and one WMH data point, showed that the primary axis of separation between WMH and NAWM remains well-distinguished even when weighting is equalised across individuals. Across all three analyses, the dominant components consistently distinguished NAWM from WMH and recovered similar patterns of between-metric covariance. This convergence indicates that the main PCA structure reflects genuine, reproducible microstructural differences rather than differential subject contribution or region-specific missingness.

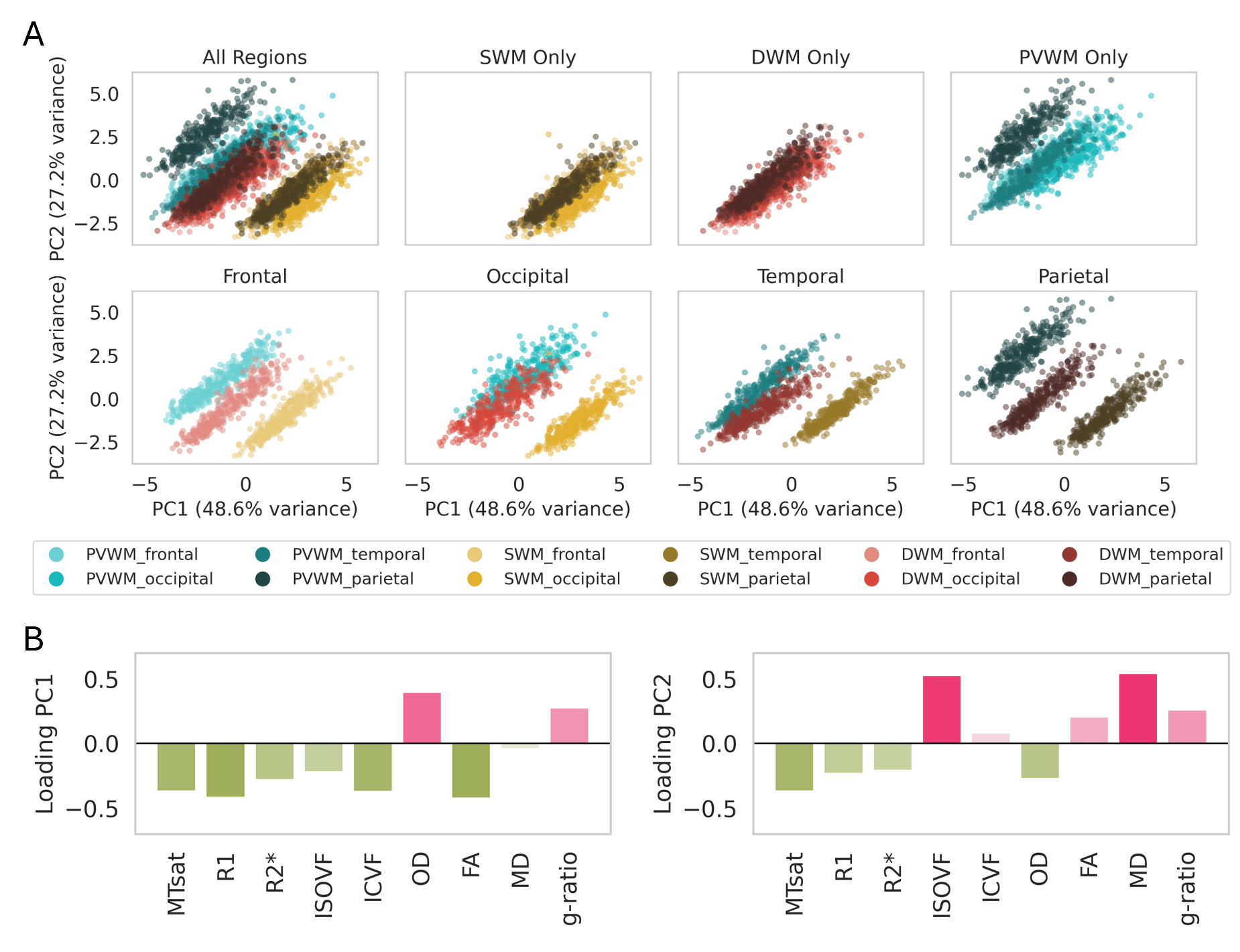

**Supplementary Figure 13:** Principal Component Analysis of all NAWM regions. Each point represents the NAWM microstructural profile of a given participant in a specific region. Colors indicate the underlying anatomical region. Axes show the first two principal components with their explained variance. All metrics were z-scored once across the entire NAWM dataset to ensure a common scaling prior to PCA. PC1 (48.60% variance explained) and PC2 (27.24%) were statistically significant in a permutation test (p = 0.0001).

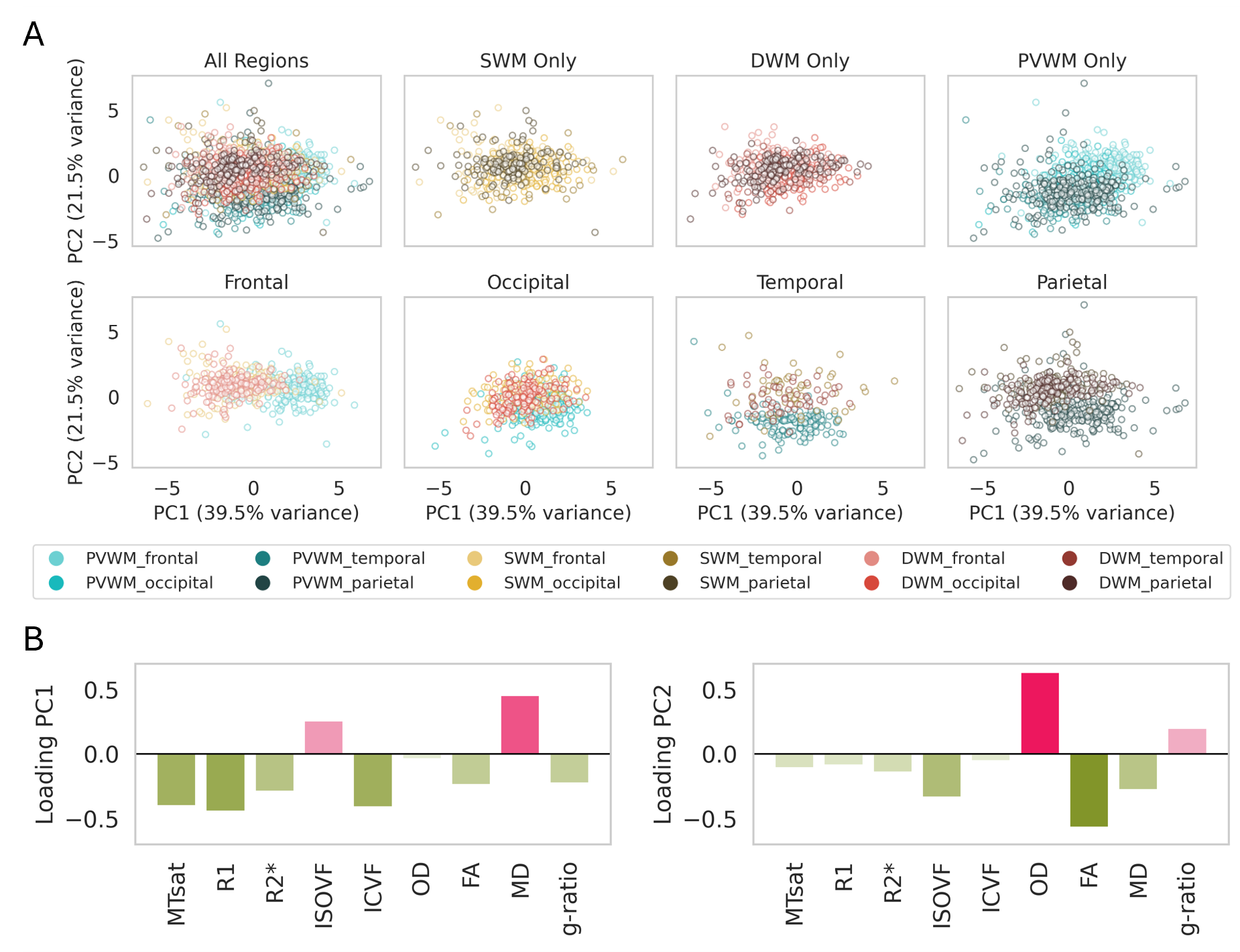

**Supplementary Figure 14:** Principal Component Analysis of WMH microstructure across all regions. Each point represents an individual participant's WMH profile in a specific region. Colors indicate anatomical regions.. Axes show the first two principal components with the explained variance. All metrics were z-scored once across the entire WMH dataset to ensure a common scaling prior to PCA. PC1 (39.54% variance explained) and PC2 (21.53%) were statistically significant in a permutation test (p = 0.0001).

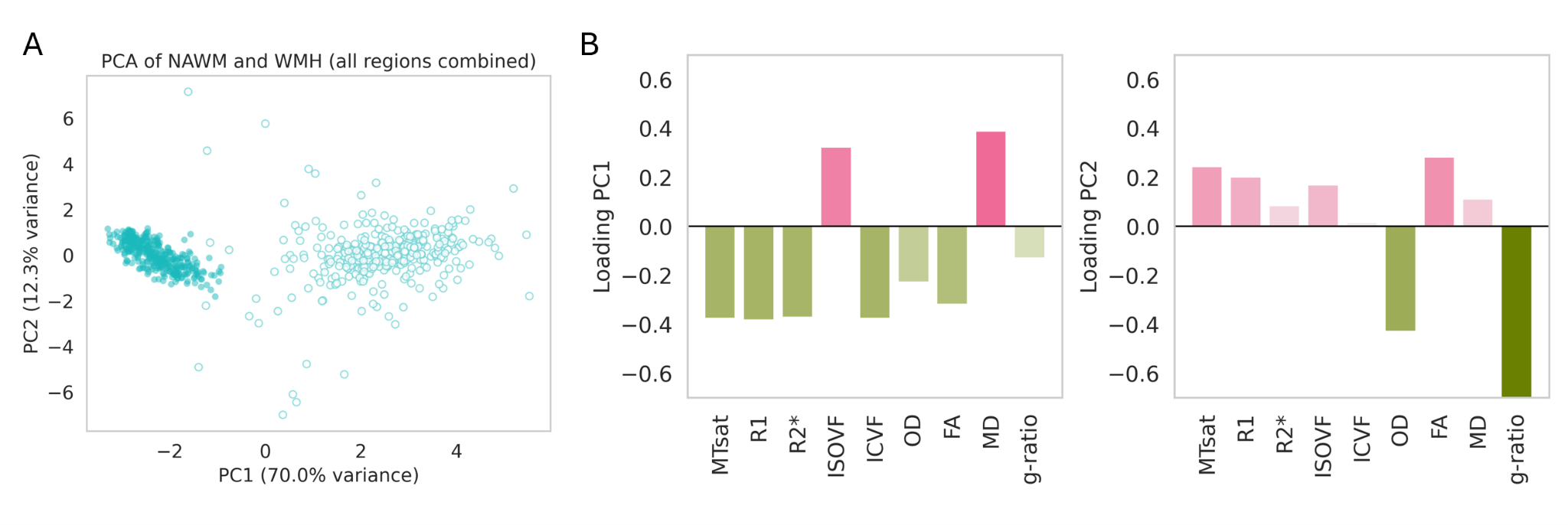

**Supplementary Figure 15**: Principal Component Analysis of WMH microstructure across all regions. Each point represents an individual participant's WMH profile in a specific region. Colors indicate anatomical regions. Axes show the first two principal components with their explained variance. All metrics were z-scored once across the entire WMH dataset to ensure a common scaling prior to PCA. PC1 (69.96% variance explained) was statistically significant in a permutation test (p = 0.0001), whereas PC2 (12.33%) was not (p = 0.67), indicating that only PC1 captures variance reliably above chance.

Then, we validated the stability of the PCA components using bootstrap resampling, permutation testing and split-half replications. First, bootstrap resampling of 1,000 iterations was performed. For each metric, the 95% bootstrap confidence intervals for the loadings were narrow, and the original loadings lay fully within these intervals. This demonstrates that the component structure is robust to resampling variability and not driven by a particular subset of subjects (**Supplementary Figure 16A**). We performed extensive permutation testing (10,000 permutations per component) in which metric values were randomly permuted across rows to disrupt covariance structure. The observed variance explained by PC1 and PC2 was significantly greater than expected under the null distribution (permutation p-values < 0.001), confirming that both components reflect statistically meaningful covariance patterns rather than chance structure (**Supplementary Figure 16B**). To further assess robustness, we conducted 1,000 split-half replications. In each iteration, participants were randomly divided into two equal halves and PCA was run independently in each half. The loading vectors from the two solutions were nearly identical (median loading correlation: PC1 = 0.9996; PC2 = 0.9990), indicating extremely high reproducibility of the component structure. This independently confirms that the PCA findings are not sample-specific (**Supplementary Figure 16C**).

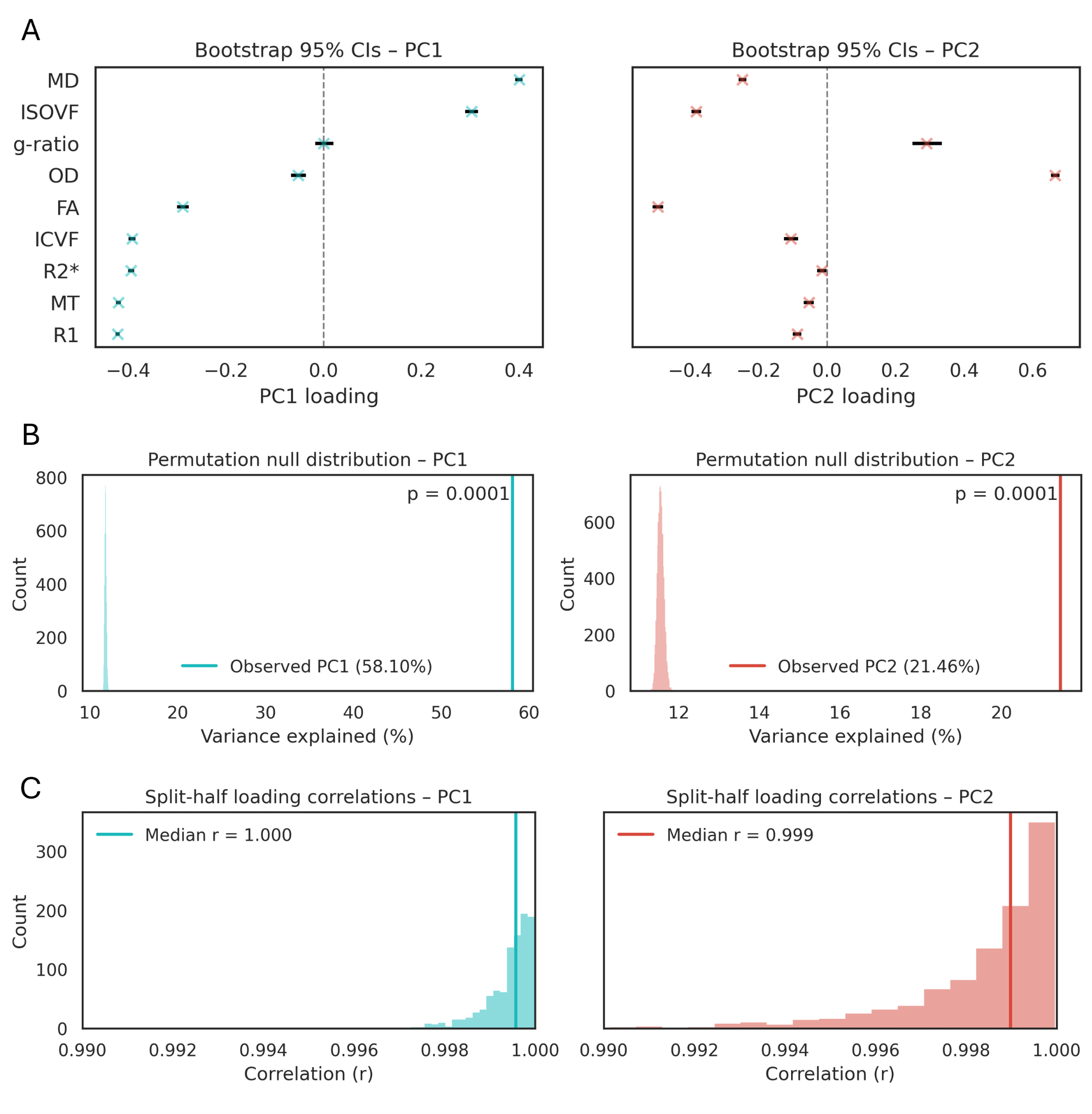

**Supplementary Figure 16**: **A)** Bootstrap resampling (1,000 iterations) confirming the stability of PCA loadings: for all MRI metrics, the 95% bootstrap confidence intervals were narrow, and the original loadings fell entirely within these intervals. **B)** Permutation testing (10,000 iterations) demonstrated that the variance explained by PC1 and PC2 was significantly greater than expected under the null (p < 0.001), indicating that both components capture meaningful covariance structure rather than random fluctuations. **C)** Split-half replication (1,000 iterations) showed excellent reproducibility of the component structure, with near-perfect correlations between loading vectors across halves (median r: PC1 = 0.9996; PC2 = 0.9990). Together, these analyses confirm that the PCA results are highly stable and not dependent on any particular subset of participants.

**Polynomial model selection**

To characterise spatial WMH profiles, polynomial models of increasing order were compared. Although higher-order models improved in-sample fit, cross-validated error showed an elbow at degrees 2-4, indicating limited added value beyond this range (**Supplementary Figures 17, 18** and **Supplementary Table 3**). Visually, curves were highly similar across model orders, capturing consistent non-linear patterns with only subtle differences. We therefore retained cubic models as a parsimonious and stable representation.

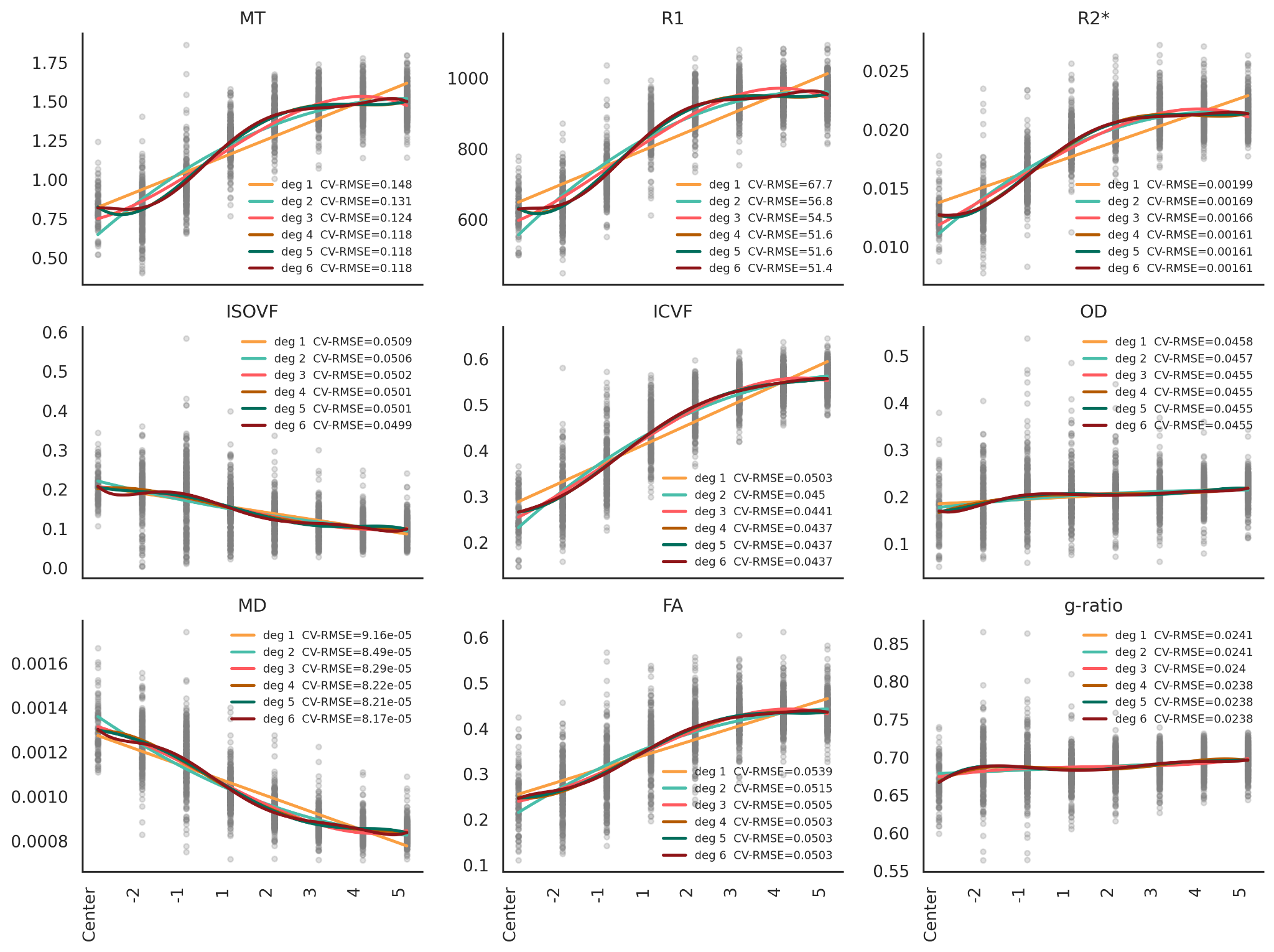

**Supplementary Figure 17:** Polynomial model comparison of participant-based microstructural gradients around WMH. Fits of degrees 1-6 are shown with cross-validated RMSE; gains in predictive accuracy diminish with increasing model complexity.

**Supplementary Table 3:** Summary of polynomial model selection for participant-wise microstructural gradients around WMH. For each metric, the optimal polynomial degree is reported based on in-sample goodness-of-fit (R²), penalised model selection criteria (AIC and BIC), and cross-validated performance using GroupKFold (elbow RMSE).

| Metric | Best R² | Best AIC | Best BIC | Elbow RMSE |
| --- | --- | --- | --- | --- |
| MTsat | 6 | 6 | 6 | 3 |
| R1 | 6 | 6 | 6 | 2 |
| R2* | 6 | 6 | 4 | 2 |
| ISOVF | 6 | 6 | 6 | 3 |
| ICVF | 6 | 4 | 4 | 2 |
| OD | 6 | 6 | 3 | 3 |
| MD | 6 | 6 | 6 | 3 |
| FA | 6 | 6 | 4 | 3 |
| g-ratio | 6 | 5 | 4 | 4 |

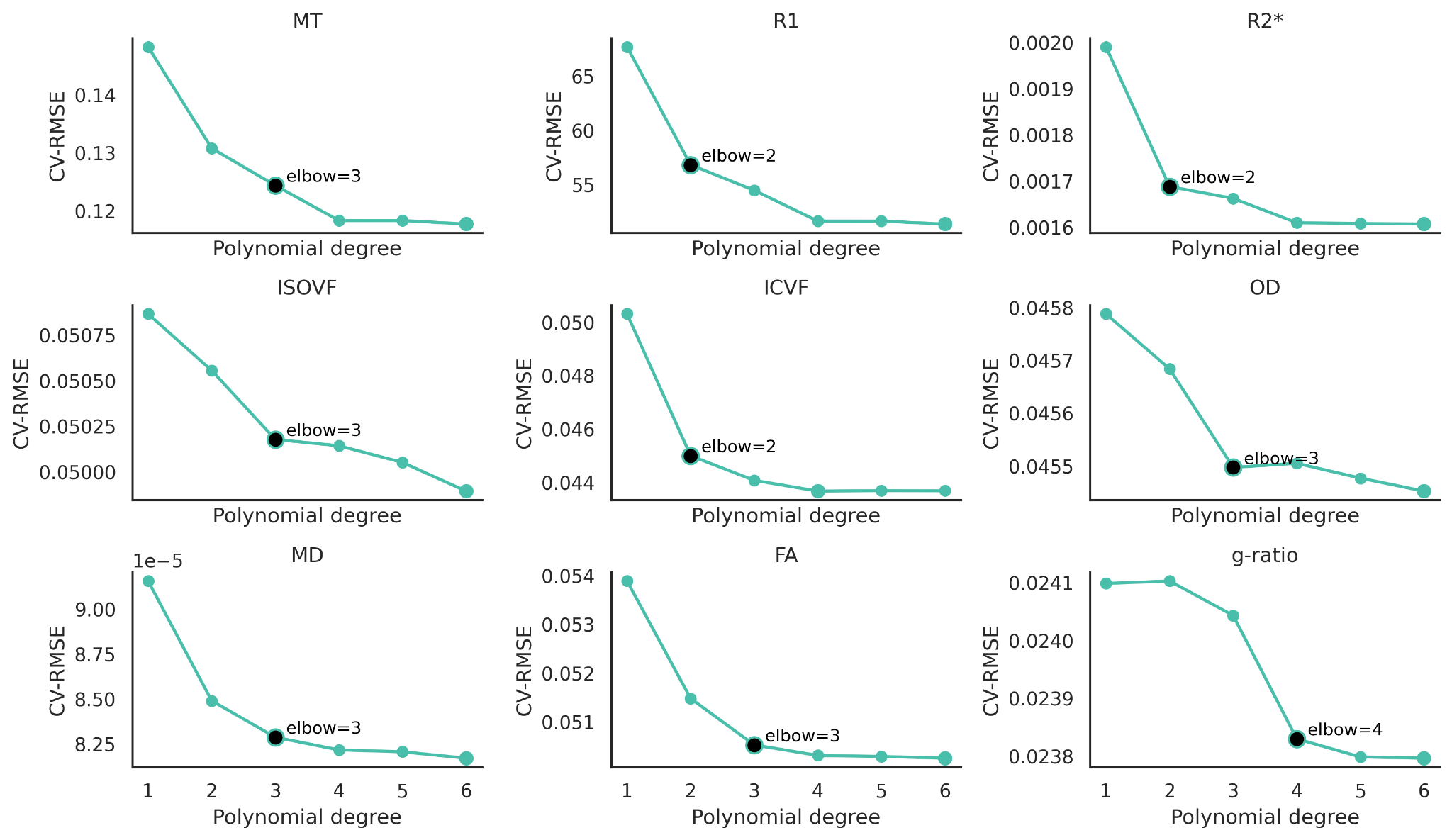

**Supplementary Figure 18:** Cross-validated model performance for participant-based microstructural gradients around WMH. Mean RMSE is shown as a function of polynomial degree (1-6) per metric: elbow points (black markers) indicate where improvements in predictive accuracy become marginal.

**Partial Least Squares split-half analysis.**

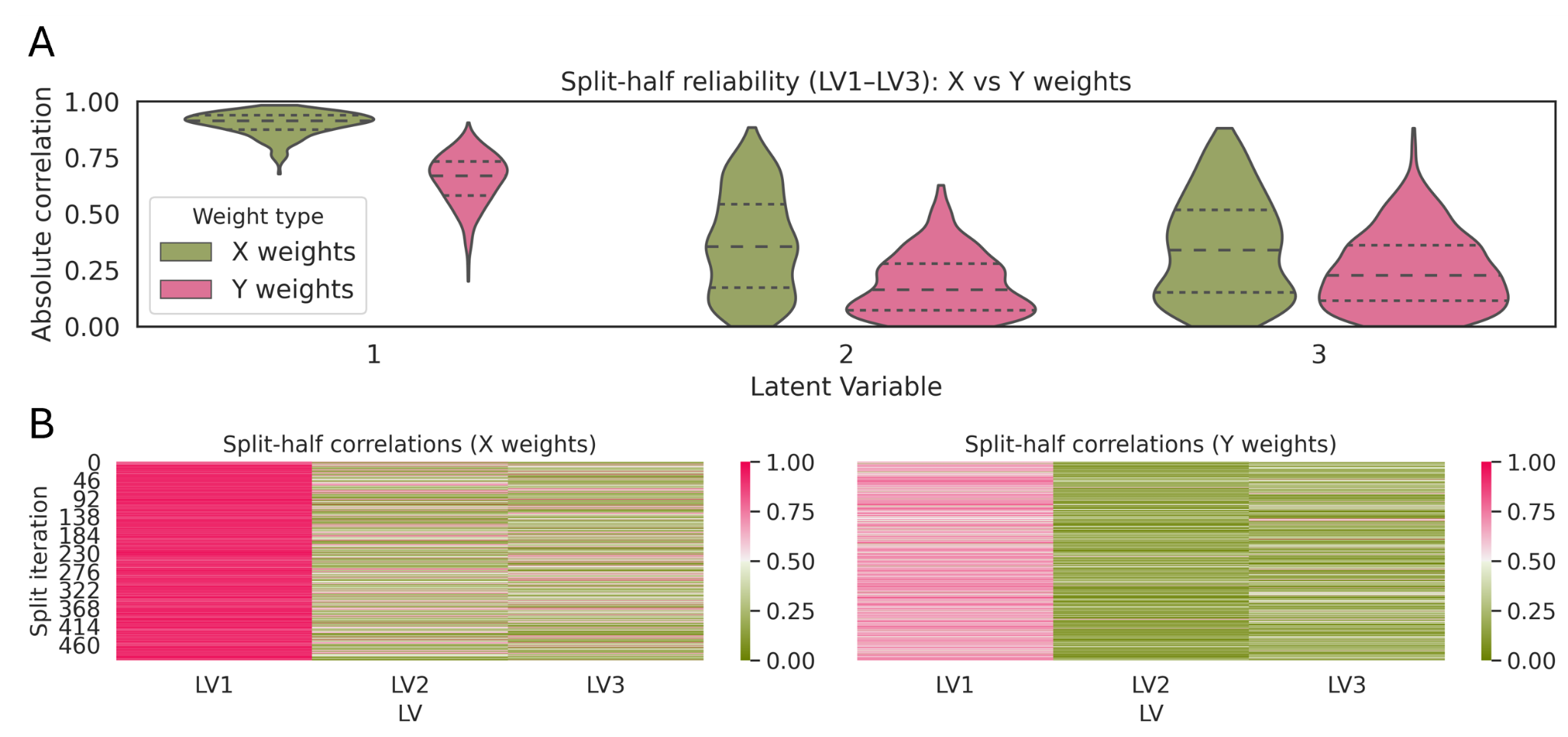

**Supplementary Figure 19**: Split-half reliability of PLS latent variables. Top: Violin plots of distributions of absolute split-half correlations for X (brain microstructure) and Y (cardiovascular / demographic) weights for the first three latent variables (LV1-LV3) across repeated random half-splits of the sample. Bottom: Heatmaps display the split-wise absolute correlations between weight vectors from the two halves for each LV, separately for X weights (left) and Y weights (right).

**Global effect between WMH and NAWM.**

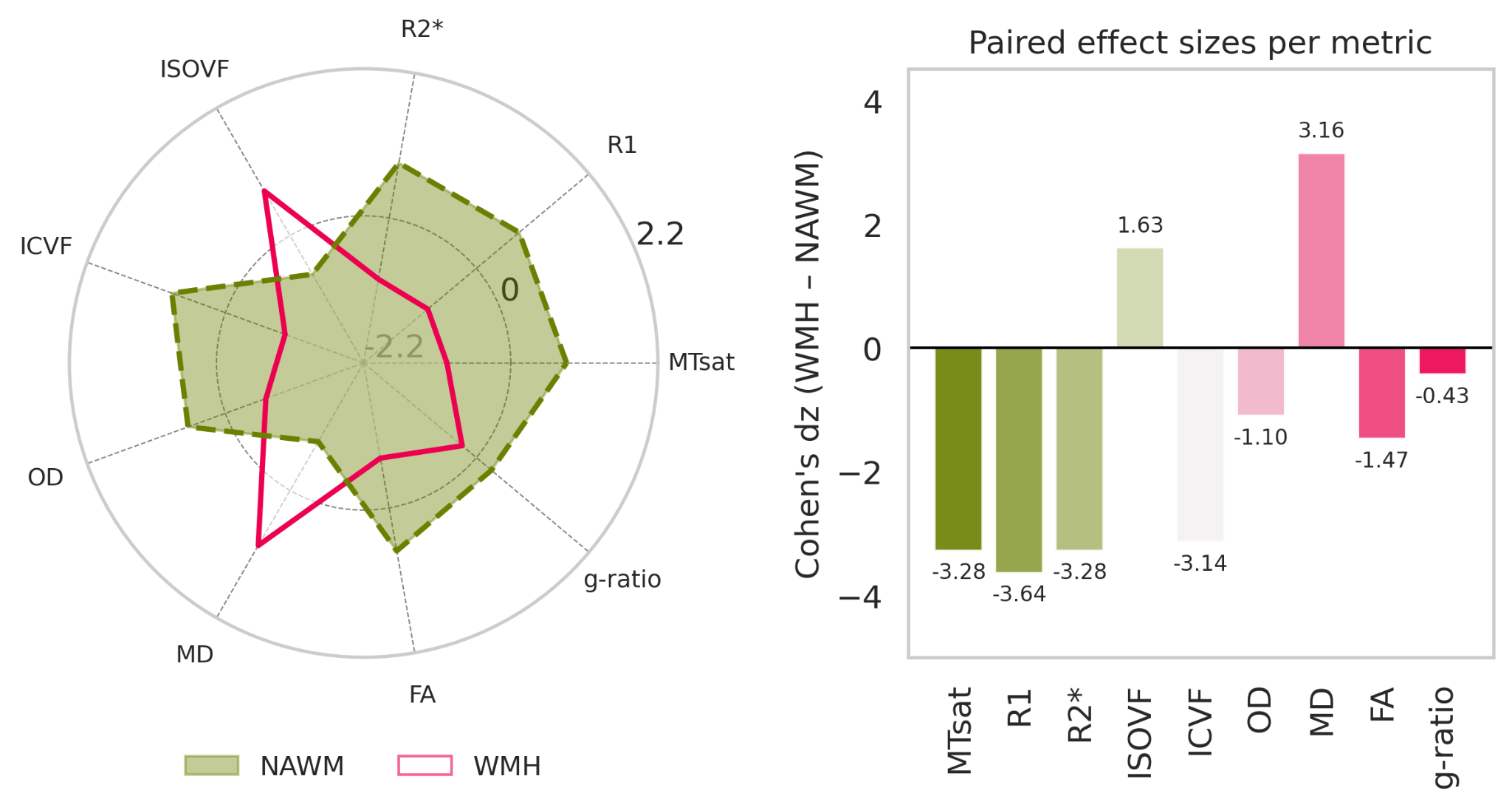

**Supplementary Figure 20:** Global comparison of WMH and NAWM microstructural profiles. Left: Radar plot showing the mean standardized value of each MRI microstructural metric in WMH and NAWM, computed within the same participants. Right: Paired effect sizes (Cohen’s dz) quantifying the magnitude of WMH-NAWM differences for each metric. Positive dz values indicate higher values in WMH relative to NAWM.

**Supplementary Table 4:** Summary of the strongest WMH-NAWM microstructural differences across regions. For each MRI metric, the table reports the minimum and maximum paired effect sizes (Cohen’s dz) observed across all lobar and depth-wise regions, along with the corresponding anatomical locations and FDR-corrected p-values. Cohen’s dz quantifies the within-subject WMH-NAWM contrast, where larger magnitudes indicate stronger and more consistent tissue alterations. This summary highlights which regions exhibit the weakest and strongest lesion-related microstructural disruptions for each metric. Pink shading indicates negative Cohen’s dz values (WMH < NAWM), whereas green shading indicates positive Cohen’s dz values (WMH > NAWM).

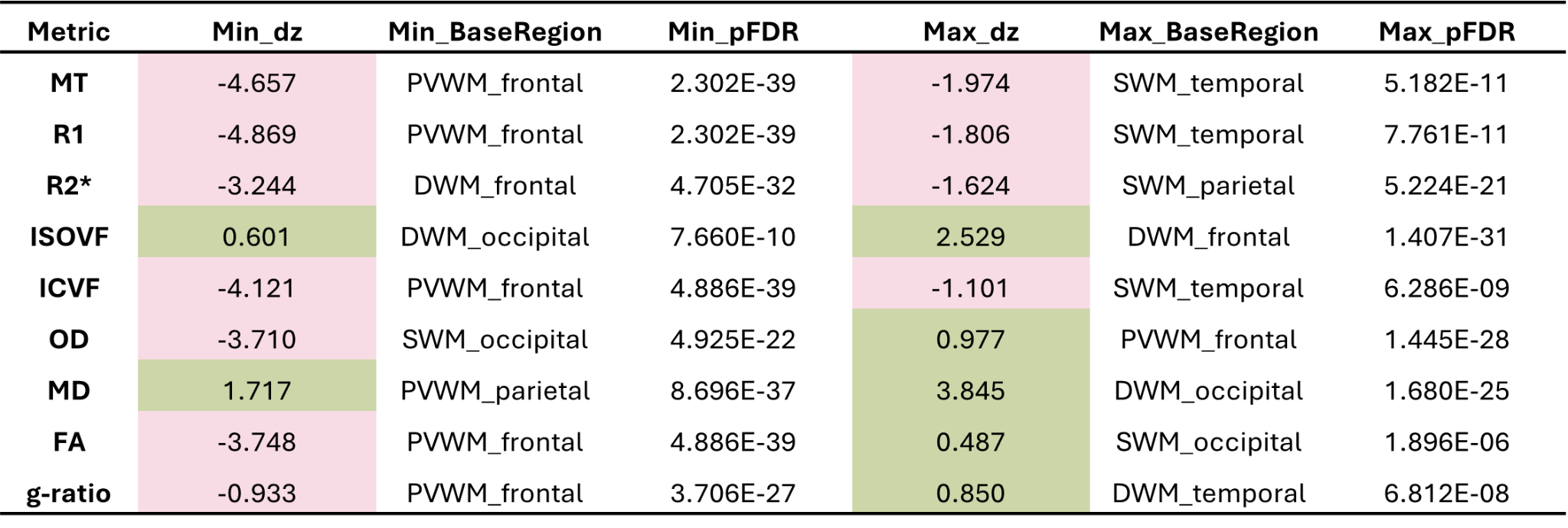

**Standardization and outlier detection investigation**

To investigate the effects of standardization and outlier detection in Figure 3, Plot WMH and NAWM metrics vs WMH or %WMH, we compared results from different models. We implemented a version of the analysis in which no z-scoring is applied to the modeled predictors or outcomes. In this version, the regressions are performed entirely on raw or log-transformed values, and standardized β coefficients are computed analytically from the raw slopes using the identity β_std = β_raw × (SD_x / SD_y). This approach is mathematically equivalent to fitting the model on standardized variables but avoids any explicit rescaling of the inputs. Importantly, the raw-scale regression results are virtually identical to the standardized ones: the direction, magnitude, and statistical significance of the associations are unchanged. This demonstrates that standardization played no substantive role in driving the observed effects and that our conclusions do not depend on the scaling choice.

For completeness, we also repeated all regressions without any outlier removal. The results remained identical, confirming that outlier trimming had little effect on the model, and did not change the interpretation. To ensure full transparency, we now include all three sets of results, standardized regression with trimming, raw regression on the same trimmed dataset, and raw regression without any trimming, in a **Supplementary Table 5**. Across all models, the effect sizes and p-values are consistent, further demonstrating that our conclusions are robust and not an artifact of repeated z-scoring, circular scaling, or outlier handling. Based on these results, we decided to show the results without trimming in the main paper to alter as little as we could the variability of our metrics.

**Supplementary Table 5:** Comparison of standardized and raw regression coefficients across modeling strategies. For each metric and each tissue class (WMH, NAWM), we calculated: 1) Z-standardized regression with outlier trimming, 2) Raw-scale OLS on the same trimmed dataset, 3) Raw-scale OLS with no outlier trimming.

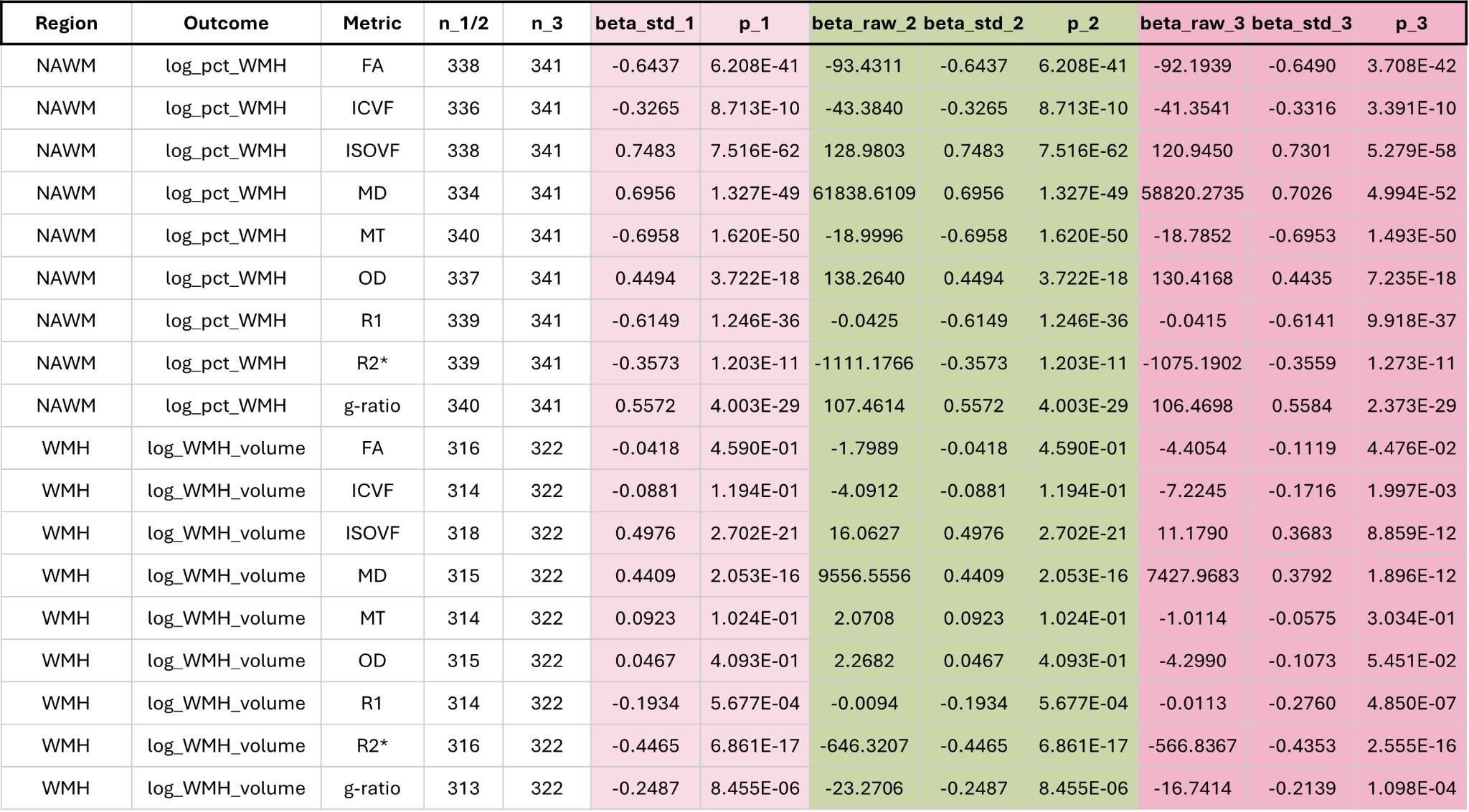

**PC relationships with age, tissue class, sex and log WMH volume**

We examined how PC1 (reflecting degeneration) and PC2 (reflecting tissue geometry) relate to age, sex, and WMH burden. CVR could not be included because the principal components were derived from the full dataset of 363 participants, for whom CVR measurements were not uniformly available. Using linear mixed-effects models with a random intercept per participant to account for multiple regional observations, we tested associations of PC scores with age, tissue class (WMH versus NAWM), sex, and log-transformed WMH volume, including an interaction between age and tissue class (see **Supplementary Table 6**). Sex was not significantly related to either PC1 or PC2. In contrast, age showed strong positive associations with both components, indicating that degeneration- and geometry-related patterns increase with advancing age. WMH voxels had substantially higher PC1 loadings than NAWM and moderately higher PC2 loadings, showing that PC1 in particular captures pathology characteristic of WMH. The age-by-tissue interaction was strongly negative for both components, demonstrating that age-related increases in PC scores were less strong in WMH compared with NAWM, likely because WMH voxels already lie at the extreme end of these latent dimensions. Total WMH volume was positively associated with PC1, indicating that individuals with greater WMH burden show stronger degeneration-related loading overall, whereas WMH volume was not related to PC2. In summary, degeneration-related variation (PC1) is strongly shaped by age, WMH tissue class, and total WMH burden, while geometry-related variation (PC2) increases with age and is higher in WMH but shows no dependence on WMH volume, providing a clearer interpretation of the tissue-specific patterns captured by the principal components.

**Supplementary Table 6**: Results of linear mixed-effects models predicting PC1 and PC2 from age, tissue type (normal-appearing white matter, NAWM, vs white matter hyperintensities, WMH), their interaction, sex (male vs female), and log-transformed WMH voxel count. Models include a random intercept for participant (PR) to account for multiple regional observations per subject. For each predictor and principal component, the table reports fixed-effect estimates (β), standard errors, z-values, raw p-values, and Benjamini–Hochberg FDR–adjusted p-values, with significance defined at FDR < 0.05.

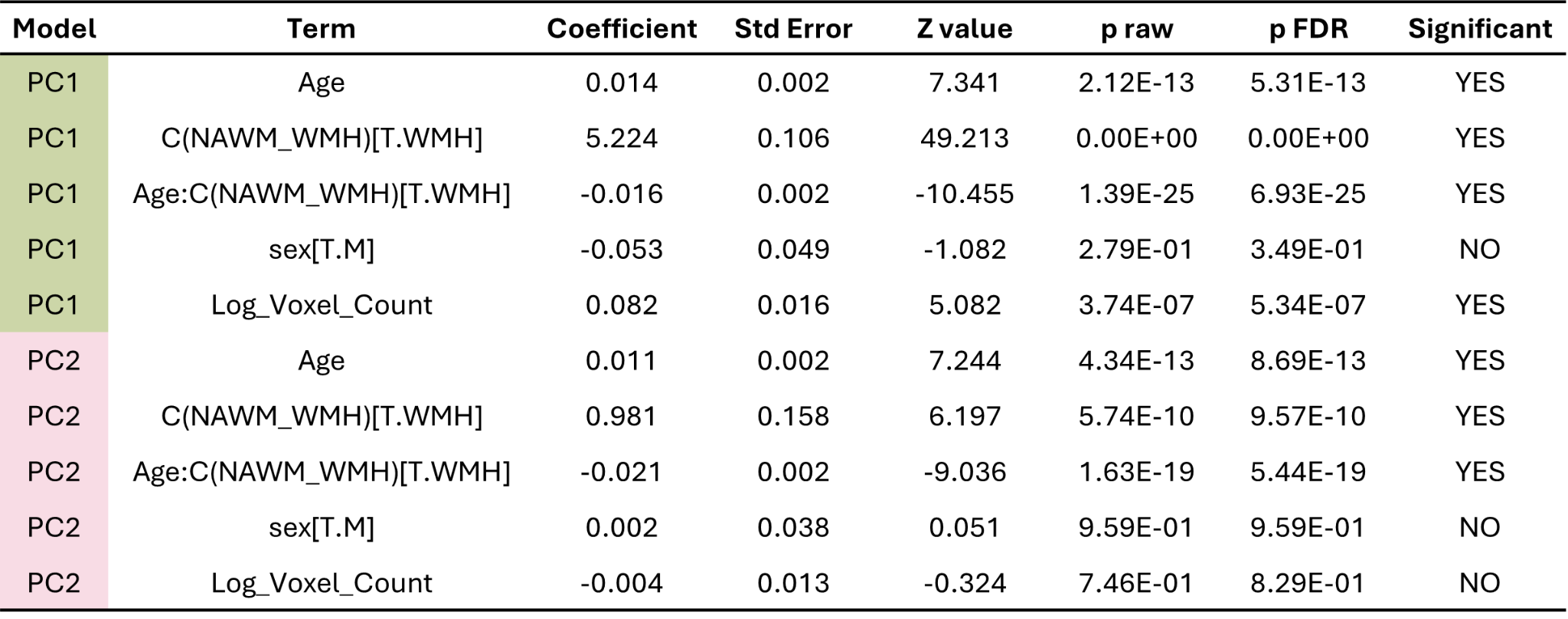

**Association between microstructural gradients and lesion size**

**Supplementary Table 7**: P-values for the cubic lesion-volume × layer interaction term for each microstructural metric. All interactions remained highly significant (FDR-corrected p < 0.001).

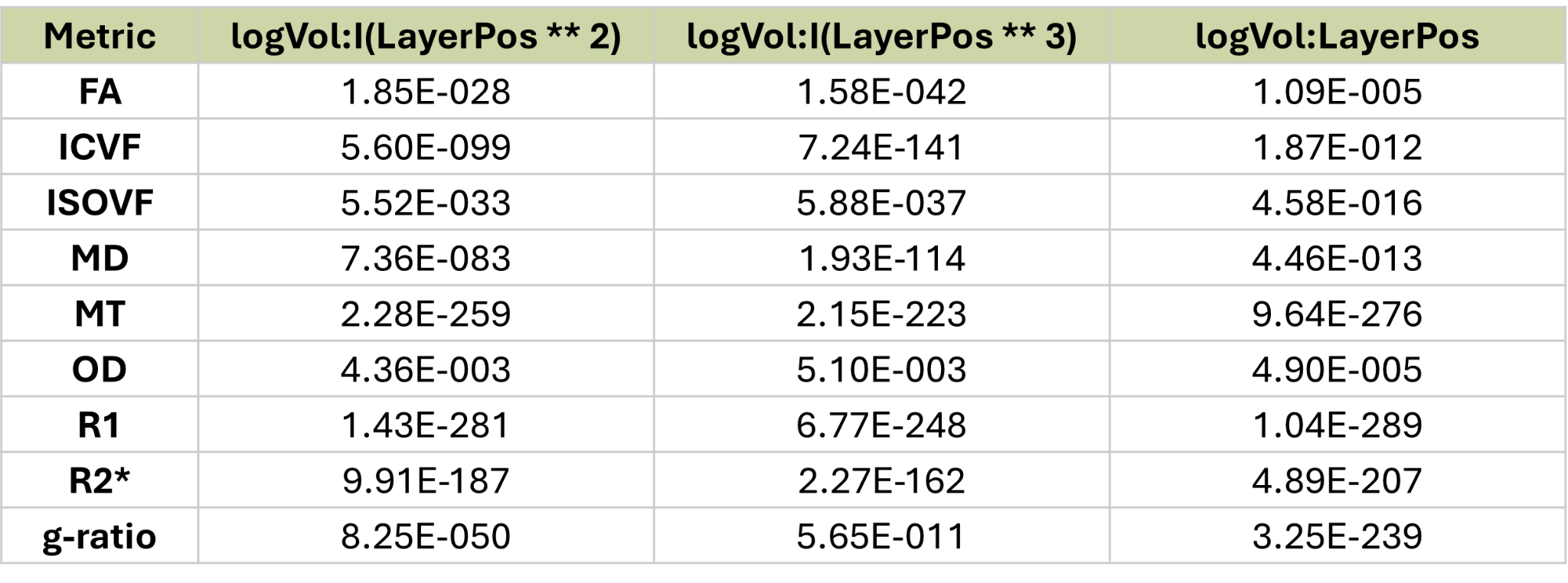

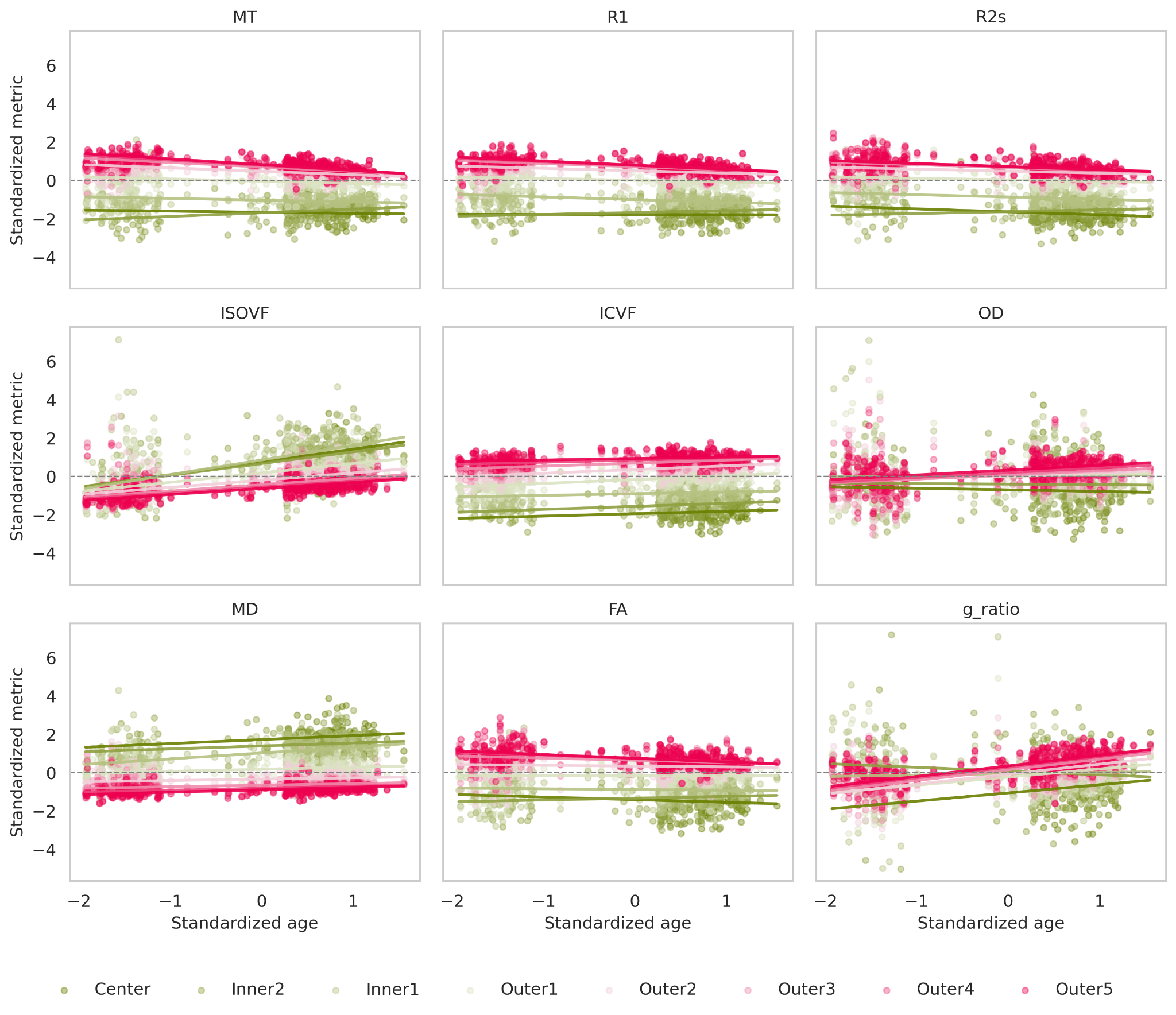

**Supplementary Figure 21:** Scatterplots and model fits of relationships between standardized age and standardized tissue microstructural metrics across WMH layers. Each panel displays a different MRI metric. Points correspond to individual subject–layer observations, colored by WMH distance (Center to Outer5). Lines represent predicted values from the mixed-effects model *metric_z ~ Age_z × Layer + logVol_z* with random intercepts and random age slopes, plotted at mean WMH volume. The figure illustrates layer-dependent age effects, where slope magnitude and direction vary systematically across modalities and spatial distance from WMH.

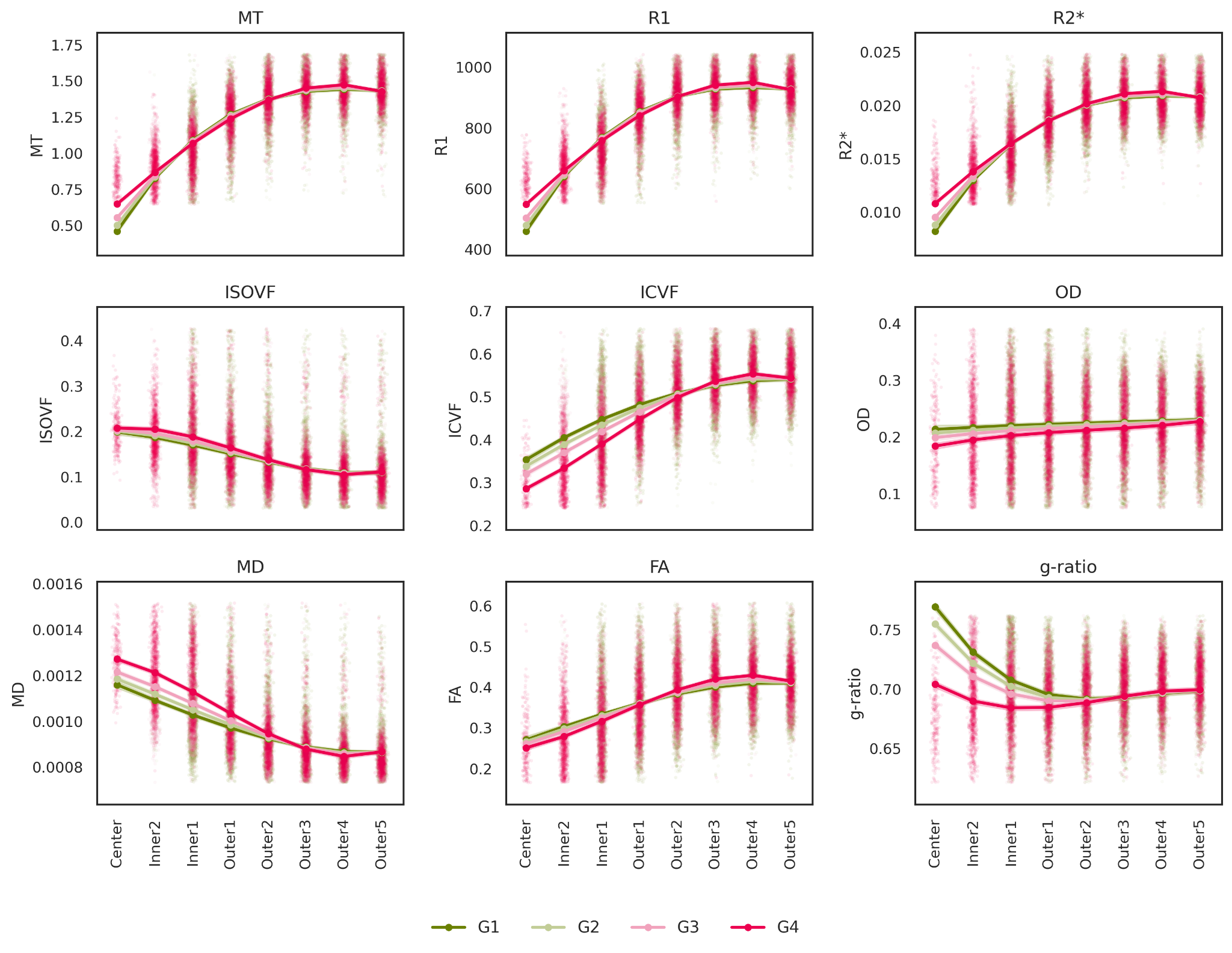

**Supplementary Figure 22:** Unsmoothed depth profiles from WMH centre to surrounding NAWM, stratified by lesion size. Mixed-effects model estimates (points and lines) are shown at discrete layer positions with 95% confidence intervals (shaded bands); raw data points are overlaid excluding the extreme 1% tails for visualisation purposes.

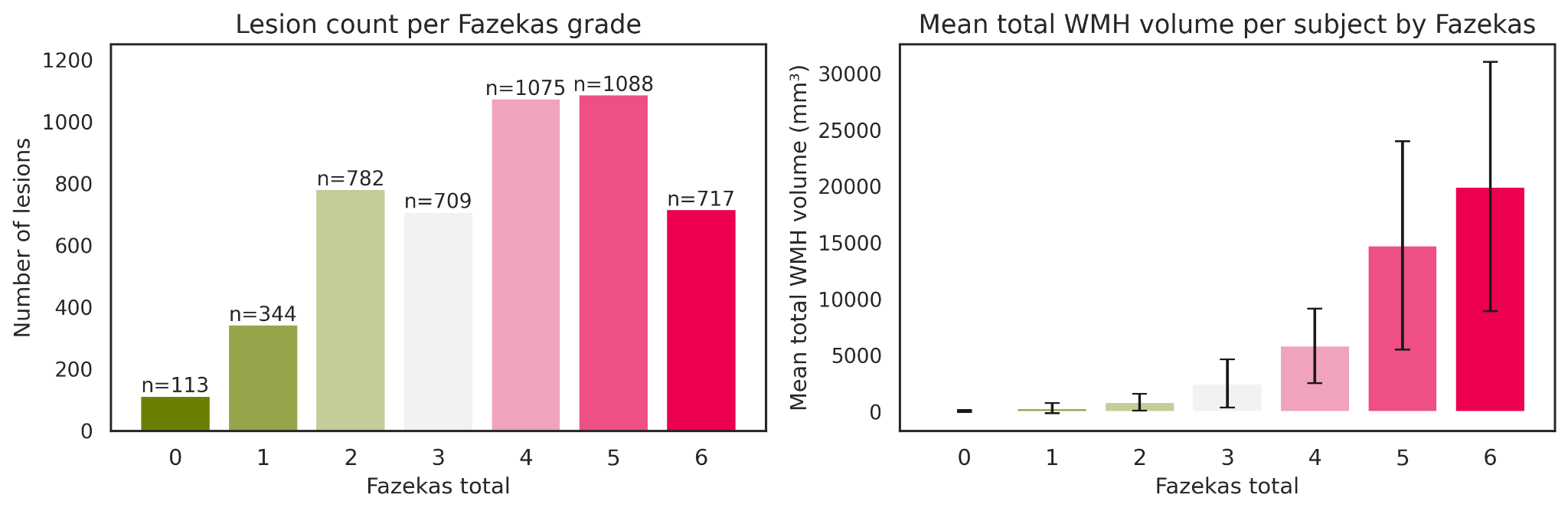

**Supplementary Figure 23:** A) Total number of white matter hyperintensity (WMH) lesions across subjects for each Fazekas total score. B) Mean total WMH volume per subject (mm³) for each Fazekas total score (error bars represent standard deviation).

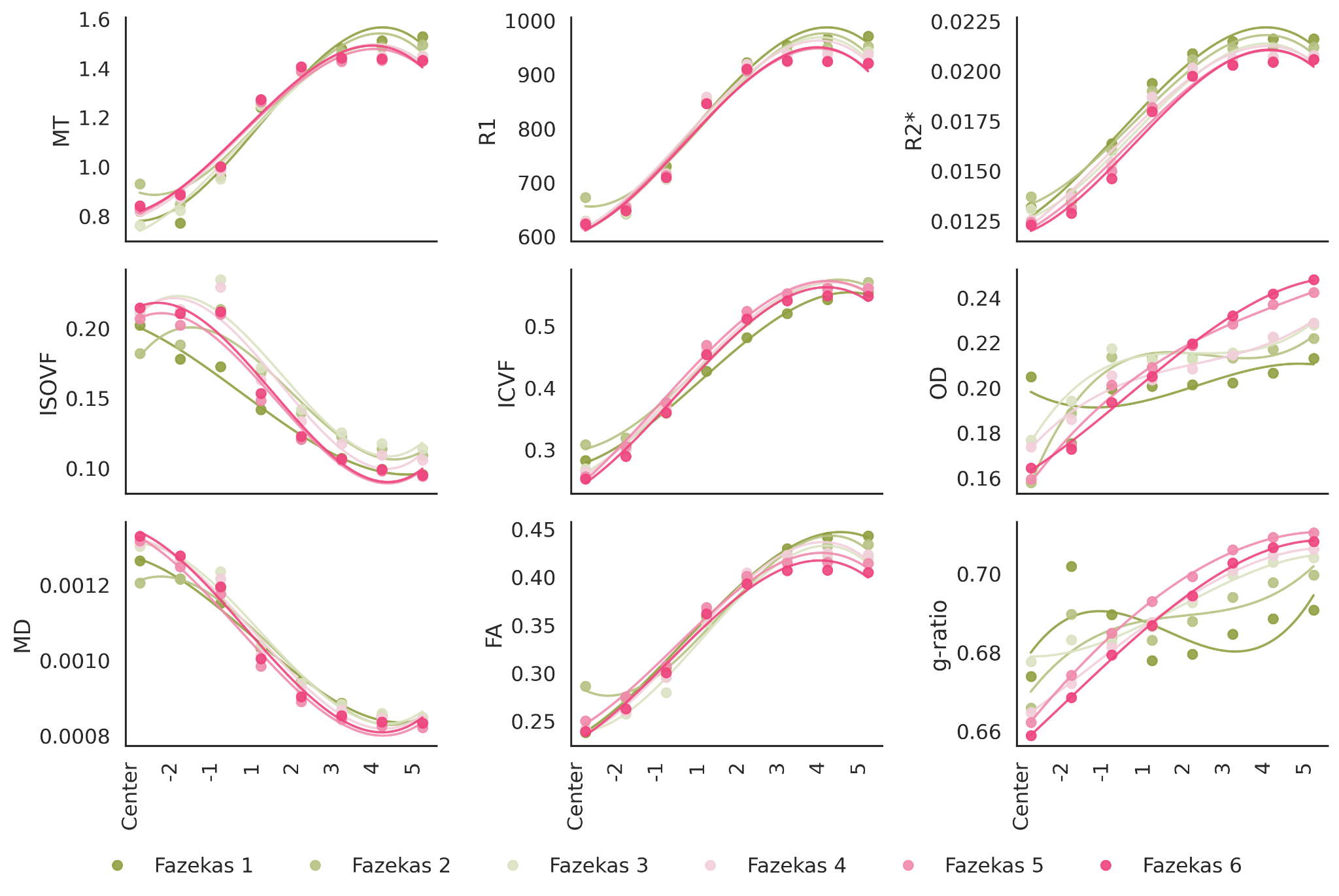

**Supplementary Figure 24**. Layer-based tissue microstructural profiles stratified by Fazekas score, using the geodesic total-WMH distance method in the full sample. Participants with Fazekas 0 were excluded, yielding n = 257 individuals. For each microstructural metric, cubic polynomial fits (colored lines) illustrate how values change from the WMH core outward into the surrounding NAWM across Fazekas groups, highlighting progressive alteration in tissue microstructure as global WMH burden increases.

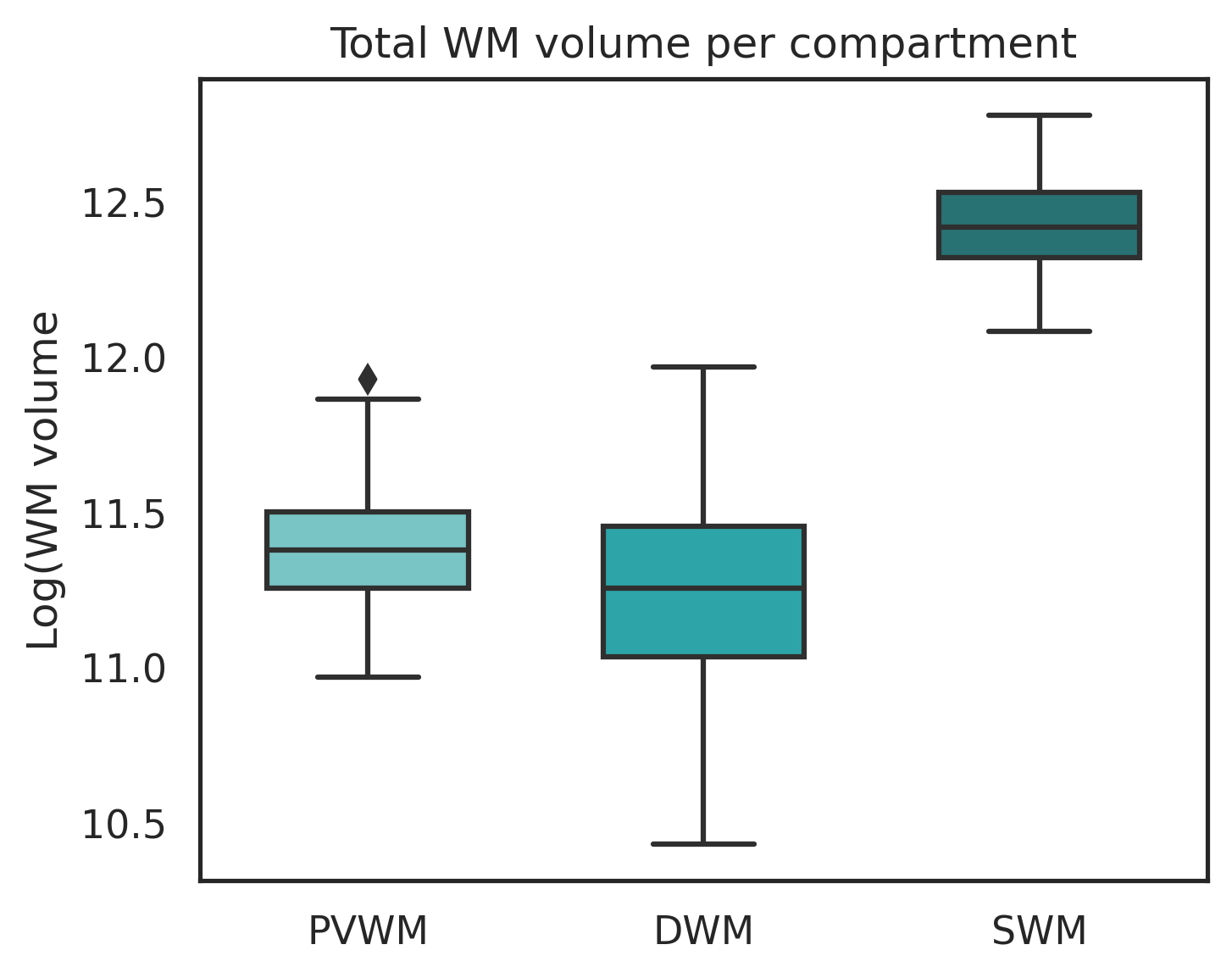

**Supplementary Figure 25:** Total white-matter volume of the PVWM, DWM, and SWM compartments. Boxplots show the distribution of log-transformed white-matter volume (in voxels) for the periventricular (PVWM), deep (DWM), and superficial (SWM) white-matter partitions across all participants.
