## Supplementary material for "White matter microstructure fingerprint of cerebral small vessel disease": Table 1

**Table 1.** Demographic and CVR characteristics (count, %, or mean ± SD) for the main sample (n=363) and the CVR sample (n=202) and for each Fazekas category. **Abbreviations:** systolic (SBP) and diastolic blood pressure (DBP), total cholesterol (Chol), Triglycerides (Trig), waist-to-hip ratio (WHR), body mass index (BMI), glycated hemoglobin (HbA1c), high-sensitivity C-reactive protein (CRP), hemoglobin, thyroid-stimulating hormone (TSH), free thyroxine (FT4).

|  | **Fazekas** | **All** | **0** | **1** | **2** | **3** | **4** | **5** | **6** |
| --- | --- | --- | --- | --- | --- | --- | --- | --- | --- |
| **Main sample**  **n=363** | **n PR** | 363 | 101 | 87 | 64 | 41 | 31 | 23 | 16 |
|  | **Age** | 55.5 ± 22.3 | 31.8 ± 12.5 | 48.4 ± 20.9 | 67.3 ± 11.0 | 74.7 ± 6.9 | 75.6 ± 6.3 | 75.7 ± 5.4 | 78.8 ± 6.4 |
|  | **Percent Male** | 48.8% | 51.5% | 57.5% | 48.4% | 36.6% | 51.6% | 39.1% | 25.0% |
| **CVR sample**  **n=202** | **n PR** | 202 | 9 | 35 | 53 | 38 | 30 | 21 | 16 |
|  | **Age** | 69.4 ± 6.9 | 63.7 ± 5.6 | 66.1 ± 5.7 | 66.7 ± 6.1 | 71.9 ± 6.3 | 71.8 ± 6.9 | 71.3 ± 5.6 | 75.3 ± 6.6 |
|  | **Percent Male** | 46.5% | 55.6% | 65.7% | 49.1% | 34.2% | 46.7% | 38.1% | 31.2% |
|  | **SBP** | 130.9 ± 19.4 | 127.6 ± 18.7 | 128.4 ± 17.3 | 125.5 ± 17.6 | 130.1 ± 19.3 | 137.2 ± 17.2 | 134.1 ± 19.5 | 141.8 ± 27.1 |
|  | **DBP** | 76.6 ± 10.2 | 76.9 ± 10.2 | 76.2 ± 10.0 | 75.4 ± 9.1 | 76.9 ± 9.1 | 77.5 ± 10.3 | 77.5 ± 13.5 | 77.1 ± 12.8 |
|  | **Chol** | 5.1 ± 1.0 | 5.1 ± 1.2 | 5.1 ± 1.0 | 5.2 ± 1.0 | 5.2 ± 1.1 | 4.8 ± 0.8 | 5.5 ± 1.2 | 5.3 ± 1.0 |
|  | **LDL** | 3.0 ± 0.9 | 3.0 ± 1.2 | 3.0 ± 0.9 | 3.0 ± 0.9 | 2.9 ± 0.9 | 2.8 ± 0.8 | 3.4 ± 1.0 | 3.2 ± 0.9 |
|  | **HDL** | 1.6 ± 0.4 | 1.5 ± 0.4 | 1.4 ± 0.5 | 1.6 ± 0.4 | 1.7 ± 0.4 | 1.5 ± 0.4 | 1.6 ± 0.3 | 1.6 ± 0.3 |
|  | **Trig** | 1.3 ± 1.1 | 1.1 ± 0.5 | 1.6 ± 2.4 | 1.3 ± 0.6 | 1.2 ± 0.6 | 1.1 ± 0.5 | 1.2 ± 0.4 | 1.2 ± 0.5 |
|  | **Glucose** | 5.5 ± 0.8 | 5.3 ± 0.3 | 5.5 ± 0.9 | 5.5 ± 0.8 | 5.5 ± 0.9 | 5.8 ± 1.0 | 5.5 ± 0.5 | 5.0 ± 0.4 |
|  | **Insulin** | 9.7 ± 5.6 | 10.7 ± 2.8 | 8.5 ± 4.5 | 9.5 ± 6.3 | 9.9 ± 5.9 | 11.4 ± 7.2 | 10.5 ± 4.1 | 7.3 ± 3.3 |
|  | **WHR** | 0.9 ± 0.1 | 0.9 ± 0.1 | 0.9 ± 0.1 | 0.9 ± 0.1 | 0.9 ± 0.1 | 0.9 ± 0.1 | 0.9 ± 0.1 | 0.9 ± 0.1 |
|  | **BMI** | 25.5 ± 3.9 | 26.5 ± 3.0 | 25.8 ± 4.5 | 25.5 ± 4.3 | 24.9 ± 3.5 | 25.9 ± 3.7 | 26.5 ± 4.3 | 24.2 ± 2.5 |
|  | **Bioimpedance** | 34.4 ± 8.6 | 34.9 ± 7.3 | 30.8 ± 9.3 | 33.4 ± 8.1 | 35.2 ± 8.5 | 35.3 ± 7.8 | 37.3 ± 7.7 | 37.5 ± 9.9 |
|  | **Alcohol** | 5.6 ± 6.4 | 9.7 ± 8.9 | 5.5 ± 6.5 | 5.5 ± 6.0 | 4.4 ± 5.2 | 5.1 ± 6.3 | 5.0 ± 4.4 | 8.1 ± 9.5 |
|  | **Tobacco** | 1.6 ± 4.7 | 0.4 ± 1.3 | 1.8 ± 5.9 | 2.5 ± 5.9 | 0.4 ± 1.5 | 1.6 ± 4.9 | 1.2 ± 3.1 | 2.0 ± 5.2 |
|  | **Hba1C** | 37.6 ± 5.5 | 36.8 ± 3.5 | 37.5 ± 5.1 | 37.4 ± 7.1 | 37.8 ± 4.2 | 39.6 ± 6.6 | 36.5 ± 3.3 | 36.1 ± 3.0 |
|  | **TSH** | 2.3 ± 1.3 | 1.8 ± 1.2 | 2.4 ± 1.5 | 2.3 ± 1.2 | 2.6 ± 1.5 | 2.1 ± 1.0 | 2.2 ± 1.4 | 2.2 ± 1.2 |
|  | **FT4** | 15.2 ± 2.1 | 15.5 ± 2.3 | 14.7 ± 1.8 | 15.3 ± 2.3 | 15.7 ± 2.4 | 15.4 ± 1.7 | 15.2 ± 2.2 | 14.5 ± 2.0 |
|  | **Hemoglobin** | 140.6 ± 11.2 | 142.2 ± 17.8 | 142.4 ± 12.6 | 141.2 ± 9.8 | 138.2 ± 10.4 | 142.0 ± 12.2 | 140.2 ± 10.3 | 137.7 ± 8.8 |
|  | **CRP** | 2.2 ± 2.6 | 1.8 ± 1.4 | 1.7 ± 1.4 | 2.0 ± 2.0 | 2.1 ± 3.1 | 2.8 ± 3.4 | 2.0 ± 1.8 | 2.9 ± 4.6 |
